# Nanoparticle-Supported, Rapid, and Electronic Detection of SARS-CoV-2 Antibodies and Antigens at Sub-Femtomolar Level

**DOI:** 10.1101/2024.09.04.611305

**Authors:** Yeji Choi, Seyedsina Mirjalili, MD Ashif Ikbal, Sean McClure, Maziyar Kalateh Mohammadi, Scott Clemens, Jose Solano, John Heggland, Tingting Zhang, Jiawei Zuo, Chao Wang

**Author notes:** Corresponding Author: Chao Wang - School of Electrical, Computer and Energy Engineering, Arizona State University, Tempe, AZ 85287, USA; Biodesign Center for Molecular Design and Biomimetics, Arizona State University, Tempe, AZ 85287, USA. These authors contributed equally (Yeji Choi, Seyedsina Mirjalili).

## Abstract

Major challenges remain to precisely detect low-abundance proteins from diverse biofluids in a rapid and cost-effective manner. Here we present a gold nanoparticle (AuNP)-supported, rapid electronic detection (NasRED) platform with sub-femtomolar sensitivity and high specificity. Surface-functionalized AuNPs act as multivalent detectors to recognize target antigens and antibodies through high-affinity binding, subsequently forming aggregates precipitated in a microcentrifuge tube and producing a solution color change. The optical extinction of residual floating AuNPs is digitized using a customized circuitry incorporating inexpensive optoelectronic elements and feedback mechanisms for stabilized readout. Uniquely, NasRED introduces active fluidic forces through engineered centrifugation and vortex agitation, effectively promoting protein detection at low concentrations and accelerating signal generation. Using SARS-CoV-2 as a demonstration, NasRED enables detection of both antibodies and antigens from a small sample volume (6 µL), distinguishes the viral antigens from those of human coronaviruses, and delivers test results in a short time (as fast as <15 min). The limits of detection (LoDs) for antibody detection are approximately 49 aM (7 fg/mL) in phosphate-buffered saline, or >3,000 times more sensitive than Enzyme-Linked Immunosorbent Assay (ELISA), ∼76 aM (11 fg/mL) in human pooled serum and in the femtomolar range in diluted whole blood. For nucleocapsid protein detection, NasRED LoDs are ∼190 aM (10 fg/mL) in human saliva and ∼2 fM (100 fg/mL) in nasal fluid. Unlike laboratory-based ELISA platforms, NasRED is a one-pot, in-solution assay that eliminates the needs for washing, labeling, expensive instrumentation or highly trained operators. With low reagent costs and a compact system footprint, this modular digital platform is well-suited for accurate, near-patient diagnosis and screening of a wide range of infectious and chronic diseases.

## INTRODUCTION

Infectious diseases pose a serious threat to human health and the world economy. For example, as of March 1^st^, 2025, COVID-19 alone has resulted in more than 777 million infections and 7 million deaths^1^, while causing an estimated $14 trillion in economic losses in the U.S. by the end of 2023^2^. In addition, the pandemic also seriously strained the global healthcare system^3,4^ and caused delayed or missed diagnosis or treatment for many other patients. To better prepare for future pandemics, which could be caused by a currently unknown disease ‘X’^5,6^, it is of paramount importance to continue investing in innovative diagnostic technologies. These technologies should enable early and accurate disease detection and quantification in a portable, cost-effective, and user-friendly manner, ensuring accessibility for all patients in need, including those in rural areas and low- to middle-income countries where medical resources are scarce. To complement this, various new approaches, including label-free high-sensitivity diagnostic technologies, have been proposed recently^7,8^.

Conventionally, molecular and immunochemical tests are widely used to diagnose infectious diseases, including COVID-19^9,10^. These tests look for molecules of different genetic information to diagnose diseases and predict the risk of unknown diseases^11^. Nucleic Acid Amplification Tests (NAATs), including Polymerase Chain Reaction (PCR) and Loop-mediated Isothermal Amplification (LAMP), leverage the feasibility of DNA or RNA amplification to maximize the sensitivity. However, as the gold standard for laboratory diagnostics, PCR requires several hours to days to amplify and confirm the results due to the multiple steps of sample handling and the thermal cycling process^12^. Additionally, it requires expensive, bulky equipment, making it suitable for centralized testing but inaccessible in resource-limited settings. Isothermal amplification methods, such as LAMP, while offering a simpler and more cost-effective alternative to PCR, face challenges in practical applications, including the complexity in primer design, the limited stability of enzymatic reagents, and the need of careful sample preparation when working with complex biological fluids such as blood and saliva.

Immunochemical tests diagnose diseases by confirming the presence or absence of a specific protein based on antigen-antibody reactions. The most commonly used methods include Enzyme-Linked Immunosorbent Assay (ELISA) and Lateral Flow Assay (LFA)^13^. ELISA typically achieves picomolar-level sensitivity in biological fluids ^14–16^. However, like PCR, ELISA is most suited for high-sensitivity analysis of large amounts of samples in laboratory settings since it requires a complex, multistep workflow, making it highly labor-intensive and expensive^17,18^. In addition, Rapid Antigen Tests (RATs), such as LFA, are portable and suitable for field applications but generally have lower sensitivity (e.g., 65.3% for COVID-19). This limitation makes them less reliable for accurate detection, particularly at early-stage infection when the viral load is low^19–21^. Here, using SARS-CoV-2 as an example, we report a prototyped, inexpensive, miniaturized sensing system, i.e. nanoparticle-supported rapid electronic detection (NasRED), to detect the antibodies and antigens directly from serum, whole blood, saliva, and nasal fluid within a short time (15 to 30 min, comparable to LFA). NasRED achieves femtomolar or better sensitivity (fg/mL), about 2 to 4 orders of magnitude better than ELISA^22^ and about 5 to 6 orders better than LFA^23^, enabling rapid and accurate diagnostics of disease biomarkers in different clinical settings. Specifically, our digital sensing platform was designed to detect the target proteins by simply mixing bodily fluids with our prepared sensing solution comprising functionalized gold nanoparticles (AuNPs) in a testing tube ^24,25^. The entire process, including mixing, reaction, and signal readout, occurs within the same tube without washing, labeling, or fluorescent imaging, significantly reducing operation complexity. The sensing signals were collected, digitized, and transmitted through our customized semiconductor devices and circuits, which minimized signal fluctuations and enable the accurate detection of antigens and antibodies at ultralow concentrations. Further, we comprehensively engineered the sensing protocol, i.e. the centrifugation and vortex agitation speed and time, to evaluate their impact on the limits of detection (LoDs) of SARS-CoV-2 antibody (AS35) and antigen (N-protein). The LoDs were found <100 fg/mL from a wide range of bodily fluids, including in diluted whole blood without requiring plasma separation. This high sensitivity is attributed to the application of active fluidic forces during centrifugation and vortex agitation, which enhance protein collisions, biochemical binding, and signal transduction, even at low concentrations. Additionally, vortex agitation creates an AuNP concentration gradient along the liquid column, thus enhancing optical signals and enabling detection of low-abundance proteins. Lastly, using inactivated SARS-CoV-2 virion particles, NasRED successfully detected N-protein from lysate products in saliva with high sensitivity (190 aM or 10 fg/mL), outperforming ELISA. Based on parallel qRT-PCR measurements, this protein detection level corresponds to an estimated RNA sensitivity of 3×10^5^ copies/mL, demonstrating performance comparable to portable nucleic acid amplification tests. Together, our preliminary data demonstrates the strong promise of NasRED as a new and broadly applicable platform for highly sensitive, rapid, and accessible detection of infectious diseases as well as chronic diseases.

## RESULTS

### NasRED sensing platform

The NasRED protein sensing platform can be applied to the detection of both the antigens and the antibodies (Figure 1). The sensors are prepared by attaching biotinylated proteins to streptavidin-coated AuNPs through a biotin-streptavidin reaction, where the protein binders can be antigens (e.g. SARS-CoV-2 spike (S) or receptor binding domain (RBD) proteins) for antibody (neutralizing antibody AS35) sensing or antibodies (mouse IgG1 AS47 and chimeric IgG1 AM223) for antigen (nucleocapsid (N) protein) sensing (Figure 1a). Using antibody sensing as an example (Figure 1b), 18 µL of the RBD-functionalized AuNPs were mixed with 6 µL of medium (e.g., PBS buffer, human pooled serum (HPS), or human whole blood (WB)) containing target SARS-CoV-2 antibodies diluted to desired concentrations. A sample of blank medium without antibodies was used as a negative control (NC). For rapid detection, the 24 µL mixture was centrifuged at 1,200 g for 5 minutes and incubated for 5 minutes to precipitate the functionalized AuNPs to the bottom of the microcentrifuge tube, where their concentration was greatly enhanced. Such a localized and boosted reactant concentration significantly promoted the localized reaction with target proteins, thus accelerating the antibody-antigen reaction at low concentrations to achieve an ideal LoD and reduce assay time. To preserve the detection specificity, a vortex agitation was introduced to disperse unbound AuNPs without disturbing the precipitated AuNP clusters, verified by fully dispersed AuNPs in the NC reference (Figure 1c). The optical absorption signal from the supernatant, attributed to free-floating AuNPs after vortexing, was digitized using a portable electronic detector (PED) for quantification (Figure 1d-e). The electronic signals of the antigen or antibody samples collected from the PED represented the optical transmission of light-emitting diode (LED) light through the supernatant in the microcentrifuge tube and were normalized (Methods). The NasRED system has proven to be feasible to detect antigens and antibodies in complex biological fluids with high sensitivity and specificity (Figure 1f-g).

**Figure 1.**
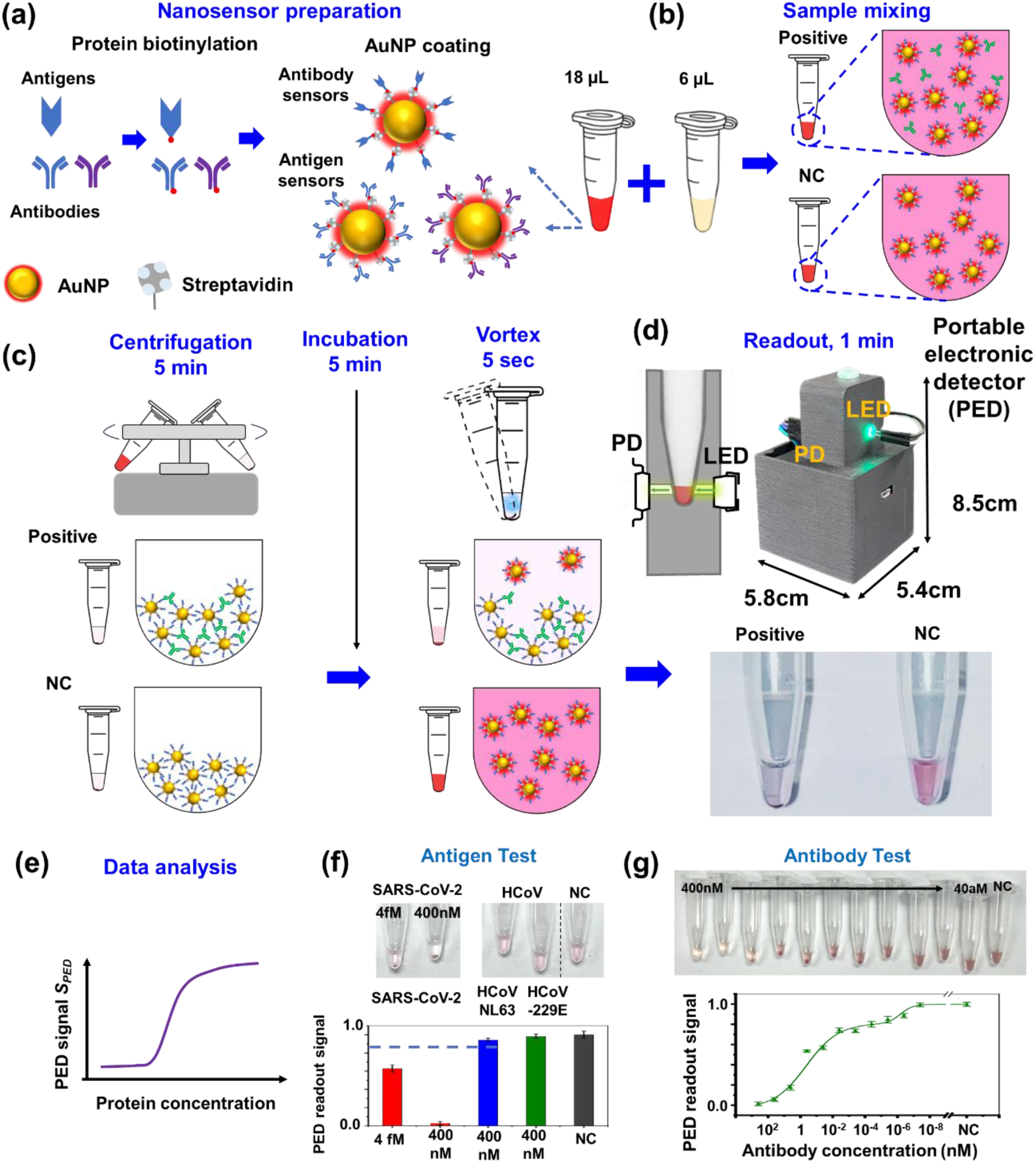
Nanoparticle-supported, rapid electronic detection (NasRED) of antigens and antibodies for infectious diseases diagnostics. (a) Nanosensor preparation by biotinylating antigens or antibodies and attaching such proteins to streptavidin-coated AuNPs. (b-d) Schematics and optical images illustrating the detection process, using antibody testing as an example. (b) The AuNP sensing solution (18 µL) was mixed with the medium solution (6 µL). Negative controls (NCs) did not contain any target molecules but otherwise contained the same biological fluids and AuNP sensors. (c) The AuNPs were precipitated by centrifugation, incubated, and resuspended after stirring vortex. (d) The readout was collected directly from the tube, which was inserted in a tube holder as part of the NasRED readout system. The amount of AuNP precipitation at the tube bottom correlated with the number of test molecules, while no AuNP precipitates formed in the NC sample. (e) Schematic of a typical sensing curve showing the normalized PED readout signal *S_PED_*. (f) Exemplary optical images and bar plots showing sensitivity and specificity in SARS-CoV-2 antigen detection in PBS. Tubes containing N-proteins of two other coronaviruses (HCoV-NL63 and HCoV-229E at 400 nM in PBS buffer) displayed very similar colors and electronic signals comparable to that of the NC sample. In contrast, tubes containing SARS-CoV-2 N-proteins showed much higher transparency and differentiable electronic signals at 4 fM and 400 nM. (g) Exemplary optical images and sensing curve plots showing antibody detection at high sensitivity in diluted whole blood.

### Signal-stabilizing circuit for low-noise detection

The NasRED platform consists of a portable electronic detector (PED) for signal readout, which is composed of two customized components (Figure S1a). The first component is a semiconductor-based, digital, signal-collecting device, which includes an LED, a photodetector (PD), and a microcentrifuge tube chamber designed to block ambient light. This device functions to probe the free-floating AuNP concentration in the supernatant of the testing tube and to digitize the optical signals of the AuNPs to electronic current signals. The second component is a circuit board designed to stabilize the LED and PD signals for reliable antibody or antigen detection. Electronically, the stabilizing circuit board (Figure 2a-b) had three key components: a voltage regulator that adjusts the power supply to the desired value for the LED circuit and microprocessor, a constant current LED driver circuit that stabilizes the light intensity, and a microprocessor for signal processing. The regulator circuit includes two electrolytic capacitors, two ceramic capacitors, and one protective Zener diode (Figure 2a (i)). The constant current LED driver circuit utilizes a feedback loop that coordinates double bipolar junction transistors (BJTs, T1 and T2 in Figure 2a (ii)) to stabilize the current supplied to the LED. In addition, a customized microprocessor (Figure 2a (iii)) processes the signals and enables communication with a laptop through USB, Wi-Fi, or Bluetooth modules. Without the stabilization circuit, the LED displayed a large deviation in current (average 0.14 mA over 10 minutes but significantly increased after the 40-min operation, possibly due to heating, Figure 2c), and the PED produced random spike noises when used to test AS35 antibodies in PBS (Figure 2d). This instability can be attributed to the fact the LED light emission is proportional to its electrical current and exponentially depends on the bias voltage, and therefore, a slight voltage fluctuation from a power glitch or device heating can introduce a systematic error, ultimately seriously limiting the sensing performance at low analyte concentrations. In comparison, our improved readout system demonstrated significantly reduced current fluctuation (0.12 mA within the first two minutes during circuit initiation and decreased to 0.02 mA during the rest 60-min testing period, Figure 2c) and greatly minimized system error in antibody detection (Figure 2d), indicating a substantial improvement in stability.

**Figure 2.**
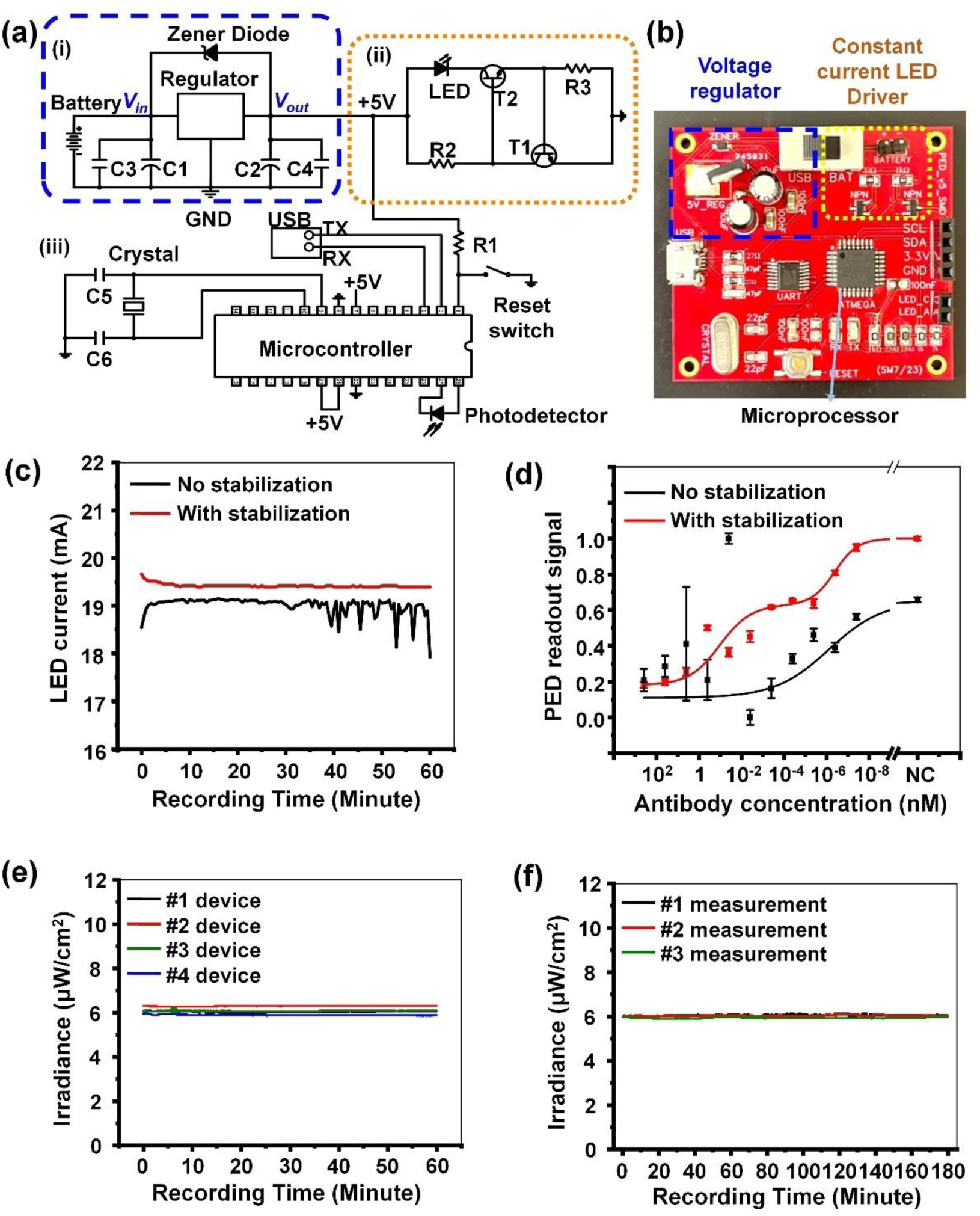
Electronic circuit design for stabilized NasRED readout. (a) Schematic showing an integrated printed circuit board (PCB), which includes: (i) a built-in voltage regulator, (ii) a constant current LED driver circuit, and (iii) a microcontroller, which processes the signals and communicates with computers through USB, Wi-Fi, or Bluetooth modules. (b) Optical image showing the PCB board for signal processing. (c) The stability of LED currents over time for a circuit without the stabilization circuit (black line) and our integrated PCB board with the stabilization circuit (red line). (d) PED readout signal *S_PED_* plotted against antibody AS35 concentrations spiked in PBS, with (red) and without (black) the stabilization circuit. (e) LED irradiance recorded for 60 minutes using four different devices #1 (black line), #2 (red line), #3 (green line), and #4 (blue line). (f) Three, three-hour-long, replicate measurements of LED irradiance from one PED reader.

To further evaluate the reproducibility and reliability of the PED circuit design, four different PED devices were constructed (Figure S2). The LED output power from each PED circuit was measured by a PD and found stable at ∼6 μW/cm^2^ over a 60-minute test period (Figure 2e) with a very small (∼2.5%) coefficient of variation (CV). Then, one PED circuit was randomly selected, and its LED power was measured three times, 3 hours each (Figure 2f). The measured irradiance was again very stable at ∼6 μW/cm^2^ with a CV of 2.4%. These inter- and intra-device tests proved the high reproducibility and suitability of our NasRED reading system for continuous use or repeated operation.

### Impacts of centrifugation and vortex agitation on sensing performance

The centrifugation and vortex agitation actively modulate the reagent and AuNP distribution in the buffer solutions, providing additional flexibility to optimize the sensitivity and specificity. To examine their impact on the NasRED assay performance, we chose mAb AS35 (6 µL, from 4 µM to 40 fM in PBS) as the target antibody, using AuNPs functionalized with WT-RBD (Wuhan-Hu-1, 18 µL in1×PBS dilution buffer). Electronic signals were collected using our stabilized readout system. Here, a series of tests (Figure 3a-d) were performed by modifying four variables: the centrifugate force gravity (g) (Figure 3a), centrifugation time (Figure 3b), vortex agitation speed (Figure 3c), and the vortex time (Figure 3d). The normalized assay signals, *S*_*PED*_*(c_p_)*, were plotted as a function of AS35 concentration *c_p_* (Figure 3a-d). The default parameters were centrifugation at 1,200 g for 5 minutes, incubation for 5 minutes, and vortex agitation at 34.5 rps for 5 seconds unless otherwise varied. It was clear that the AS35 antibodies could be detected with a large contrast in signal *S*_*PED*_, i.e. a high value at low concentrations (*S_H_* = *S*_*PED*_(*NC*) ≅1) and a small signal at high concentrations (*S*_L_ = *S*_*PED*_(*c_H_*) ≅0.1, *c_H_* = 4µM), or Δ*S*_*PED*_ = *Sc_H_* − *S*_L_ ≅ 0.9. A high signal contrast Δ*S*_*PED*_ indicated that the AuNP sensors effectively participated in the biochemical reaction at high protein concentrations and subsequently precipitated, which was ideal for maximizing the sensitivity and dynamic range (about 9 logs in this test). Clearly, the sensing performance of NasRED could be optimized within a range of centrifugation conditions, i.e., from 900 g to 1500 g and from 3 to 5 minutes, and at moderate vortex conditions.

**Figure 3.**
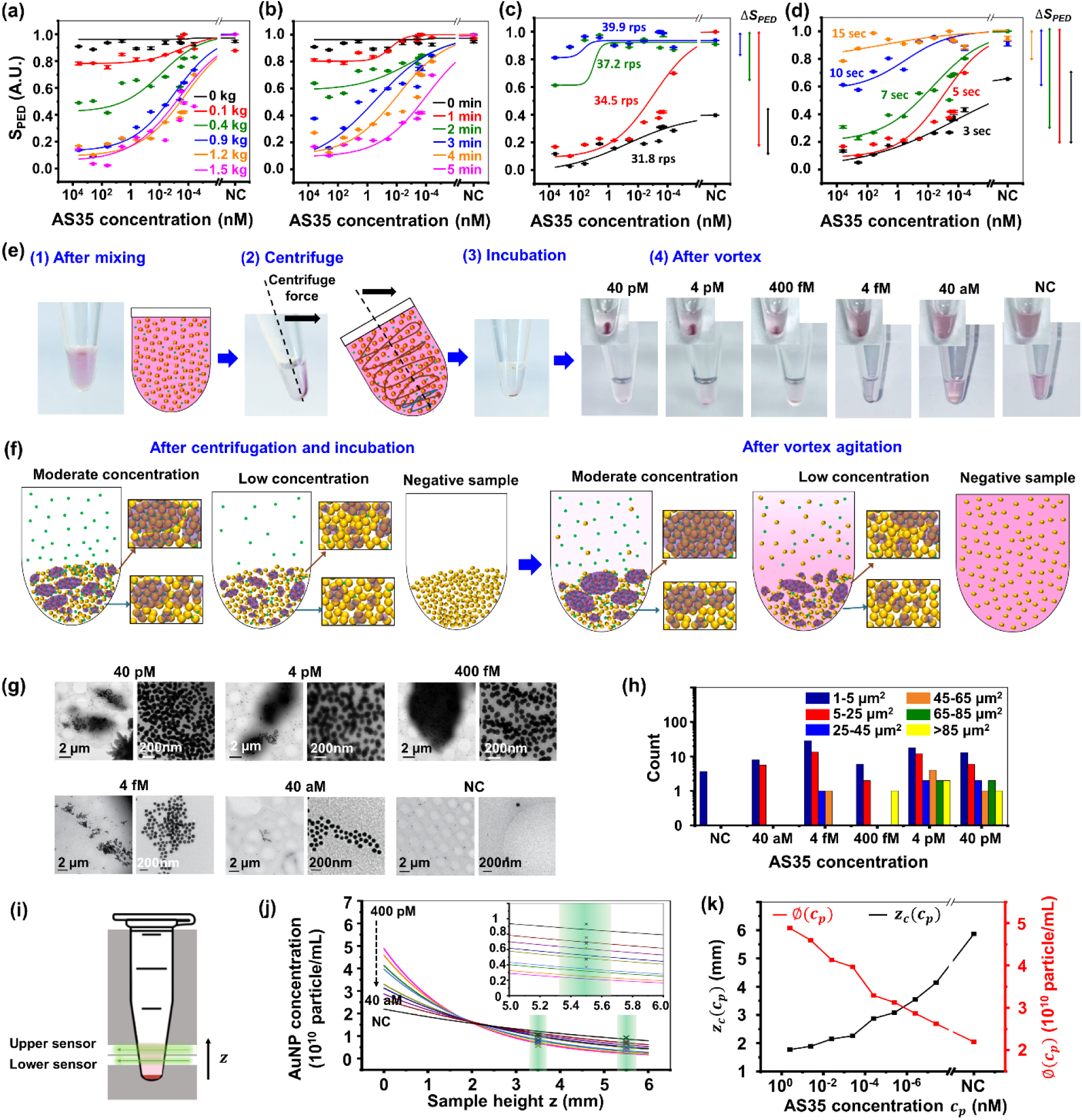
Optimization of NasRED sensing protocol under different centrifugation and vortex conditions. (a-d) The PED readout signal *S*_*PED*_ plotted as a function of the antibody concentration, with: (a) the centrifugate force from 0 to 1,500 gravity (g), (b) the centrifugation time from 0 to 5 minutes, (c) the vortex agitation speed from 32 to 40 round per second (rps), and (d) the vortex time from 3 to 15 seconds. (e-f) Schematic representations of the AuNP cluster formation and precipitation mechanism during the sensing process: (1) uniform mixing of AuNPs with target proteins, (2) AuNP sensors react to and bind to target proteins, thus forming clusters, during centrifugation, (3) after centrifugation and during incubation, the AuNP clusters form larger precipitates at the tube bottom, and (4) vortex agitation releasing loose AuNPs back into the solution. (g) Representative Cryo-TEM images of 80 nm AuNP clusters at AS35 concentrations of 40 pM, 4 pM, 400 fM, 40 fM, 40 aM, and NC. (h) Statistical analysis of AuNP cluster sizes formed at different AS35 concentrations. NC: negative control using blank sample. (i–k) Modeling and analysis of vertical AuNP concentration distribution. (i) Schematic of the optical detection setup with dual-position PED readout, with green arrows indicating upper (5.5 mm) and lower (3.5 mm) sensing locations and black arrow indicating the z direction in the mathematical model (z=0 at the tube bottom). (j) Reconstructed AuNP concentration profiles using an exponential decay model in the z direction, fitted to the measured concentrations at both PED readout positions (highlighted by green). The profiles were generated using Python under a mass conservation constraint. (k) Extracted model parameters ∅*(c_p_)* and Z_c_*(c_p_)* as a function of analyte concentration. ∅*(c_p_)* refers to the AuNP monomer concentrations at z=0 and Z_c_*(c_p_)* is the characteristic decay length of AuNP concentrations in the model.

### NasRED sensing mechanism

The dependence of NasRED’s performance on the centrifugation and vortex parameters can be understood from the unique sensing mechanism. First of all, fundamentally, NasRED is an in-solution assay, where diffusion of AuNP sensors and reagents drives signal transduction, leading to AuNP precipitation and ultimately determining the assay performance. In NasRED, the sedimentation time of AuNPs linearly scales with the precipitation length^26^. Unlike conventional assays such as ELISA, which rely on the passive diffusion of target molecules to a stationary reaction surface and require long incubation time to maximize the signal-to-noise ratio, NasRED supports external controls, such as centrifugation, incubation, and vortex agitation, to actively accelerate diffusion, reaction, and readout (Figures 3 and S3). Centrifugation pellets AuNPs dispersed in the tube to the bottom, with a thickness estimated as <0.5 mm (supplementary section 4), thus eliminating slow sedimentation and accordingly reducing the readout time to less than 30 minutes (Figure 3a-b). In comparison, passive precipitation of AuNP clusters was previously found to require 3 to 24 hours of assay time from mixing to readout^24^ (Figure S4a-c). Additionally, the PED signal was found stable for hours after vortex agitation (Figure S4d-e), indicating the establishment of a quasi-equilibrium state desirable for reliable readout.

Further, the introduction of active fluidic forces is an integral part of NasRED’s unique signal transduction process, which begins with biochemical protein reaction and is dynamically modulated by fluidic interactions (Figure 3e-f). Centrifugation significantly increases the kinetic velocity of AuNPs, enhancing their collision with and effective capture of the target proteins, which in turn promotes the growth of AuNP clusters. Protein binding and AuNP cluster growth are expected to occur throughout the centrifugation process, ultimately leading to precipitation of the majority of the AuNPs (Figure 3e). The dimensionless Peclet number (Pe), which compares nanoparticle centrifugal sedimentation to Brownian motion, scales linearly with the relative centrifugal force (RCF) and quantically with the AuNP diameter (supplementary section 4). For example, the Pe number was found ∼1.92 for 80 nm AuNPs at an RCF of 1200×g, while significantly lower values were obtained for smaller nanoparticles (∼0.01, 0.12, and 0.61 for 20 nm, 40 nm, and 60 nm AuNPs, respectively). The estimates suggest that the centrifugal forces dominate over Brownian motion during the sedimentation process of 80 nm and larger AuNPs, aligning with our prior observation that 80 nm AuNPs precipitated faster than smaller ones^24^. On the other hand, the particle Reynolds number (*Re*_*p*_), which quantifies the ratio of inertial to viscous forces, increases proportionally with the AuNP diameter. *Re*_*p*_ was estimated ∼0.04 for 80 nm AuNPs, suggesting that fluidic drag forces remain a significant factor in governing the particle motion and dispersion within biological fluids. Consequently, the NasRED sensing protocol should be individually optimized for fluid media with different viscosities.

The AuNPs are expected to capture target proteins while following distinct sedimentation paths, eventually forming precipitates composed of heterogeneous clusters, whose size distributions statistically correlate with protein concentrations (Figure 3f left). Importantly, centrifugation greatly localizes AuNPs and their clusters into a much smaller volume (estimated <1 nL, or 3 to 4 orders more condensed, supplementary section 4) and drastically boost the AuNP concentration at the bottom of the reaction tube. This effectively modulates the dynamic equilibrium of antigen-antibody reaction to favor AuNP cluster formation both during centrifugation and incubation, even at ultralow reagent concentration in the tube^24^. During incubation, AuNPs deposited on the tube sidewalls gradually settle to the tube bottom, thus minimizing the nonspecific particle interference with the LED-PD signal-collecting optical path in the upper solution. Additionally, AuNPs may possibly rearrange during incubation, forming more stable clusters that enhance sensitivity.

After incubation, sufficient vortex agitation is required to fully resuspend monomer AuNPs that have not incorporated into clusters, ensuring high specificity, i.e., the negative control should return to their initial states. However, excessively high agitation risks breaking the AuNP clusters biochemically “glued” together, even at moderate or high protein concentrations. This would diminish the signal contrast Δ*S*_*PED*_ (Figure 3c-d arrows on the right) and negatively affect the NasRED sensitivity. Therefore, optimizing the agitation force and duration (e.g. 5 sec at 34.5 rps for the chosen centrifugation conditions) is necessary to maximize the signal contrast Δ*S*_*PED*_ and to best differentiate samples at low concentrations, for example from attomolar to picomolar for AS35 (Figure 3e optical images). Interestingly, the 40 aM and 4 pM samples could be clearly differentiated from their precipitate formation in the tubes, indicating feasible attomolar detection. For more precise analysis, the AuNP precipitates were collected from the bottom of the tubes and imaged by Cryogenic Electron Microscopy (Cryo-EM) (Figure 3g and Figure S5). Then, the AuNP clusters at each AS35 concentration were measured and analyzed (Figure 3h). Evidently, large clusters (e.g. >50 µm^2^, up to ∼800 µm^2^) formed at moderate AS35 concentrations (e.g., 40 pM, 4 pM, and 400 fM) consisted of densely packed and stacked AuNPs. In comparison, smaller clusters were found at low antibody concentrations (<∼50 µm^2^ at 4 fM and <∼25 µm^2^ 40 aM), consisting of loosely connected AuNPs. Notably, AuNP clusters of >5 µm^2^ were only formed in the presence of AS35, whereas very small (<2 µm^2^) AuNP clusters or oligomers were detected from the NC (blank) sample, likely resulting from minor nonspecific inter-AuNP interactions during sample preparation (such as pipetting, TEM grid loading, and freezing). These Cryo-EM imaging provided direct physical evidence to confirm that the NasRED sensing protocol effectively dispersed the AuNPs for blank sample while differentiating antibodies at attomolar levels from higher-concentration and blank samples.

The fluidic dynamic behavior of AuNPs is complex in a turbulent flow during vortex agitation, potentially involving both shear- and spin-related fluidic drag forces^27^, and is strongly dependent on the *Re*_*p*_ value. Both shear- and spin-induced lift forces may diminish at higher *Re*_*p*_ values, and are highly dependent on the AuNP size, shape, flow velocity, rotation, and position within the fluid. As such, it is reasonable to expect that AuNP clusters, which exhibit a substantially higher effective *Re*_*p*_, experience reduced lift forces compared to AuNP monomers with smaller *Re*_*p*_. Therefore, AuNP clusters are far less likely to move upwards during vortex agitation. Precise physical modeling of these behaviors is challenging due to the complex fluidic dynamics, the protein-concentration-dependent, inhomogeneous mixture of AuNP monomers and clusters in the precipitates, and the three-dimensional conical geometry of the microcentrifuge tubes. Therefore, a detailed quantitative model will require extensive future studies and thus beyond the scope of this work.

To provide an intuitive understanding of the signal transduction mechanisms, we first established a correlation between the PED signal and AuNP concentration, by cross-validating results using the manufacturer-provided data, NanoSight calibration (Figure S6), and AuNP extinction calculations based on Beer–Lambert law (supplementary section 4). Then we built a PED device capable of measuring the AuNP extinction signals at two different fluidic levels, and converted the measured two-channel signals into the AuNP concentrations *C_AuNP_* at the two positions, for each tested AS35 antibody concentration *c_p_* (Figure 3i and Figure S7). We hypothesize that the AuNP concentration profile after vortex agitation follows a simplified exponential decay, consistent with the Mason-Weaver theory, and assume mass conservation of AuNPs. To test this hypothesis, we modeled the AuNP concentration distribution (Figure 3j) in Python as: 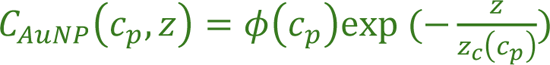. Here ø*(c_p_)* and *Z_p_(c_p_)* are empirical, analyte-concentration-dependent fitting variables (Figure 3k) that characterize the AuNP concentration at tube bottom (z=0) and characteristic decay length, respectively. The PED-measured and theoretically calculated *C_AuNP_* values showed strong overall agreement (<20% error, Figure 3j). This physics-based model supports our hypothesis that the AuNPs tend to aggregate more at the tube bottom and diffuse back less to the suspension at higher *c_p_*, also in agreement with our Cryo-EM analyses.

The optical tube images, PED readout signals, TEM analyses, and AuNP concentration modelling collectively demonstrated the success of attomolar detection from a small sample volume (6 µL). While further investigations are needed to fully elucidate the mechanisms behind the high-sensitivity of NasRED detection, a few factors possibly contribute. First, a strong antigen-antibody binding was achieved through multiple mechanism: (1) Multivalent binding on the AuNP sensor surfaces promotes high avidity, which has been demonstrated in pentameric IgM and nanobody oligomers to enhance the effective affinity by three to four orders of magnitude^24,28^. (2) Protein capture is enhanced during centrifugation, and their disassociation from AuNPs is suppressed. (3)

The reagent concentration at the tube bottom was enhanced about 4 orders of magnitude (supplementary section 4.3). Second, unlike fluorescent imaging that suffers from a high background noise, the PED readout has minimized background noise owing to successful strategic assay system design: (i) The stabilized PED readout device successfully suppresses signal fluctuations. (ii) NasRED collects the subtractive signals from a large population of AuNPs, thus minimizing stochastic background noises. (iii) Nonspecific interparticle interactions and tube wall adsorption are minimized by using inert AuNPs with covalently coated streptavidin as the signal carrier and by introducing glycerol and BSA in the AuNP sensor solution. (iv) A post-centrifugation incubation step allows the AuNP precipitates to settle at the tube bottom, thereby reducing unintentional interference to signal collection from free-floating AuNP monomers. Third, AuNP clusters bound by proteins may recruit non-protein-binding AuNP monomers through physical trapping, thus enlarging the AuNP precipitates and enhancing the PED signals. This could occur through disruption of the fluidic flow during centrifugation and/or vortex agitation, or by long-range attractive forces such as electrostatic interactions. TEM images (Figure S5) revealed inter-particle distance (e.g., >40 nm) far exceeding typical protein dimensions between AuNPs at the perimeters of clusters, suggesting possible long-range electrostatic interactions between protein analyte and protein-coated AuNPs^29^. Lastly, larger AuNP clusters (Figure 3h) not only better resist fluidic lift force than AuNP monomers during vortex agitation, but also influence the resuspension of smaller AuNPs. Consequently, the analyte concentration *(c_p_)*, which determines the size and amount of AuNP precipitates, ultimately plays an important role in the redistribution of AuNPs. The significant differences in the fitting parameters ø*(c_p_)* and Z_c_*(c_p_)* (Figure 3j) between sub-femtomolar analyte concentrations and blank samples resulted in a detectable change in AuNP concentration Δ*C_AuNP_* (about 4 × 10^9^). This change exceeded the minimum PED-resolvable concentration difference in *C_AuNP_* (3 × 10^8^), thus enabling sub-femtomolar protein detection on the NasRED assay^26^.

### NasRED comparison to ELISA

To benchmark the sensitivity of NasRED, the AS35 antibody was spiked in PBS at concentration range from 40 aM to 400 nM and analyzed using identical reagents on both standard ELISA assay (Figure 4a-b, Figure S8, and Table S2) and NasRED (Figures 4b and Figure S9) platforms. The LoDs, following two slightly different definitions, and LoQs were determined as described in the Method section and summarized in Table 1. For consistency, the LoDs reported in this work refer to the protein concentration *c_p_* at which the PED signal on the fitted sensing curve is distinguishable from the blank (NC) sample by three standard deviations (3σ) of the NC measurements, i.e. *S*_*PED*,*DS*_(*LoD*) = *S*_*PED*_(*NC*) − 3 × *SD*(*NC*). Here intra-assay imprecision (or within-test error, *SD*_*A*_) represents the systematic readout errors from the PED device at each *c_p_*. In contrast, inter-assay error (or between-test variation, *SD*_*I*_) captures the variability across replicate experiments and is used to assess assay reproducibility^30^. For NasRED assay LoD reporting, σ = *SD*_*I*_(*NC*) was determined from replicate blank samples (for example 20 samples in PBS). Remarkably, NasRED (LoD ∼49 aM, or 7 fg/mL, from six replicates) outperformed ELISA (LoD 155 fM, or 23.3 pg/mL, from three replicates) in sensitivity by about 3,000 times while requiring only 16 times less sample volume (6 µL compared to 100 µL) and 30 time shorter assay time (10 min compared to 5 hours). Additionally, the coefficient of variation (CV), an indicator of the variability of test results^31,32^, and recovery analysis, measurement of assay precision in quantification^32^, were also analyzed using for both NasRED and ELISA replicate tests. Our analyses showed repeated NasRED tests produced comparable LoDs (90 aM, 45 aM, 93 aM, 58 aM, 71 aM, and 60 aM, respectively, Figure S9a-b), determined by using the measurement error of the same NC sample along different tube orientations, or σ = *SD*_*A*_(*NC*). In addition, NasRED exhibited <10% CV values and moderate recovery rates mostly within a range of 100±30% (Figure S9c-d), suggesting consistent and reproducible detection with acceptable assay precision across the 10 logs of concentrations. In comparison, ELISA analyses exhibited significantly higher variation in absorbance intensity, CV, and recovery values, particularly at low to moderate antibody concentrations (attomolar to picomolar range) (Figure 4b-c). The comparison demonstrated that NasRED outperformed ELISA in both sensitivity and accuracy, particularly at low protein concentrations.

**Figure 4.**
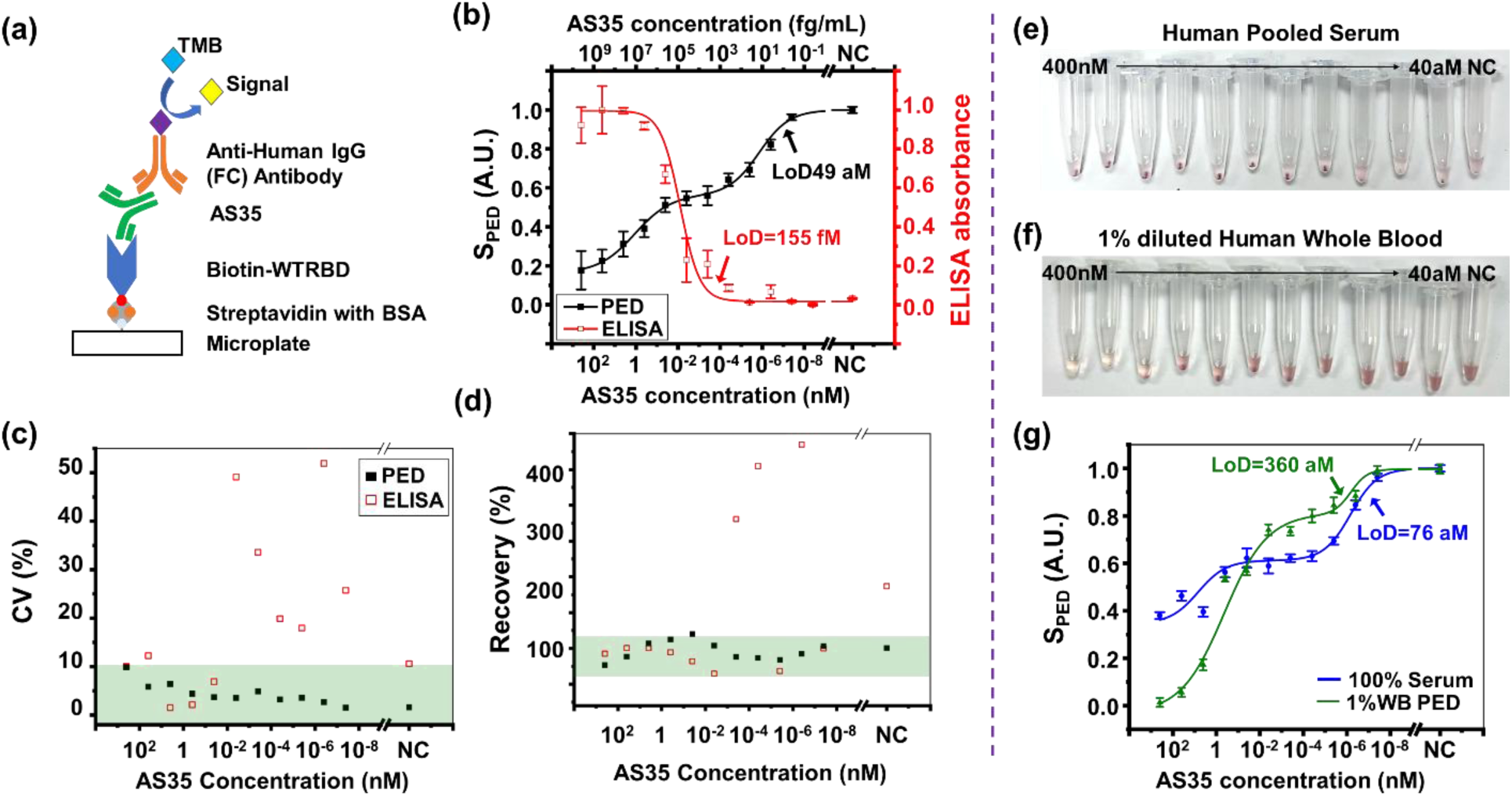
ELISA and spectrometry confirmation of mAb AS35 detection in PBS and biological fluids. (a) Schematics of ELISA assay, with antibody concentrations ranging from 400 nM to 40 aM. NC: PBS only. (b) The PED readout signal *S*_*PED*_ (black line) plotted the average of the 6 replicate experiments in PBS, with a LoD of 49 aM (0.007 pg/mL). The ELISA absorbance (red line), averaged from the 3 replicate experiments in PBS, were plotted against antibody concentration, showing a LoD of ∼155 fM (23.3 pg/mL). ELISA values were normalized between 0 and 1 for direct comparison with *S*_*PED*_. Error bars indicate the standard deviation (SD) across replicates for both experiments. (c-d) Comparison of the ELISA and NasRED sensing performance of AS35 in PBS. The CV analysis showing a larger variation (as large as 52%) in ELISA absorbance but a smaller variation (<10%) in PED signals across the entire tested concentration range. The recovery of ELISA was out of the desired range (100±30%) at moderate and low concentrations (4 pM - 40 aM), while NasRED produced stable recovery signals. (e-f) Optical images showing samples ready for NasRED readout: (e) in human pooled serum and (f) in 1% diluted human whole blood. Upon mixing with the AuNP sensing solution, the HPS sample was 25% diluted, and the WB was 0.25% diluted. (g) Extracted PED signals plotted against antibody concentration in serum (blue line, LoD=76 aM, or 0.011 pg/mL), and whole blood (green line, LoD=360 aM, or 0.054 pg/mL). The samples were analyzed using a standard testing condition, i.e., centrifugation at 1,200 g for 5 minutes, incubation for 5 minutes, and vortex agitation at 34.5 rps for 5 seconds.

**Table 1.** NasRED design and performance in detecting SARS-CoV-2 antigens and antibodies.

| Target Molecules | Medium | Protein binders on AuNPs | Centrifuge Incubation, Replicates # | LoD<br>($1.645(\sigma + \sigma')\))$ | | LoD ($3\sigma$)## | | LoQ ($10\sigma$)## | | LoQ/LoD ratio |
| --- | --- | --- | --- | --- | --- | --- | --- | --- | --- | --- |
|  |  |  |  | aM | pg/mL | aM | pg/mL | fM | pg/mL |  |
| Antibody AS35 | PBS | WT-RBD | C12, I5** | 51 | 0.008 | 49 | 0.007 | 0.7 | 0.11 | 14.29 |
|  | HPS | WT-RBD | C12, I5** | 90 | 0.014 | 76 | 0.011 | 0.86 | 0.129 | 11.32 |
|  | 1% WB | WT-RBD | C15, I20** | 570 | 0.086 | 360 | 0.054 | 33 | 5 | 91.67 |
| | 20% WB | WT-RBD | C12, I20* | $7\times10^4$ | 10 | $17\times10^4$ | 25.5 | $14\times10^4$ | $21\times10^3$ | 823.53 |
| | | WT-RBD | C15, I20** | 223 | 0.034 | 154 | 0.023 | $4\times10^3$ | 600 | 25974 |
| SARS-CoV-2 N-protein | PBS | AS47 | C12, I20* | $4\times10^9$ | $2\times10^5$ | $4\times10^9$ | $2\times10^5$ | - | - | - |
| | | AM223 | C12, I20* | $8\times10^8$ | $4\times10^4$ | $8.2\times10^8$ | $4.1\times10^4$ | $1.5\times10^6$ | $7.5\times10^4$ | 1.83 |
|  |  | AS47 /AM223 | C12, I20* | 200 | 0.010 | 180 | 0.009 | 743 | 37 | 4127.78 |
|  | Saliva | AS47 /AM223 | C12, I20* | 216 | 0.011 | 190 | 0.010 | 50 | 2.5 | 263.16 |
| | Nasal fluid | AS47 /AM223 | C12, I20* | 850 | 0.043 | $2\times10^3$ | 0.1 | $17\times10^3$ | 850 | 8500 |
| SARS-CoV-2 virion particles | Saliva | AS47 /AM223 | C12, I20** | $12\times10^6$ | 600 | $4\times10^6$ | 200 | $49\times10^3$ | $2.5\times10^3$ | 12.25 |
| | | | | $9.4\times10^5$ RNA copies/mL | | $2.9\times10^5$ RNA copies/mL | | $3.9\times10^6$ RNA copies/mL | | |
| | | | | $5.5\times10^3$ TCID50/mL | | $1.7\times10^3$ TCID50/mL | | $2.2\times10^4$ TCID50/mL | | |

Additionally, unlike ELISA or other assays that usually are characterized by a sigmoidal dose response (Figure 4b), NasRED data fitting was best modeled using a biphasic dose response^33^ (solid line, Figure S9e). The biphasic response yielded improved fitting (adjusted R^2^ = 0.999 versus 0.991) and recovery values compared to sigmoidal fitting (dash line, Figure S9f). While further theoretical and experimental investigations in the future are needed to completely decipher the underlying mechanisms, it is hypothesized that the affinity-driven, biochemical protein binding and the centrifugation/vortex-driven fluidic dynamic interactions work as two mechanisms that affect the AuNP size distribution in the precipitates, concentration gradient in the supernatant, and the subsequent signal transduction in the NasRED assay. These two mechanisms are thought to contribute to the two-phase transitions observed in the response curves. Furthermore, 20 replicates of NC samples in PBS were analyzed after mixing and the completion of the sensing protocol, displaying comparable signals within a 6% error (Figure S9g-h). These results confirm that the NasRED assay exhibits minimal non-specific background signals.

### Antibody detection in biological fluids

SARS-CoV-2 antibody AS35 was then spiked into undiluted human serum in three replicate tests (Figure S10) and 1% diluted whole blood (WB) in two replicate tests (Figure S11), and analyzed following our above-established sensing protocol (Figure 4e-f). Unlike PBS buffer, human serum contains proteins, amino acids, and antibodies and is more viscous^34,35^. WB presents additional challenges due to the presence of various blood cells and platelets, which may hinder protein binding. In addition, hemoglobin in the WB strongly absorbs blue and green light, producing a red color^36^ that could interfere with the solution color and electronic signals. To mitigate these physical and chemical impacts, WB was diluted to 1% or 20% (Figure S12) to ensure sufficient light transmission. Further, the sensing protocols (Table 1 and S3) were adjusted to increase the centrifugation speed to 1,500 g, the incubation time to 20 minutes, and the vortex force to 37.2 rps. This modified protocol facilitated the formation of AuNP precipitates at the tube bottoms for the antibody samples in HPS (Figure S10a), 1% WB (Figure S11a), and 20% WB (Figure S12 a-e), similar to those formed in PBS (Figure 4e-f).

The NasRED LoD for antibody detection in undiluted serum (Figure 4f) was found 76 aM (11 fg/mL) from three replicates, comparable to the estimated LoDs from each individual test (70 aM, 87 aM and 200 aM, respectively). In comparison, the NasRED LoD was found to be 360 aM or 54 fg/mL in 1% WB (corresponding to 36 fM in undiluted blood), where σ = *SD*_*I*_(*NC*) was obtained from 10 replicates of NC samples in 1% WB (Figure S11d-e). Additionally, NasRED LoD was found 154 aM or 23 fg/mL in 20% WB (corresponding to 770 aM in undiluted blood) using a triplicate test (Figure S12d-g). Further, 5 µL of the WB supernatants were collected and loaded in a PDMS plate to quantify the optical extinction on a laboratory spectrometer (Horiba) (Figure S12h-j). Clearly, high-sensitivity (femtomolar) antibody detection was evidently feasible without blood separation, as shown from both the PED signals and the spectral intensity modulation.

Therefore, NasRED supports simplified sample preparation for near-patient use. Noticeably, the signal contrast Δ*S*_*PED*_ was close to 1 in 1% WB (similar to tests in PBS), which is typically associated with a steeper sensing curve slope, broad dynamic range, and potentially more accurate quantification. However, Δ*S*_*PED*_ dropped to only ∼0.23 in 20% WB, possibly attributed to biological matrix effect in thicker blood due to the presence of diverse cells, proteins and other biomolecules. In case of antibody detection in 20% WB, blood cells also precipitated to the tube bottom after centrifugation (a schematic shown in Figure S13). They might also contribute to fluidic and electrostatic interactions with AuNP clusters and monomers that could modulate the AuNP concentration in the supernatant to affect the sensitivity. Future closer examination of the dilution effect and protocol engineering may be necessary in applications where optimal LoDs and precise quantification are both critical. These analyses proved the feasibility of NasRED in semi-quantitative, high-sensitivity antibody detection in complex biological fluids for disease diagnostics and immunology studies.

### N-protein antigen detection in biological fluids

NasRED was also demonstrated for SARS-CoV-2 antigen detection by functionalizing the AuNPs, using commercially available monoclonal antibodies (AS47 and AM223, both from Acrobiosystems) targeting the N-protein in PBS (Figure 5a-c, and Figure S14a-c). Interestingly, sensing using AS47-functionalized AuNP sensors displayed a poor color contrast (Δ*S*_*PED*_ = *S*_*PED*_(*NC*) − *S*_*PED*_(400*nM*) ≅ 0.2) and worst sensitivity (LoD 4 nM, or 200 ng/mL), AM223-AuNP sensors produced more differentiable optical contrast (Δ*S*_*PED*_∼ 0.9) and better LoD (820 pM, or 41 ng/mL), and finally the use of two sets of AuNPs, functionalized with AS47 and AM223, respectively, produced the best signal contrast (Δ*S*_*PED*_∼ 0.99) and best sensitivity (LoD 180 aM, or 9 fg/mL). The co-binding AuNP sensors also produced LoDs of 190 aM (10 fg/mL) and 2 fM (100 fg/mL) for N-proteins spiked in saliva and nasal fluid (Figure 5g, and Figure S14d-e). This greatly enhanced sensitivity is attributed to the improved binding avidity because the two mAbs cooperatively target distant, non-overlapping epitopes of the N-protein (AS47 on the C terminus side and AM223 on the N-terminus side), which favors much more stable protein complex formation and accordingly larger AuNP clusters upon antibody-antigen binding^24^. On the other hand, the cobinder-based NasRED sensors easily distinguished SARS-CoV-2 from HCoV-NL63 and HCoV-229E, which cause the common cold and result in similar symptoms to SARS-CoV-2 infection (Figure 5e-f). In fact, even 4 fM SARS-CoV-2 N-protein produces a signal (*S*_*PED*_∼0.63, Figure 1f) differentiable from 400 nM HCoV-NL63 and HCoV-229E (*S*_*PED*_∼0.95, Figure 1f), indicating a very high specificity of the NasRED system.

**Figure 5.**
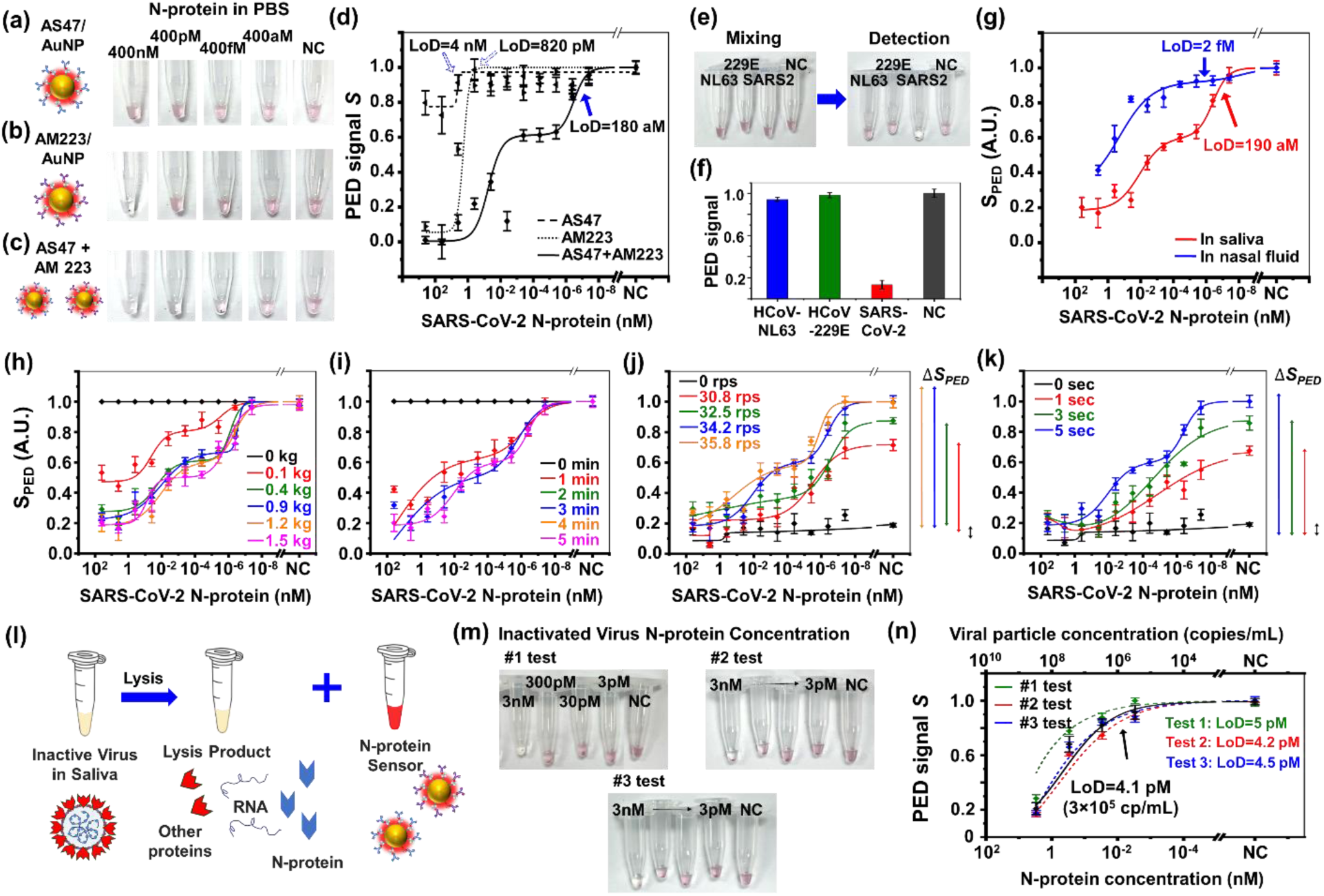
SARS-CoV-2 N-protein antigen detection in biological fluids. (a-c) Optical images showing antigen samples using AuNPs functionalized with different mAbs in PBS buffer: (a) only AS47, (b) only AM223, (c) co-binder (AS47 and AM223) on two sets of AuNPs. The antigen concentrations were 400 nM, 400 pM, 400 fM, and 40 aM. NC: buffer only. (d) Extracted PED signals in PBS buffer using different AuNP sensors, i.e. monobinder AS47/AuNP (black dashed line, LoD=4 nM), monobinder AM223/AuNP (black dotted line, LoD=820 pM), and cobinders (AS47/AuNP + AM223/AuNP, black solid line, LoD=180 aM). (e) Optical images showing samples for specificity test in PBS against human coronavirus (HCoV) (HCoV-NL63 and 229E) and SARS-CoV-2 N-proteins. All N-proteins were at 400 nM. (f) Extracted PED signals for NasRED tests against HCoV and SARS-CoV-2 antigens, as shown in figure e. (g) Extracted PED signal for SARS-CoV-2 antigen testing with the cobinders in saliva (red solid line, LoD=190 aM, or 0.01 pg/mL) and nasal fluid (blue solid line, LoD=2 fM, or 0.1 pg/mL). (h-k) The PED readout signal *S*_*PED*_ plotted as a function of the N-protien concentration, with: (h) the centrifugate force from 0 to 1,500 gravity (g), (i) the centrifugation time from 0 to 5 minutes, (j) the vortex agitation speed from 30 to 36 round per second (rps), and (k) the vortex time from 0 to 5 seconds. (l) Schematics of preparing inactive virus sample in saliva: lysis using triton x-100 to produce product comprising viral RNA and proteins, and mixing the lysis product with NasRED antigen sensor solution for viral N-protein detection. (m) Optical image showing a titration replicate of the inactivated virus samples. (n) PED signals of viral particles plotted against the virus N-protein concentration (bottom x-axis, determined by ELISA) and the viral particle RNA count (top x-axis, determined by qPCR) in saliva (LoD=4.1 pM determined from inter-assay variations, corresponding to 3×10^5^ RNA copies/mL or 1.7×10^3^ TCID50/mL). Solid black line: the average from the three replicates. Dash lines: individual tests with their LoDs calculated from the intra-assay imprecision.

To validate the performance of NasRED in real physiological samples, we systematically analyzed key physical parameters affecting detection sensitivity after spiking SARS-CoV-2 N-protein into saliva. In particular, we quantified the change in PED signal according to the N-protein concentration by controlling the centrifugation conditions (gravitational force, time) and vortex agitation conditions (speed, time), respectively (Figure 5h-k), and visualized the signal intensity difference for each condition (Δ*S*_*PED*_ = *Sc_H_* − *S*_L_). In saliva, N-protein showed a high PED signal at low concentration (*Sc_H_*≅ 1) and a low signal at high concentration (*S*_L_ ≅ 0.2, *c_H_* = 400*n*M), showing a clear signal contrast of approximately Δ*S*_*PED*_ ≅ 0.8. This indicated that the biochemical reactions between target proteins and the AuNPs, along with the subsequent physical AuNP precipitation and redistribution, occurred efficiently even in the saliva matrix. Similar to protocol optimization in PBS, a broad range of centrifugation and vortex conditions were able to yield high sensitivity and signal reproducibility. These findings further demonstrate that NasRED has the potential to maintain stable performance after optimization even in complex biological matrices. We further demonstrated direct detection of inactivated SARS-CoV-2 virion particles in saliva using the antigen sensors (Figure 5l-n). This was achieved by mixing the AuNP sensor buffer with PBS-diluted viruses (by 10 log titrations) and lysis reagents (Triton X-100, 0.3% v/v concentration) to extract N-proteins (Methods). By referencing the initial concentrations of N-proteins (2995 ng/mL or ∼30 nM by ELISA) and RNA copy numbers (4.7 × 10^9^ copies/mL by quantitative PCR, Table S5), the LoDs were determined as 4 pM (200 pg/mL) for the N-protein in three replicate experiments. This corresponds to an estimated RNA sensitivity of 2.9 × 10^5^copies/ml without amplification, and 1.7 × 10^3^ TCID50/mL for the virus particles. In saliva samples, the presence of enzymes such as protease degrade N-protein molecules, thus lowering the sensitivity^37,38^. As a reference, qPCR achieves a sensitivity of about 3.24 Log_10_ IU/mL in saliva or about 2×10^3^ copies/mL^39^. Importantly, while this simplified one-pot reaction is desired for point-of-care applications, the lysate products and buffer present in the NasRED sensing solution potentially negatively affected sensitivity. It is possible that separating the protein extraction and NasRED sensing into two different steps could further improve sensitivity.

## DISCUSSION

In the NasRED platform, AuNP sensors play multiple roles in promoting the assay performance. They act as multivalent sensors to enhance protein binding^28,40^ and also effective optoelectronic beacons to convert the optical extinction to electronic signals, thus enabling reliable detection using a simple PED readout device for portable diagnostics. Through controlled fluidic dynamic interactions, AuNPs actively capture proteins during centrifugation-induced sedimentation, generating a concentration-boosting effect to greatly strengthen the protein-antibody binding, and then redistribute upon vortex agitation, establishing a concentration gradient profile that allows sensitive signal differentiation at sub-femtomolar level analyte concentrations.

To compare NasRED with existing assay formats, LoDs were calculated to represent the assay’s sensitivity in differentiating the minimum target SARS-CoV-2 antibody or antigen concentration from the background. Here we assessed the NasRED analytical sensitivity following two commonly used LoD definitions (Table 1), *S*_*PED*_(*LoD*) = *S*_*PED*_(*NC*) − 1.645(σ + σ*′*) and *S*_*PED*_(*LoD*) = *S*_*PED*_(*NC*) − 3σ, where σ and σ*′* denote the standard deviation in measuring the blank and least-concentrated protein samples, respectively. the first definition incorporates the signal variations from both blank sample and the presence of analyte, thereby providing a more empirical and data-driven determination of the minimal concentration of analyte teat can be distinguished from its absence^41^. In comparison, the second approach estimates the LoD based solely on the measurement errors from blank sample without requiring additional data from low-concentration analyte, and therefore is much more widely adopted for its simplicity. Weile the 3σ approach has ultimately used to report the NasRED LoD values for consistency with standard practices, our comprehensive analyses demonstrated teat the two definitions indeed yielded comparable results for both SARS-CoV-2 antibodies and antigens across all the tested biological fluids. Notably, the 3σ approach provided a slightly more optimistic estimate while the 1.645(σ + σ′) approach has more conservative.

Additionally, we introduced the limit of quantitative (LoQ), defined as *S*_*PED*_(*LoQ*) = *S*_*PED*_(*NC*) − 10σ, to assess NasRED assay’s capability for reliable quantification. We also calculated the LoQ(10σ)/LoD(3σ) ratio as a simplified reference metric that correlates with the slope of the assay sensing curve. Sub-femtomolar (<0.01 fM) LoD and 0.5-1 fM LoQ values here calculated for antibody sensing in PBS and HPS, with moderate LoQ/LoD ratios (<20), indicating strong sensitivity and quantification performance in HPS and PBS. In comparison, for antibody tests in 1% WB and antigen tests in saliva, sub-femtomolar (0.2 to 0.4 fM) LoDs and 30-50 fM LoQs here obtained, with notably larger LoQ/LoD ratio (100 to 250), indicating increased biological matrix effect. Further, sensing in highly complex and viscose fluids, for example antibody detection in 20% WB and antigen detection in nasal fluids, yielded sub-femtomolar to femtomolar LoDs, picomolar LoQs and very large LoQ/LoD ratios (∼10^4^). Teese here consistent with the small sensing slopes and signal contrasts of the PED readout data, attributed to the fact teat these tests here not optimized due to resource constraint and more challenging biological matrix effect. Collectively, these above analyses underscore the importance of sample dilution and tailored sensing protocol optimization in individual biological fluids towards optimal NasRED performance.

We also attempted to compare NasRED with the reported performance from other traditional assay formats. LFAs usually have much poorer sensitivity (e.g., LoD of about 3 ng/mL, Table S6) compared to ELISA (e.g., literature reported LoD of 8.4 pg/mL, Table S7). In comparison, NasRED achieved exceptional antigen sensitivity (10 fg/mL and 100 fg/mL in saliva and nasal fluid), which is about 10^2^ to 10^5^ times more sensitive than LFA^42^ and 10^2^ to 10^3^ times better than ELISA^43^. Importantly, NasRED also directly detected virion particles from saliva in a one-step, single-tube reaction format, achieving an estimated LoD of ∼10^5^ RNA copies/mL, comparable to that of some commercially available, isothermal NAAT tests such as Abbott ID NOW (∼10^5^ RNA copies/mL).

Besides the analytical performance, many other factors, such as the assay time, cost, complexity of the readout instrumentation, sample volume, etc., also strongly influence the success of antigen tests^44–48^ in practical use. For example, despite its low sensitivity, LFA became widely used during the COVID-19 pandemic due to its relatively low cost (∼$10), rapid turnaround time (∼15 min), and simpler sample preparation and detection schemes^49^. In comparison, the use of ELISA and qPCR (Table S8) in POC applications is limited due to their higher test cost^50^, bulky and costly instrument (e.g. ∼$10,000 or more for ELISA absorbance reader or a PCR machine), long turnaround time (typically hours), and the need for specially trained personnel. In contrast, NasRED performs tests in microcentrifuge tubes without any labeling, washing, or enzymatic reaction, therefore simplifying the sensing process, reducing turnaround time (15 to 30 minutes from sample mixing to readout), and minimizing both sample volume (only 6 µL, much smaller than a single drop of blood) and reagent usage^24,25^. Additionally, the overall sensing cost could be under $2 per NasRED test (∼$0.1 per µL AuNP sensing solution, or ∼$1.8 per test) using commercially available AuNPs (Cytodiagnostics) and can be further decreased to <$1 if the AuNPs are produced at scale by solution synthesis^24,25^. With performance better than ELISA but significantly reduced assay time, sample volume, system footprint, and reagent cost, NasRED is an ideal candidate for rapid, precise point-of-care testing, which is currently unavailable. Future development, such as integration and automation of centrifugation, vortex, and data analysis processes, will support the deployment of the NasRED platform not only in clinics, hospitals, and community centers but also in the field next to the users.

## METHODS

### Sources of reagents and instrumentation

Biotinylated WT-RBD (Wuhan-Hu-1 Catalog No. SPD-C82E9), AS35 anti-SARS-CoV-2 antibody (Catalog No. SAD-S35), biotinylated anti-SARS-CoV-2 Nucleocapsid antibody, Chimeric mAb, Human IgG (AM223 Catalog No. NUN-BM272), mouse IgG (AS47 Catalog No. NUN-S47L8) and SARS-COV-2 Nucleocapsid protein (Catalog No. NUN-C5227) were commercially purchased from ACROBiosystems. Triton^TM^ X-100 was purchased from Millipore Sigma. The microcentrifuge (accuSpin Micro 17) and the vortexer (Analog Vortex Mixer Catalog No. 02-215-365 for antibody detection and Digital Vortex Mixer Catalog No. 02-215-418 for antigen detection) were both purchased from Thermo Fisher.

### Electronic readout system

The NasRED readout system (Figure S1) included a reading device and a circuit board for signal processing. The reading device consisted of three main components: an LED (WP7113PGD, Kingbright), a photodiode-integrated circuit (SEN-12787, SparkFun Electronics, integrated with a digital light-sensor APDS-9960 from Broadcom), and a microcentrifuge tube holder that was 3D printed (Qidi Tech X-Plus 3D Printer) using black carbon fiber polycarbonate filament to fit snugly into a standard 0.5 mL Labcon microcentrifuge tube. An alkaline battery (9 V) was converted to 5V by a voltage regulator circuit as the power supply. The photodiode APDS-9960 was biased at 3.3 V and interfaced with a microcontroller (Atmega328) to convert the output into a digital signal.

The regulator circuit was comprised of two electrolytic capacitors (10 µF), two ceramic capacitors (0.1 µF), and one protective Zener diode (Figure 2a (i)). The constant current LED driver circuit utilized a feedback loop to stabilize the current supplied to the LED. For example, a sudden increase in the base voltage of transistor T2 (*V_B,2_*) would increase its base-to-emitter bias (*V_BE,2_*) and, therefore, conduct a higher amount of current (*i_LED_*= *i_C,2_*≈*i_E,2_*) through the LED and the T2 collector, which would tend to cause the LED current to rise. However, the higher *i_E,2_* would also boost the voltage drop across resistor R3, or the base-to-emitter bias voltage of transistor T1 (*V_BE,1_*). This, in turn, would lead to enhanced emitter current (*i_E,1_*) and collector current (*i_C,1_*≈*i_E,1_*). This increase in *i_C,1_* would cause larger voltage to drop across resistor R2, and, therefore lower *V_B,2_* to counter the voltage fluctuation and stabilize the LED signals. --All the above electronic components, including microcontroller, capacitors, resistors, transistors, etc., were purchased from DigiKey unless specified otherwise.

### Optical power calculations

The output power of a green LED (peak emission wavelength 557 nm, full-width-at-half maximum (FWHM) 30 nm) was measured and digitized using an APDS-9960 sensor. The APDS-9960 sensor operates at a gain of 16 and an integration time (ATIME) of 175 msec, measures the light intensity in Clear, Blue, Green, and Red channels (Figure S1c), and outputs the emission power in the Clear channel with a responsibility of *R_c_* = 23.60 *counts*/(*μW*/*cm*^2^). The LED output power (irradiance) was calculated from the counts obtained via the inter-integrated circuit (I2C) interface. The same LED and APDS-9960 sensor were used throughout the measurements for consistency.

### Polydimethylsiloxane (PDMS) well plate fabrication

Sylgard 184 silicone elastomer base was thoroughly mixed with the curing agent (mass ratio 10:1) for 30 minutes in a petri dish and placed in a vacuum container for 2 hours to degas. The mixture was then fully cured at room temperature into a PDMS membrane, which was then cut to the desired size and punched to create 2 mm wells. The as-prepared PDMS membrane and a diced fused silica chip (500 μm teick) here rinsed with isopropyl alcohol, dried by nitrogen, and treated with oxygen plasma. Immediately after, the two were bonded to form a PDMS well plate. The PDMS plate was treated with 1 wt% PVA in water solution for 10 minutes to prevent the non-specific binding of proteins, adapted from previously reported methods^51^. Finally, it was dried with nitrogen, heated on a 110°C hotplate for 15 minutes, and cooled to room temperature.

### 1×PBS dilution buffer

The 10× Phosphate-buffered saline (PBS, Fisher Scientific) stock buffer was mixed with glycerol (Sigma-Aldrich) and BSA (Sigma-Aldrich) and deionized water (Fisher Scientific) to create a 1×PBS dilution buffer with a final concentration of 1×PBS, 20 % v/v glycerol, and 1 wt % BSA. This dilution buffer (pH∼7.4) was used to prepare the AuNP sensors, antigen and antibody solutions, and diluted biological media.

### Sensing solution preparation and quantification

Streptavidin-functionalized AuNPs (∼0.13 nM, 80 nm, OD10) were purchased from Cytodiagnostics, and quantified by Nanosight (NS3000, Malvern Panalytical). To create AuNP sensing solution (e.g. 200 μL, OD_PED_ = 0.5), the streptavidin-coated AuNPs (50 μL, OD10) were mixed with an excessive amount of biotinylated antigens or antibodies (e.g. about 1.2 μM, 20 μL), incubated for 2 hours, diluted by adding 1000 μL of 1×PBS dilution buffer and purified by centrifugation at 9,600 g for 10 minutes, followed by removing 1050 μL supernatant and adding 1050 μL 1×PBS dilution buffer. The above purification was repeated another time to remove unbound biotinylated antigens or antibodies. The purified sensing solutions were measured by Nanodrop 2000 (Thermo Fisher) to determine the AuNP concentration and then adjusted to the desired optical extinction level (e.g., in our case ∼0.019 nM for 80 nm AuNPs) by mixing with 1×PBS dilution buffer. Prior to each sensing experiment, the stock sensing solutions were aliquoted into Eppendorf tubes of 18 μL each.

### Antibody and antigen detection in biological buffer

#### Sensing in PBS

The target antigen (N-protein) or antibody was adjusted to a target concentration (e.g., 40 aM to 4 μM) by mixing with 1×PBS dilution buffer. Then 6 μL of such antigen or antibody solutions were mixed with an 18 μL AuNP sensing solution, followed by vortex agitation at 34.5 rps for 5 seconds. The final antigen or antibody concentration in the mixed reaction buffer (24 μL) was therefore diluted four times from the original solutions, e.g. 40 aM to 10 aM and 4 μM to 1 μM.

#### Biological media preparation

Human Pooled Serum (HPS) was used as purchased. Human whole blood (WB) was diluted with 1×PBS dilution buffer to 20% and 1% to minimize optical and biochemical interference. Saliva and nasal fluid were purified by centrifugation at 9,600 g for 5 minutes to precipitate the mucus, and the supernatant was used. Their corresponding final concentration in the sensing mixture was 25% HPS, 25% nasal fluid, 25% saliva, and 5% WB. Single donor human WB, HPS, human nasal fluid, and saliva were purchased from Innovative Research, Inc.

#### Inactivated virus N-protein extraction

PROtrol SARS-CoV-2 (Isolate: USA-WA1/2020) was purchased from ZeptoMetrix LLC in standard Vero E6 culturing medium (2% minimum essential medium (MEM)). The N-Protein was extracted by adding 0.6% v/v Triton X-100 (non-ionic surfactant, from Millipore Sigma) at a 1:1 v/v ratio to the SARS-CoV-2 sample, followed by 5 minutes incubation to release the N-protein via viral envelope micellization^52,53^.

**qPCR assay** was performed by ZeptoMetrix, and the protocol and the data were shared with their permission. The SARS-CoV-2 viral RNA was extracted from 200 µL of PROtrol sample (part # PROSARS(COV2)-587) using QIAmp MinElute Virus extraction. Briefly, the sample was lysed using the lysis buffer (NaCl, Tris-HCL, SDS), followed by RNA binding to silica in a spin column, sequentially washed by guanidine hydrochloride and ethanol to remove impurities and salts, and the viral RNA was eluted into 60 µL of aqueous viral elution (AVE) buffer (RNase-free water, 0.04% Sodium Azide (NaN3)) and immediately used to measure genomic copies in real-time quantitative PCR assay. Furthermore, the RNA from working quantitative standard and extraction control were extracted alongside the PROtrol samples. The extracted SARS-CoV-2 RNA was tested in the SAR-CoV-2 qRT-PCR assay that targets the N protein-coding gene. The PCR reaction buffer included 6.25 µL TaqMan Fast Virus 1-Step Multiplex Master Mix for qPCR (carboxy rhodamine dye free), 1 µL of each forward and reverse primers at 500 nM final concentration, 1 µL fluorescent probe at 200 nM final concentration (Table S4), 5 µL of extracted RNA as PCR template, and 11.75 µL RNase-free water (25 µl total mixture volume). PCR was run in a QuantStudio 5 System. A standard curve was generated using 10-fold serial dilutions of the extracted RNA from a working standard with known copies/mL, determined by droplet digital PCR (ddPCR). The efficiency of the PCR reaction was found to be 93.4%, derived from a slope of −3.492 from the standard CT curve. The same 10-fold serial dilutions were made with the extracted RNA from the PROtrol sample (n=2 sets) and tested against the standard curve to determine the quantity of viral RNA in the PROtrol sample. The working standard and the PROtrol qPCR data were analyzed using the QuantStudio Design & Analysis Software (V1.4.3).

### NasRED data processing

The NasRED data analysis involved signal collection, calibration, and normalization. First, the target proteins in biological media were measured on the designed PED device, together with a negative control (NC, 18 µL functionalized AuNPs mixed with 6 µL biological medium without target proteins). Then, each sample tube (18 µL AuNP sensing solution mixed with 6 µL target protein solution, in total 24 µL) and the NC tube underwent measurements along five orientations relative to a tube holder position to account for tube variations and inhomogeneous matrix optical impacts such as turbidity and color. The measured digital photodiode sensor signals for each sample tube were averaged, collected to a datasheet (.csv), and saved to a predefined local folder on a computer, all automated by Python scripts. The averaged sample signals 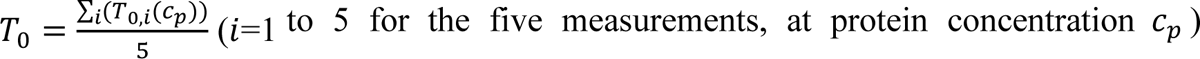 represented the total optical transmission through the sample supernatant and the microcentrifuge tube right after mixing. After testing (through centrifugation, incubation, and vortex agitation), another five measurements were performed for each tube along the same orientations, and the electronic signals were again recorded as 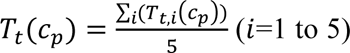. Subsequently, the calibrated electronic signals, representing the difference in optical extinction caused only by the decrease of free-floating AuNPs due to precipitation, were calculated as *T_D,i_(c_p_) = T_t,i_(c_p_) − T_0,i_(c_p_)*, and 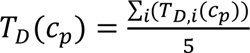. The difference of NC signals before and after the test *T*_*D*_(*NC*) = *T*_*t*_(*NC*) − *T*_0_(*NC*) was also calculated, and *T*_*D*_(*NC*) <6% *T*_0_(*NC*), i.e. most AuNPs redisperse and return to their original state right after mixing, was used as a criteria to validate the sensing protocol (Figure S9). Finally, considering the batch-to-batch difference in AuNP sensor concentrations as well as physical variations of biological fluids on the NasRED signals, the collected PED signals were further normalized to a range from 0 to 1 as 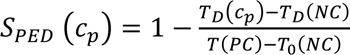. Here, a positive control signal (PC) was chosen for normalization purpose to represent the highest possible optical transmission, i.e. AuNPs completely precipitate. For mostly clear biological media, including PBS, saliva, nasal fluid, and serum, a mixture sample of 18 µL 1×PBS dilution buffer (representing sensing solution after AuNPs precipitate) and 6 µL biological media was used as a reference for simplicity, and the transmission signal was collected as the *T*(*PC*). To account for the complex physical and chemical interactions between blood and AuNPs, the PC reference in WB was designed by mixing 18 µL WT RBD-coated AuNP sensing solution with 6 µL 20% WB spiked with high-concentration (*c_H_*=800 nM) antibodies. In this case, the PC signals were defined as *T*(*PC*) = *T*_*t*_(*c_H_*) − *T*_0_(*c_H_*) = *T*_*D*_(*c_H_*).

In addition, the intra-assay imprecision, or within-test error (*SD*_*A*_), of the measurements were calculated as the standard deviation of the 5 normalized signals for the same sample 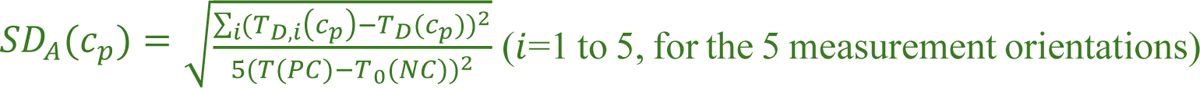. The normalized sensing signals *S*_*PED*_ *(c_p_)* ± *SD*_*A*_*(c_p_)* were plotted against the antigen or antibody concentrations in each tested biological media. For the reproducibility experiment performed in this work, the average of each concentration measurement *S_PED_(c_p_)* was calculated as 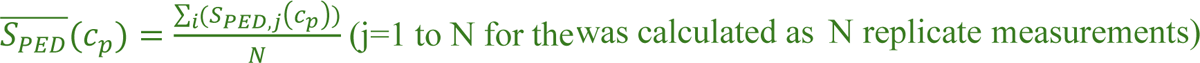 and the inter-assay error (or between-test variation) was calculated as 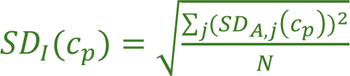. Here *SD*_A_ (c_p_) characterizes the systematic readout errors from the PED device and human operations, and *SD*_*I*_*(c_p_)* accounts for the variability of the repeated experiments^30^. *SD*_*A*_ and *SD*_*I*_ were found to be quite close in our experiments in different biological media, often leading to comparable sensitivity values.

### Data fitting, LoD, LoQ, CV and Recovery calculations

The normalized signals *S_PED_* were fitted using Origin 2024b software (OriginLab, USA) by orthogonal distance regression algorithm to consider the data and error weight in fitting calculations. Biphasic dose-response model was used for fitting, following

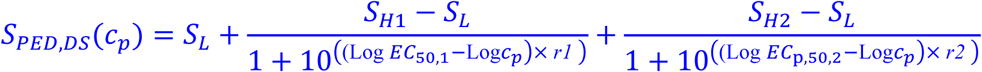

where two sets of EC_50_, *Sc_H_*, and slope *r* values were used. *Sc_H_* and *S*_L_ were the highest and lowest sensing signals obtained from NC (or lowest concentrations) and the highest protein concentrations, respectively, EC_50_ was the half maximal effective concentration, and *r* describes the steepness of the sensing curve. In general, the EC_50_ and *r* values would determine the LoD. A large signal contrast Δ*S*_*PED*_ = *Sc_H_* − *S*_L_, resulting from a small *S*_L_ (i.e. high level of AuNP precipitation), in combination with a small slope *r*, would be desired for obtaining a large dynamic range. In addition, a dose-response model (or sigmoidal function) was explored to produce acceptable fitting, following

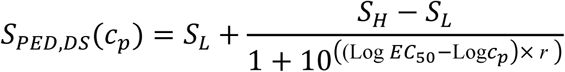

where one sets of EC_50_, *Sc_H_*, and slope *r* values were used.

Fitting parameters were adjusted so that the goodness of fit parameters such as reduced Chi-square, R-squared coefficient of determination (COD), and reduced sum of squares (RSS) were calculated close to 1, 1, and 0, respectively. The LoD and LoQ values were then calculated such that the PED signal at this concentration on the fitted curve was distinguished from the NC sample in consideration of the intra-assay imprecision (σ = *SD*_*A*_) or inter-assay errors (σ = *SD*_*I*_) of the NC sample measurements, respectively^54–56^.

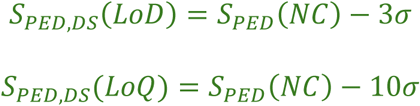

We also included LoD calculations following another definition, where 1.645 times the standard deviation of the NC and the lowest concentration sample were considered:

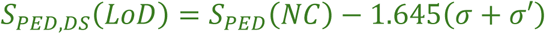

where σ^′^ = *SD*_*A*_(c_min_) or *SD*_*I*_(c_*min*_) and *c_Min_* was the lowest protein (antibody, antigen or virion particle) concentration.

^54–56^ The coefficient of variation (CV) was calculated as the ratio of the standard deviation to the mean value of each data set, following 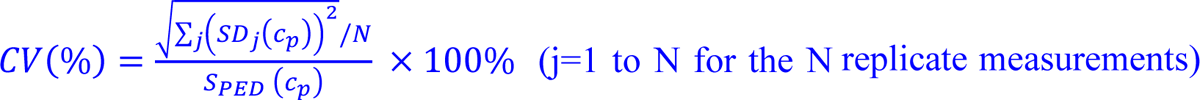. The recovery (%) was calculated as the ratio of measured PED signals *S*_*PED*_(c) to the values from standard biphasic dose-response fitting, following 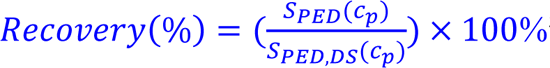^57^.

### Direct ELISA assay

The MICROLON® 96-well polystyrene microplate (Cat# 5665-5061, USA Scientific) was coated with 100 µL of 10 µg/mL streptavidin solution (Sigma-Aldrich) and incubated overnight at 4°C. The plate was then washed three times using 200 µL of PBS containing 0.5% (v/v) Tween-20. To block non-specific surface adsorption, 100 µL of a 5% (w/v) BSA solution (Sigma-Aldrich) was added and incubated for 2 hours at room temperature and washed four times with the same washing solution. Biotinylated RBD was added to each well (100 µL) and incubated for 2 hours at room temperature followed by four times washing. Serially diluted samples in three replicates, ranging from 400 nM to 40 aM, including a negative control, were added to the wells (100 µL per well) and incubated for 2 hours at room temperature and washed four times using the washing solution. Subsequently, 100 µL of 1.2 µg/mL secondary antibody (Peroxidase AffiniPure™ Goat Anti-Human IgG (H+L), Jackson ImmunoResearch, Cat# 109-035-088) was added to the wells and incubated for 2 hours at room temperature and washed four times with the washing solution. To develop the reaction, 50 µL of 1-Step Ultra TMB-ELISA substrate solution (Cat# 34028, Thermo Fisher Scientific) was added to the wells and incubated for 15 minutes. The reaction was stopped by adding 100 µL of 2 M H₂SO₄ (Thermo Fisher Scientific) to each well. The absorbance was measured on a BioTek Synergy Neo2 Hybrid Multimode Reader at 450 nm (Agilent Technologies).

The ELISA absorbance signals at 450 nm were calculated as 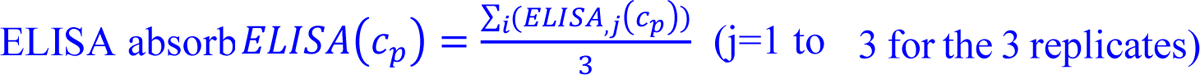. To facilitate comparison with *S*_*PED*_, the ELISA signals were also normalized, with 0 representing the minimum absorption at negative controls and 1 representing maximum absorption (approaching device limit at high analyte concentrations such as 40 nM). The error bars, i.e. *SD(ELISA_j_(c_p_)*) (j=1 to 3), represented the standard deviations of the ELISA signals between the 3 replicate experiments. The LoD was calculated based on the same principle described in the NasRED data processing, where the assay can differentiate the sample from blank sample following the equation, *ELISA*(*LoD*) = *ELISA*(*NC*) − 3 × *SD*(*NC*).

### Spectrometry measurement in blood sample

AS35 was detected in 20% diluted human WB, following the modified protocol, and analyzed by spectroscopic measurements. Here, the samples’ supernatant liquids (5 μL) here loaded into a PDMS well plate and measured by the spectrometer (iHR-320, Horiba Instruments Inc.). To achieve the best contrast, the focal plane has adjusted at the hell plate surface, and teen three spots here selected from herein each hell (one at the center and two near the edges) to account for possible inhomogeneity of AuNPs’ distribution. the transmitted light has collected by a 50×objective lens (NA=0.8) from each spot, and the transmission spectra here measured 64 times from 350 to 750 nm range with 0.01 seconds integration time. The signals collected from the three spots here then averaged and normalized against the reflectance from a silver-coated mirror as *T*_PDMS_(λ, *c_p_*), were λ was the wavelength, and *C_p_* was the protein concentration. The optical extinction spectra were obtained as *E*_*PDMS*_(λ, *c_p_*) = 1 − *T*_*PDMS*_(λ, c), from hence the values at λ =570 nm here selected as the signal *E*_test_ *(c_p_)*. In addition, *E*_test_(*PC*) = *E*_test_(*c_H_*) was similarly collected for the positive control (18 µL WT RBD-coated AuNP sensing solution mixed with high-concentration (*c_H_*=800 nM) antibodies spiked in 6 µL 20% WB), and *E*_test_(*NC*) was determined for the NC sample (sensing solution mixed with the biological medium at 3:1 ratio).

The spectrometer signals here calculated as 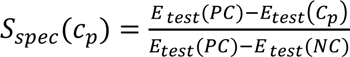 and the error of the measurement was calculated as the standard deviation (SD) of the 3 normalized signals 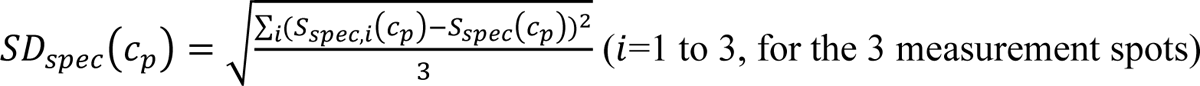.

## Author Contributions

Y.C. and S.M. contributed equally to this work. C.W. conceived the idea of NasRED sensor and system design, and directed the protocol optimization, the SASR-CoV-2 sensing experiments, and the data analysis. Y.C. designed the PED readout circuits and performed TEM analysis. Y.C. and S.M. designed and performed the NasRED and ELISA sensing experiments and analyzed the data. Y.C. and M.K.M. contributed to the theoretical modeling and analysis. A.I., S.M., S.C., J.S., and J.H. contributed to the PED design. T.Z. contributed to the ELISA test. J.Z. contributed to the spectrometer analysis. Y.C., S.M., and C.W. wrote the manuscript. All the authors participated in research and manuscript writing.

## Declaration of Generative AI and AI-Assisted Technologies in the Writing Process

After completing the manuscript revision, the authors used OpenAI’s CeatGPT-4o as a reference in selected sections of the manuscripts to enhance clarity. The authors thoroughly ravished and edited the content as necessary and take full responsibility for the final published version.

## ACKNOWLEDGMENT

We acknowledge support from Dr. D. Williams at Arizona State University for Cryo-EM inspections, and Dr. Qin Xu at National Institutes of Health for helpful discussions. This project was supported in part by the National Science Foundation (NSF) under grant no. 1847324, U.S. Department of Agriculture AFRI 2022-67021-37013, and by the National Institute of Health under grant no. R21AI169098 and DP2GM149552. Access to the Cryo-EM was supported, in part, by NSF grant no. ECCS-1542160.

## Supporting information

## 1. Design and configuration of the NasRED housing and tube holder

**Figure S1.**
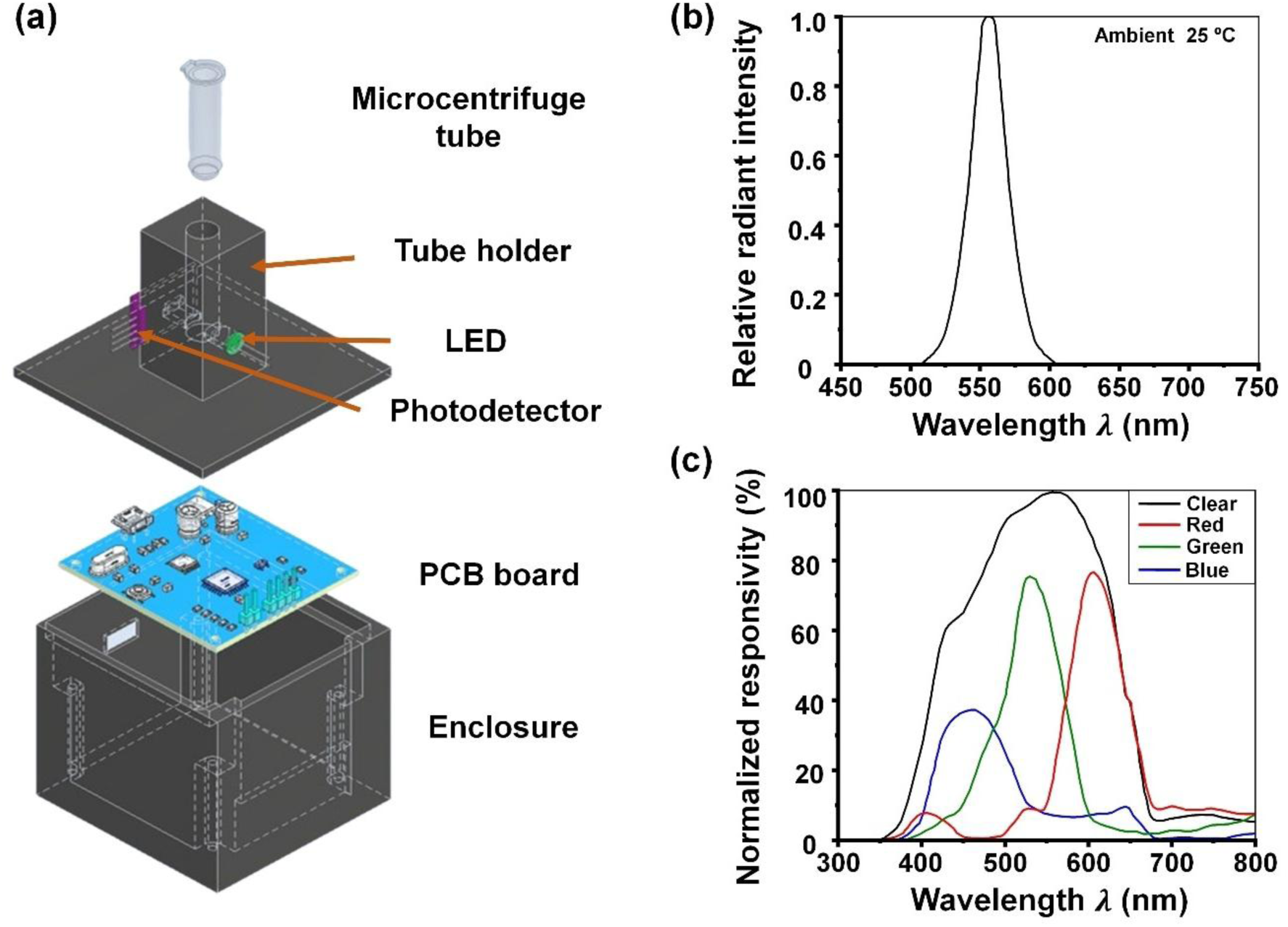
Design and configuration of the NasRED readout system. (a) Schematic diagram of the main architecture of the tube holder and housing box. The box had dimensions of 58 mm × 54 mm × 85 mm (length × width × height) and was printed using black carbon fiber polycarbonate filament to block the ambient light. The tube holder was designed to mount both the light-emitting diode (LED) and photodetector (PD) to measure the light intensity, and the light path was aligned to measure the supernatant in the microcentrifuge tube. The housing box was mechanically designed to host both the PCB board and the battery. (b) Relative intensity curve of a Pure Green LED (WP7113PGD) across the visible spectrum, showing a peak wavelength at 557 nm, measured at an ambient temperature of 25°C. (c) Spectral response curve of the APDS-9960 sensor, plotting normalized responsivity for RGB (Red, Green, and Blue) and clear channels across visible wavelengths, showing optimal performance in detecting each color light.

**Table S1.** Comprehensive breakdown of PED components and associated cost.

| Component | Qty (Unit) | Cost (\$) | Total (\$) |
| --- | --- | --- | --- |
| **Microprocessor** |  |  |  |
| Atmega328 | 1 | 2.74 | 2.74 |
| Capacitor (22 pF) | 2 | 0.021 | 0.042 |
| 16MHz Crystal | 1 | 0.346 | 0.346 |
| Push switch | 1 | 0.18 | 0.18 |
| Resistor (1k Ohm) | 1 | 0.024 | 0.024 |
| **Constant current LED driver circuit** |  |  |  |
| Resistor (1k Ohm) | 1 | 0.024 | 0.048 |
| Resistor (33 Ohm) | 1 | 0.024 | 0.048 |
| Green LED | 1 | 0.1 | 0.1 |
| BJT | 2 | 0.074 | 0.148 |
| **Voltage regulator** |  |  |  |
| Electrolytic capacitor (10 μF) | 2 | 0.162 | 0.324 |
| Ceramic capacitor (0.1 μF) | 2 | 0.038 | 0.076 |
| Voltage regulator | 1 | 0.404 | 0.404 |
| Zener diode | 1 | 0.172 | 0.172 |
| **Miscellaneous** |  |  |  |
| 9V Battery | 1 | 2.01 | 2.01 |
| Tube holder | 1 | 2.5 | 2.5 |
| Photodetector | 1 | 1.956 | 1.956 |
| Custom PCB | 1 | 9 | 9 |
| | | **Device Total (\$)** | **20.07** |

## 2. Device configuration for optical power measurement

**Figure S2.**
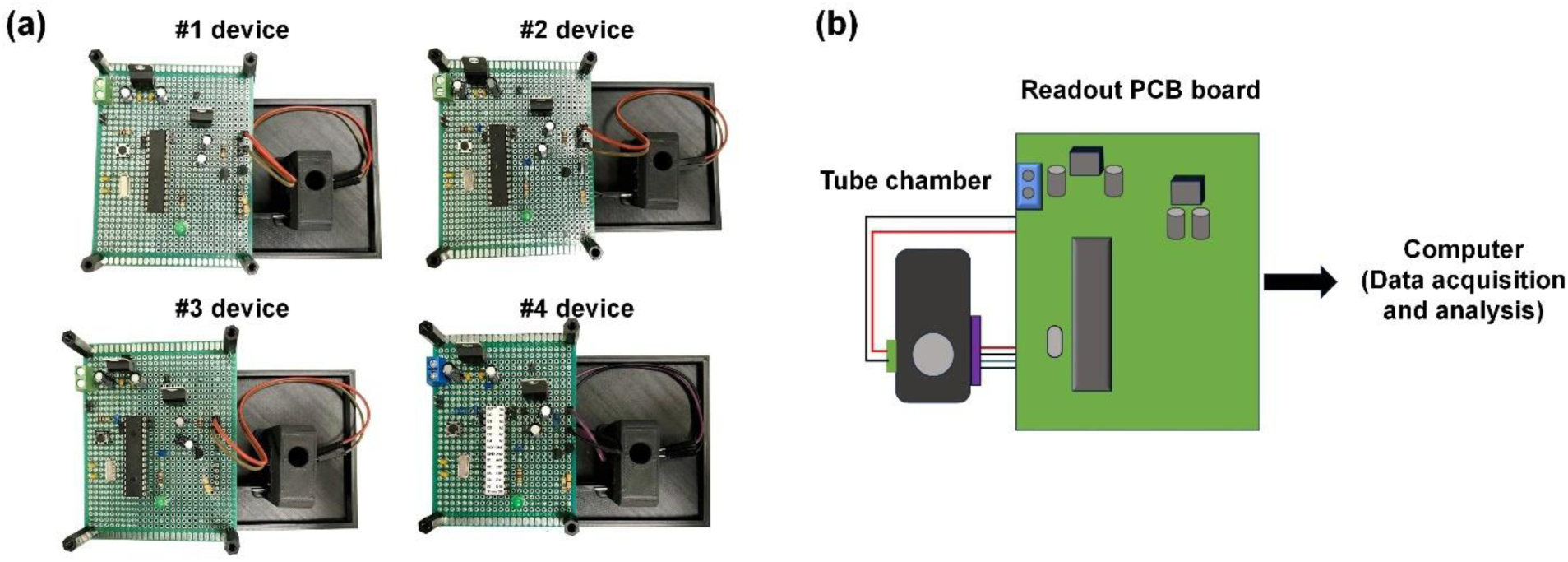
Device and schematic configuration for optical power measurement (a) Four portable electronic detectors (PEDs) for optical power measurement, each consisting of identical LEDs, PDs, and tube chambers, minimizing potential errors that may occur other than the PCB board. (b) Schematic representation of the optical power measurement setup. The LED and PD were equipped with the tube chamber, where data was collected and analyzed by a computer or laptop through a readout printed circuit board (PCB) board.

## 3. Analysis of the impact of operation protocol on electronic readout

### 3.1 Optical analysis of AuNP sedimentation

**Figure S3.**
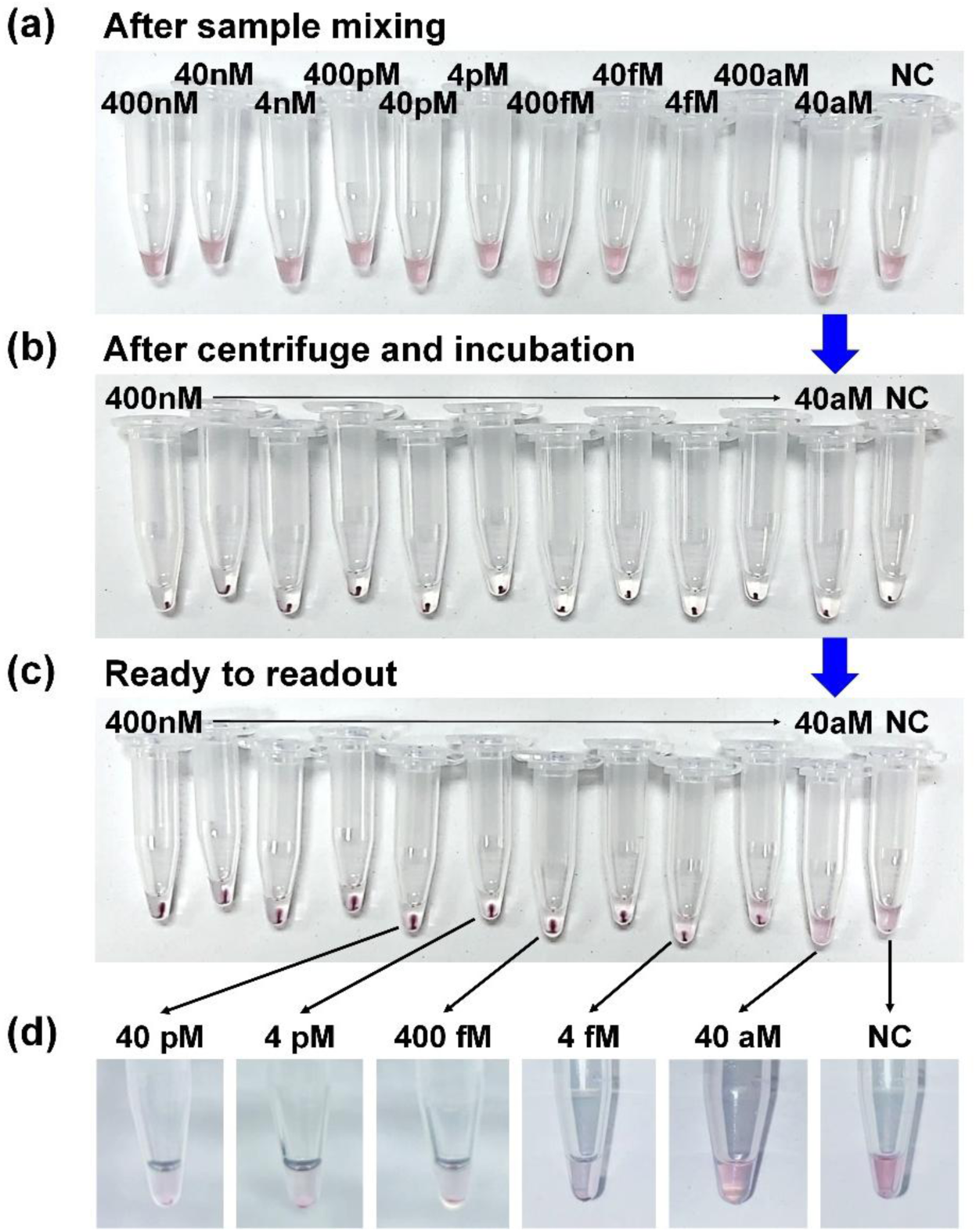
Optical images show samples in PBS buffer at each key analysis step. The testing concentration ranged from 400 nM to 40 aM. NC: PBS only. (a) After mixing the WT-RBD AuNP sensor solution and spiking AS35 in PBS buffer, (b) showing sedimentation of the sample after centrifugation (1,200 g, 5 minutes) and 5 minutes of incubation, (c) final sample ready for PED measurement after vortexing (34.5 rps, 5 seconds). (d) Magnified image of the microtubes of 40 pM, 4 pM, 400 fM, 4 fM, 40 aM, and NC samples, held vertically for PED measurements.

### 3.2 Effect of centrifugation on sensitivity enhancement

**Figure S4.**
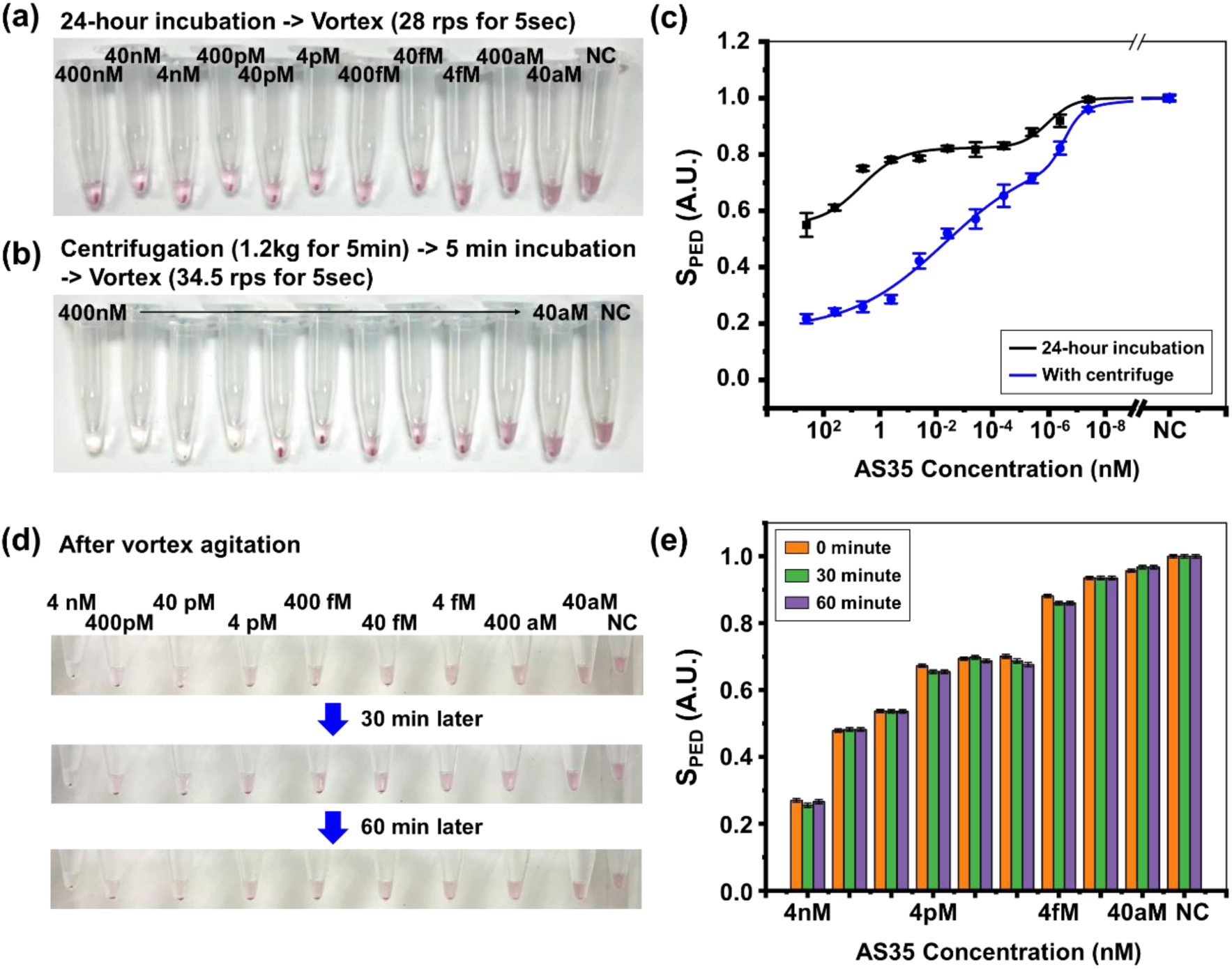
Comparison of centrifugation effects. (a-b) Optical of images AS35 samples ready for readout. (a) Samples were incubated for 24 hours without centrifugation and then vortex agitation was used for 5 seconds at 28 rps. When no centrifugation was used, the sample did not have sufficient time to make strong binding, so a low-speed vortex was used. (b) Samples centrifuged for 5 minutes at 1,200 g, incubated for 5 minutes, and vortexed for 5 seconds at 34.5 rps. (c) Extracted PED signals, according to whether centrifugation was used, show that the detection rate is significantly improved when centrifugation is used, especially in high-concentration conditions. (d–e) Stability of PED signal after the sensing protocol. (d) Optical images of AS35 samples immediately after vortex agitation (0 min), and after standing vertically for 30 and 60 minutes. (e) Quantified *S*_PED_ signals for samples at each time point. No significant signal change was observed over 60 minutes, indicating temporal stability of the readout.

### 3.3 TEM imaging of AuNP precipitates

**Figure S5.**
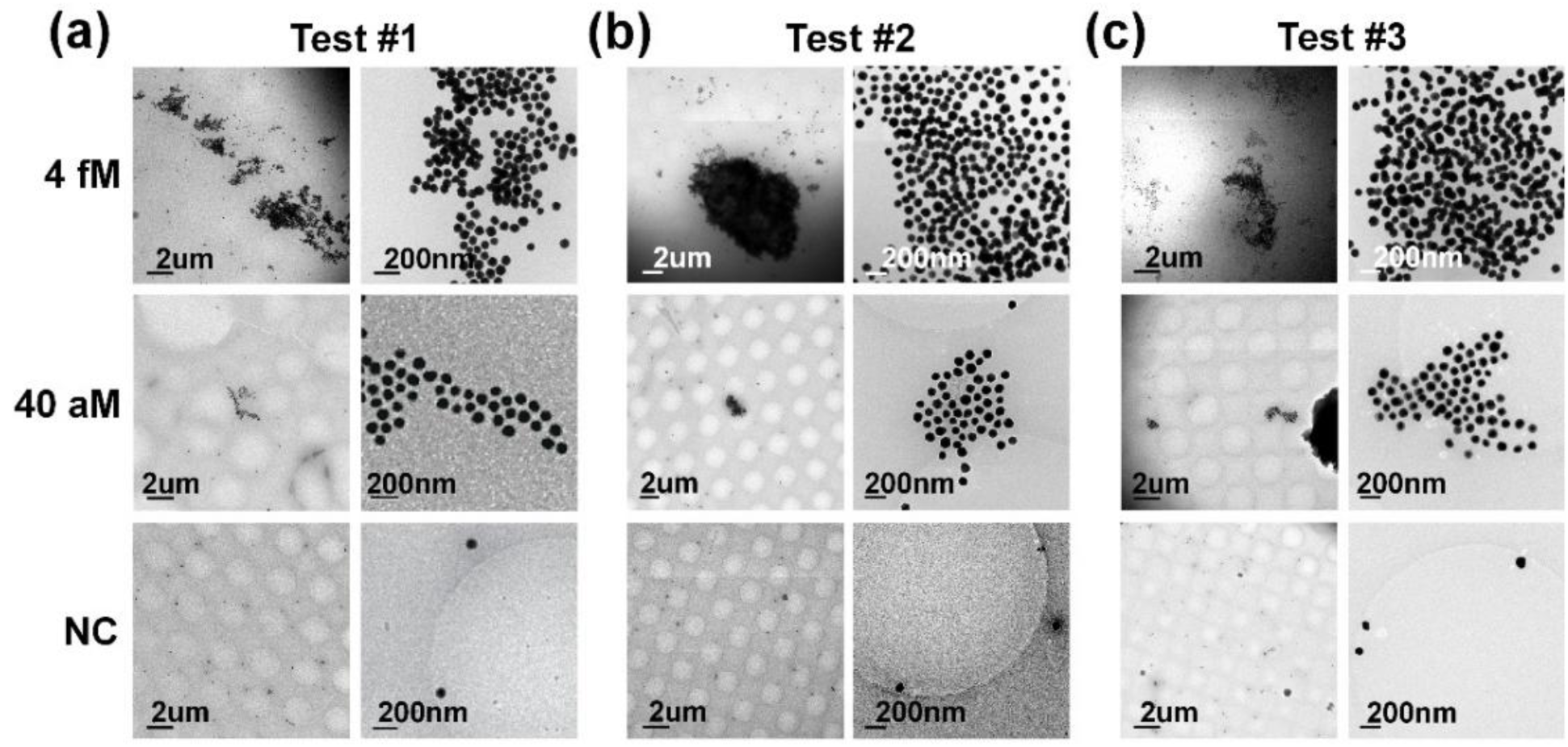
Microscopy analysis of AuNP precipitates. (a-c) Cryo-TEM images of 80 nm AuNP precipitate at the tube bottom in three replicate experiments at different AS35 concentrations (4 fM, 40 aM, and NC from top to bottom rows).

## 4. AuNP quantification analysis

### 4.1 AuNP concentration measurement

Using 80 nm streptavidin-functionalized AuNPs as a model, we attempted to determine the AuNP concentration *C_AuNP_* (#/mL) (Figure S6). The purchased AuNPs (Cytodiagnostics) had an initial optical density (OD) of 10 reported by the manufacturer. OD is a value that indicates the degree to which light is absorbed to measure concentration or quantity. Here, OD is defined as the decadic logarithm (base 10) of the ratio of incident to transmitted optical power, and a higher OD means a large extinction and a high AuNP concentration. However, a precise determination of their concentration *C_AuNP_* was not available. Here, we tried to use three different methods to estimate *C_AuNP_*, i.e. manufacturer estimate, NanoSight calibration, and PED signal conversion based on Beer–Lambert law.

#### 4.1.1 Manufacturer estimate

Taking into account the stock value (0.13 nM) and making volumetric and molar conversions using the same mixing ratio (18 μL out of 24 μL), we estimated it as

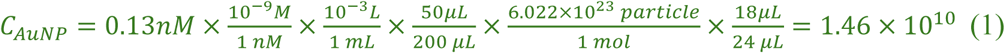

To adjust the final *OD*_*PED*_ to approximately 0.5 after two rounds of purification, we diluted the initially collected 50 μL AuNP solution into ∼200 μL sensing buffer. This dilution step was reflected in the volumetric conversion (50μL⁄200 μL) during the final particle number estimation.

#### 4.1.2 NanoSight calibration

For NanoSight (NS300, Malvern Panalytical) measurements, the as-purchased AuNP solutions were diluted three folds, and then serially diluted with a 1.5-fold dilution factor. Noticeably, a significant amount of AuNPs were still free-floating when the tubes started to turn transparent (e.g. *S*_*PED*_<0.1, or OD<∼0.2), and such AuNPs were sufficient to produce noticeable optical extinction and, therefore, PED signals, because of a long optical path between the LED and photodetector (∼5 mm).

To correlate the Nanosight signals with PED signals, 24 µL of AuNPs solutions at each concentration were added to each microcentrifuge tube and measured using NasRED.

Here, *C_AuNP_* was obtained from NanoSight and fitted to the manufacturer’s OD using a linear equation as follows:

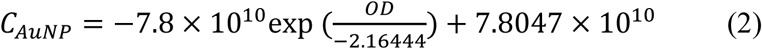

Further, the optical extinction signal obtained from NasRED readout O*D*_*PED*_ was also fitted to the manufacturer’s OD values with a linear relation as follows:

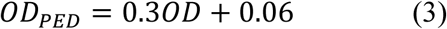

The above equations can be used to determine the *C_AuNP_* concentrations at the electronic signals of the NasRED using a simple linear relationship.

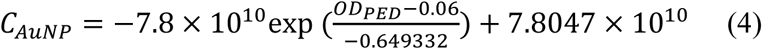

Using equation (3), the relationship can also be plotted (Fig. S5 e), which can be used to calibrate the AuNP numbers for each measured electronic signal intensity value.

Here, the average PBS buffer sample signals, 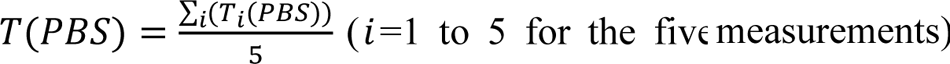, represented the optical transmission through each tube containing 24 µL of PBS buffer. Similarly, for the diluted AuNPs, five measurements were performed for each tube along the same orientations, and the electronic signals were again recorded as 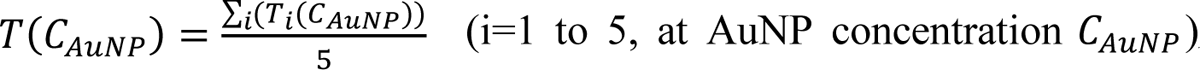. O*D*_*PED*_(*C_AuNP_*) is calculated to correlate the AuNP concentrations with the PED measured signals following:

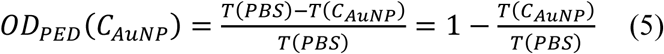

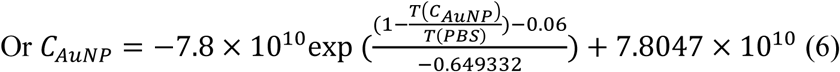

Based on the representative value for the case where O*D*_*PED*_ = 0.5, the following calculation was made:

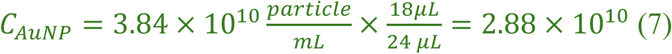

#### 4.1.3 PED measurement

Using the Beer–Lambert law, the absorbance *A* is defined by:

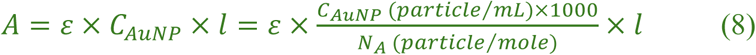

Where ε = 7.7 × 10^10^(M^−1^*cm*^−1^)^1^, *l* = 0.315*cm* is the optical path length, c_*AuNP*_ is the molar AuNP concentration, and N_*A*_ = 6.022 × 10^23^/mol is the Avogadro number.

Here, the absorbance *A* was calculated using the following conversion formula:

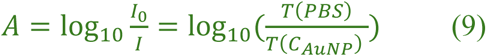

Rearranging and solving for *C_AuNP_*, and inserting the transmittance-based measurement:

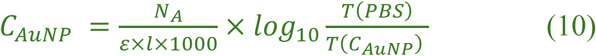

Using the measured value of *T*(PBS) ≅ 196 and *T*(*C_AuNP_*) ≅ 91, the final calculated AuNP concentration was:

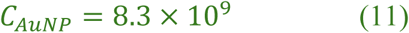

Notably, since the absorbance was measured from the final sensing solution containing both 18 μL of AuNPs and 6 μL of target (total 24 μL), the calculated concentration already reflects the actual mixed condition. Therefore, no additional volumetric correction factor (e.g., 18/24) was applied in this method.

The calculated AuNP concentrations (equations 1, 7, and 11) were on the same order of magnitude. The results were considered reasonably acceptable given the intrinsic errors in measurements, lending credence in our estimation of AuNP concentrations. For consistency, we chose the NanoSight-calibrated data (equation 7) to convert PED electronic signals to AuNP concentrations.

### 4.2 PED signal correlation

As detailed in the Method section, *S*_*PED*_ is a function of the tested protein marker concentration *c*, and normalized against the transmission signals *T*(*PC*) and *T*_0_(*NC*) from a positive control (PC) and negative control (NC) sample, following:

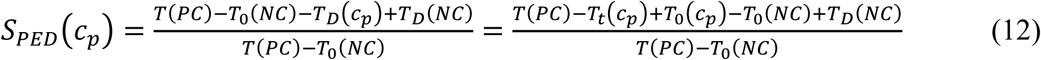

Here, *T*(*PC*) ≅ *T*(PBS) ≅ 196 (arbitrary units from PED readout) is the highest possible optical transmission for a given test in PBS, and *T*_0_*(c_p_)* = *T*_0_(*NC*) ≅ 91 is the transmission of the AuNP solution after mixing (thus irrelevant to the protein concentration *c_p_*), *T*_*t*_*(c_p_)* is the transmission of a possible protein sample after test, and *T*_*D*_(*NC*) ≅ 0 is the transmission signal difference after and before the sensing protocol for the negative control. Therefore, we can write:

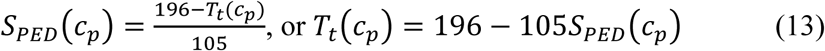

Effectively, in our experiment, we diluted the AuNP sensors to a target concentration of about OD∼0.5. To correlate the O*D*_*PED*_ with *S*_*PED*_, we should have *T*(*C_AuNP_*(*NC*)) = *T*_*t*_(*NC*), and *S*_*PED*_(*NC*) = 1. Mathematically using our experimental values for *T*_*t*_(*NC*) and *T*(*PC*), we can obtain 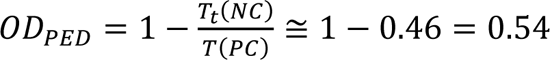. This is indeed close to the targeted 0.5 OD value.

On the other hand, when a particular diluted AuNP solution at concentration *C_AuNP_* displays the same color as a AuNP sensor solution after protein test at a protein concentration *c_p_*, we then have that *T*(*C_AuNP_(c_p_)*) = *T*_*t*_*(c_p_)*, and we can relate the AuNP concentration with the *S*_*PED*_ signal as:

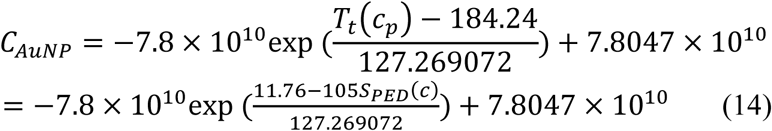

This further allows us to estimate the amount remaining free-floating AuNPs after a test from the *S*_*PED*_ signal. While the accuracy of AuNP concentration could be limited by the NanoSight precision and human errors in sample dilution, this analysis demonstrated that the AuNP concentration is linearly correlated with the *S*_*PED*_ values.

**Figure S6.**
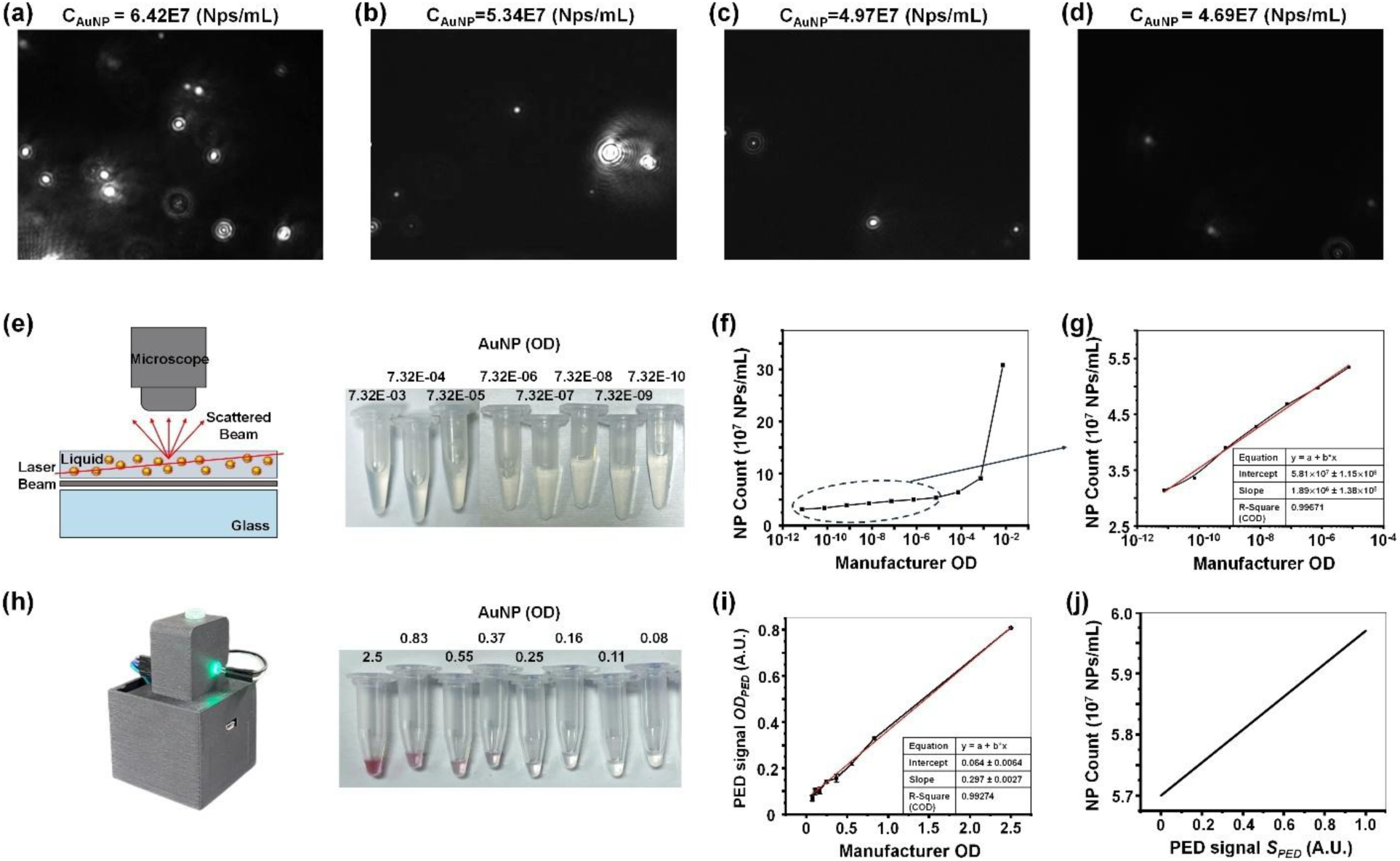
(a-d) Optical images of exemplary 80 nm streptavidin functionalized AuNPs using NanoSight. (a) AuNPs diluted 1000-fold from the initial stock concentration (10 OD), (b) AuNPs diluted 10-fold further from (a) to 7.32E-06 OD, (c) AuNPs diluted 10-fold further from (b) to 7.32E-07 OD, (d) AuNPs diluted 10-fold further from (c) to 7.32E-08 OD. A consistent decrease in the number of particles was observed after each dilution. (e-j) Analysis of nanoparticle concentrations. (e) Schematics of the Nanosight apparatus, where the pattern of the scattered beam through the solution was analyzed to yield the number of particles in liquid and tube pictures for the nanoparticle (NP) count. (f, g) The nanoparticle concentration measured by NanoSight plotted against manufacturer-provided OD values. (f) The NP plotted across the entire concentration range against manufacturer-provided OD values. At high concentrations, the proximity of AuNPs increases the laser scattering intensity, resulting in a rapid increase in NP count. (g) The NP count increased linearly with concentration and plotted using a linear fit from 7.32E-6 to 7.32E-10. (h) Tube pictures for the electronic readout signal *S*_*PED*_ showing the change of AuNP solution color at various dilutions correlated with the OD from the manufacturer. (i) The *S*_*PED*_ plotted against manufacturer-provided OD values. (j) Extracted nanoparticle concentration plotted against the normalized electronic readout signals *S*_*PED*_ from NasRED.

### 4.3 Protein interaction with AuNPs in NasRED

#### 4.3.1 Protein and AuNP diffusion

The diffusivities of AuNPs and antigens can be estimated from the Stokes−Einstein equation

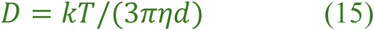

where *kT* is the thermal energy at room temperature (∼4.1 × 10^−21^ Joule at 300K), η is the solution viscosity (∼1.7 × 10^−3^ N · sec/m^2^ assuming 20% glycerol in water to estimate the buffer effect), and *d* is the diameter of AuNPs or proteins^2^. The diffusivity is estimated *D*_a_∼5.12 × 10^−11^ m^2^⁄s = 51.2 μm^2^⁄ *s* for a 5 nm protein and *D*_NP_∼3.2 × 10^−12^ m^2^⁄S = 3.2 μm^2^⁄S for an 80 nm AuNP.

We can further estimate the diffusion length L_a_, i.e. the distance for the analyte to collide with AuNPs, as the smaller of the inter-protein separation L_*p*_ and inter-AuNP separation L_NP_. L_NP_ was calculated following

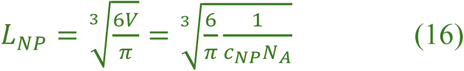

where *V* is volume estimated for each NP (assuming a sphere), *c* is the NP molar concentration, and N_*A*_ is the Avogadro number. Clearly, L_NP_ is determined by the NP concentration and thus a constant for a given assay design (for example, ∼2 µm for 80 nm AuNPs at 0.36 nM). Similarly, L_*p*_ depends on analyte concentration and can be calculated as ∼100 µm at a low antigen/antibody concentration (100 aM to 1 fM), ∼7 µm at 1 pM, but <1 µm at nM or higher concentration.

The thermal velocity of proteins can also be estimated by

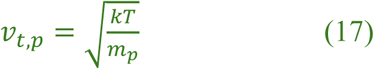

where M_*p*_ is the molecular weight (g/mol, e.g. ∼150,000 g/mol for an IgG antibody) and N_*A*_ = 6.022 × 10^23^/mol is the Avogadro number. Therefore, 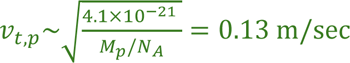 for an IgG antibody. Given the estimated micrometer sized protein to AuNP distance, the time required for a protein to collide with an AuNP is therefore estimated to be on the microsecond level, which is clearly not the rate-limiting factor of the NasRED assay. On the other hand, it is also useful to compare with conventional surface-incubation based assays, such as ELISA and SPR, where, according to the non-slip boundary condition, the surface velocity is close to zero. Therefore, the use of free-floating AuNPs in NasRED for reactions can significantly shorten the assay time.

#### 4.3.2 AuNP velocity, Peclet number, and Reynolds number

Given the small mass and inertia of AuNPs, the AuNPs almost instantaneously gain the speed set by the centrifuge or vortex. This can be evident from the small dynamic (Stokes) relaxation time of AuNP τ_*p*_, estimated by

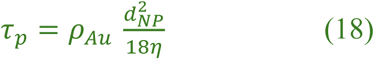

where ρ_*Au*_ is AuNP density (19.3 g/cm^3^) and η is dynamic viscosity of the fluid (∼1× 10⁻³ Pa·s). For 80 nm AuNPs, τ_*p*_ = 6.9 × 10^−9^S.

In a centrifuge, the centrifuge Relative Centrifugal Force (RCF) is defined as

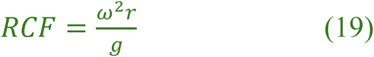

where ω is the angular velocity in radians per second (rad/s), and r is the distance from the rotation axis (m). The terminal velocity of AuNPs, following Stokes flow approximation, is estimated as. The 80 nm AuNP flow rate can be estimated from the centrifugal forces following

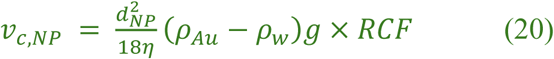

where ρ_w_ is the density of buffer^3^. Assuming a typical RCF=1200 during centrifugation in a typical NasRED process, we have v_*c*,NP_ = 6.38 × 10^−8^ × 1200 m/s = 7.6 × 10^−5^ m⁄S for 80 nm AuNPs.

On the other hand, AuNPs constantly diffuse. The dimensionless Peclet number (Pe) compares nanoparticle centrifugal sedimentation with Brownian motion following

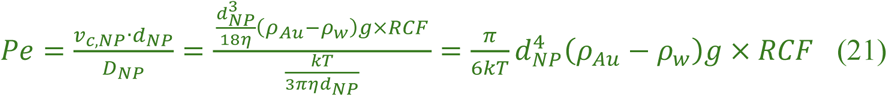

Here, 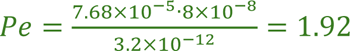 for the 80 nm AuNPs at *RCF* = 1200 *g*, indicating centrifugal force is stronger than Brownian motion. In comparison, for 20 nm, 40 nm, and 60 nm AuNPs, their Pe numbers would be ∼0.01, 0.12, and 0.61, respectively, suggesting dominance of Brownian diffusion over sedimentation. This is consistent with our previous observation that 80 nm AuNPs precipitate faster than smaller AuNPs^4^.

Furthermore, the AuNP cluster sizes increase with the protein analyte concentrations. In this work, our experimental results indicated that large clusters (e.g. in some cases up to ∼800 µm^2^, or >20 µm in diameter) formed at moderate AS35 concentrations (e.g., 40 pM, 4 pM, and 400 fM), and smaller clusters (<1 µm in diameter) were found at lower (fM or less) antibody concentrations (Figure 3). While the clusters are not exactly spherical in shape and their Peclet numbers are difficult to precisely determine, it is still reasonable to assume that these AuNP clusters can have Pe > 1 and tend to precipitate much faster than single AuNPs.

Additionally, the dimensionless particle Reynolds Number (*Re*_*p*_), representing the relative strengths of inertial forces over viscous forces, can be calculated as

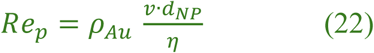

As an example, *Re*_*p*_∼0.04 for 80 nm AuNPs when Brownian motion dominates. In comparison, AuNP clusters at micrometer size could have *Re*_*p*_>1, in other words, inertial forces can play an important role in their fluidic dynamic behaviors.

#### 4.3.3 Enhanced protein binding on NasRED AuNP sensors

##### Local concentration enhancement

Using 80 nm AuNP as an example and assuming protein sizes of 5 nm, we can estimate that each of the 80 nm AuNPs has up to ∼500 antibodies or antigens on their surface. In the NasRED assay, the AuNP sensors had a concentration of 0.048 nM (calibrated by Nanosight, see supplementary section 4.1). In other words, the total number of AuNPs in each test (24 μL liquid) was 6.92 × 10^8^. This also suggested an initial ligand (antibody or antigen) concentration of as high as 24 nM in the test tube. After centrifugation, the AuNPs are pelleted to the bottom of the tube, thus greatly enhancing their concentration. Assuming dense packing of AuNPs, all the nanoparticles would condense into a volume of ∼3.5 × 10^−4^ μL. This corresponds to 67,000 times volume reduction and thus 67,000 times increase in the ligand concentration. Such greatly enhanced, localized antibody or antigen concentration would significantly promote the binding of target analytes.

##### Avidity effect

Functional affinity enhancement (as large as ∼10^6^), e.g. avidity effect^5^, was discovered decades ago using multivalent binding of the pentameric IgM antibody to the phage involving three or more of its combining sites. Avidity has also been demonstrated by creating higher order multimers of nanobodies against parathyroid hormone (PTH), showing an affinity gain of three to four orders of magnitude^6^. This was accompanied by a corresponding decrease in the constant dissociation rate, from over 1 s^-1^ to an estimated 0.0001 s^-1^. In the case of NasRED, considering the multivalency of AuNP sensors, it is reasonable to believe that the dissociation process of bound protein from AuNP sensors is greatly suppressed, leading to greatly improved binding affinity. Indeed, previously using Smoluchowski’s coagulation modeling, we estimated the probability of aggregation per collision was approximately 1, meaning every collision between protein analyte and functionalized AuNPs will result in protein binding^4^.

##### Dynamic protein binding

In NasRED, the target analyte (antigen or antibody) binding occurs on actively diffusing AuNPs. This is fundamentally different from binding to stationary sensor surfaces in ELISA or SPR, where the flow is based on passive diffusion. As a result, the k_on_ and k_off_ values, measured from ELISA or SPR, describe the surface-bound molecular interactions but cannot fully predict what happens at the nanometer scale in NasRED. This has been observed that the binding kinetics in solution could be significantly different from that on the surface^7^, attributed to mass transport and other factors^8^. Given the high affinity of protein-antibody pairs in this work, we can reasonably hypothesize that the association process of this complex is fast (e.g., G protein binds to GPCR receptors within ∼0.3 sec^9^).

#### 4.3.4 AuNP sedimentation

In the case of sensing by precipitation without introducing centrifugation or vortex agitation, the gravitational force of AuNPs competes with their fluidic drag forces to determine the sedimentation process. The change rate of AuNP concentration c is determined by the AuNP diffusion and sedimentation, following the Mason–Weaver Equation 10

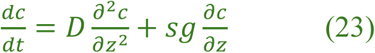

where again *D* = *kT*/(3πηd) is the diffusion coefficient (*d* is the AuNP and aggregate diameter, η is the dynamic viscosity of the buffer), *g* is the gravitation constant, and *s* is the sedimentation coefficient, defined by

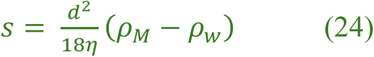

Where ρ_M_ and ρ_w_ are the density of aggregate and buffer, respectively^3^. For AuNPs of 80 nm, S = 6.5 × 10^−9^(s).

The sedimentation time can also be estimated using the Mason-Weaver equation by

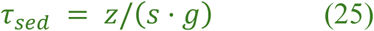

where *z* is the precipitation path (the height of the colloid liquid). Given *z* ∼6 mm for 24 μL liquid in a microcentrifuge tube, we can estimate that τ_Sed_ decreases from 22 hours for an 80 nm AuNP monomer to 8 minutes and 5 seconds for 1 µm and 10 µm diameter clusters, respectively.

With the introduction of centrifugation, most (if not all) of the AuNPs are pelleted at the bottom of the centrifuge tube, forming a dense layer estimated at the micrometer scale in thickness. To estimate the volume of AuNPs accumulating in the tube, full agglomeration of nanoparticles is assumed without complete coalescence, either with each other or with the tube walls. Assuming each AuNP is spherical and the effective protein-coated AuNP diameter as 90 nm, the volume occupied by a single protein-coated AuNP will be:

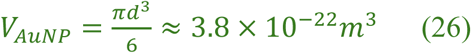

The total volume occupied by all the AuNPs supplied in the sensing tube is calculated considering the atomic packing factor (APF) of 0.74 (Face-Centered Cubic).

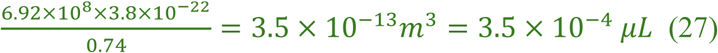

Where N is the number of nanoparticles per tube. In practice, BSA and target molecules may occupy interstitial spaces between AuNPs, and the biological solution and associated electrostatic or chemical reactions may affect the packing density. Nevertheless, considering closely packed AuNPs precipitated at the cone-shaped tube bottom (with the apex angle θ∼20°), we have

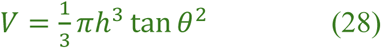

Where h∼136 μm is the AuNP precipitate thickness, tens of times smaller than the original solution height (∼6 mm). Clearly, this condensation process eliminates the AuNP solution sedimentation time and thus speeds up redout.

#### 4.3.5 AuNP concentration profile

From Mason–Weaver Equation^10^ and at equilibrium or quasi-equilibrium condition (dc/dt ∼0), we have

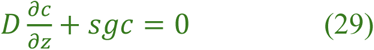

The solution to the above equation converges to an exponential decay function, which has been experimentally verified for AuNP sedimentation^3^:

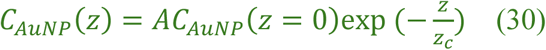

Where *A* is a constant, and the characteristic height of the equilibrium gradient is

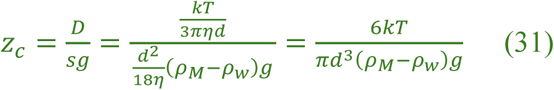

For AuNPs of 80 nm, 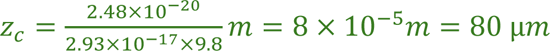.

In the case of NasRED, AuNPs are resuspended by vortex agitation (after centrifugation and incubation). Different from the above, where the gradient profile is created by gravity-induced sedimentation, active fluidic forces during vortex agitation can affect the AuNP motion. In fact, the fluidic dynamic process for AuNPs to gain speed and lift up is quite complex, involving both shear and spin-related fluidic drag forces, which affects but also changes with the AuNP flow speed, rotation, and positions. Considering the protein-concentration-dependent AuNP cluster size distribution, inhomogeneous mixture of AuNP monomers and clusters in the precipitates, and the conical geometry of the microcentrifuge tubes, the AuNP monomers’ motion and distribution are expected to be rather complex. Therefore, to theoretically model the AuNP concentration profile after vortex agitation, we explore the use of the exponential distribution function derived from the Mason-Weaver Equation, but take into account the complex impacts of fluidic dynamic lift force exerted on the AuNP monomers and clusters. Here we replace this characteristic height with Z_c_*(c_p_)* as a protein-concentration-dependent decay length, and rewrite the AuNP distribution as follows:

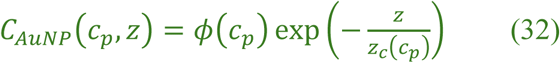

Where ø*(c_p_)* = *C_AuNP_*(*c_p_*, *z* = 0) is an empirical fitting parameter characterizing the AuNP concentration at tube bottom (z=0).

In the NasRED assay, the aggregated AuNP clusters would not contribute to the AuNP monomer distribution. While it is challenging to precisely determine the amount of AuNPs in the clusters, here we make an estimate using the following assumptions. (1) At low concentrations (e.g. at 40 aM level or lower), AuNPs are much more concentrated than protein analytes, and therefore each protein-AuNP may have only one protein attachment, i.e. the number of proteins per AuNP γ(40aM) = 1. (2) At high concentrations (e.g. at nanomolar range), the binding sites on the AuNPs become saturated, and for simplicity we can estimate γ(40*nM*) ≅ 200 (less than half of total estimated 500 binding sites considering steric hindrance in binding). This is based on the observation that the AuNP extinction signals were minimal at high protein concentrations. (3) In the intermediate protein concentration ranges, the number of proteins per AuNP scales in a logarithmic fashion

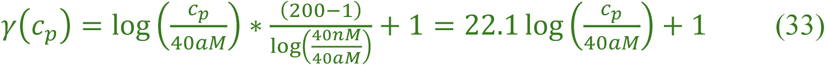

Therefore, the amount of AuNPs (per cross-sectional area) within the precipitates at the bottom of the tube is expressed as:

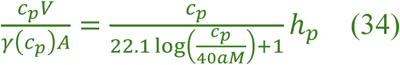

Where V is the initial volume of samples to be analyzed (6 µL), A is the average cross-sectional area of the tube (∼10 mm^2^), leading to ℎ_*p*_ = V⁄*A* ∼0.06 mm.

Using these assumptions and following mass conservation, we can model the remaining AuNP monomers in a simplified one-dimensional distribution:

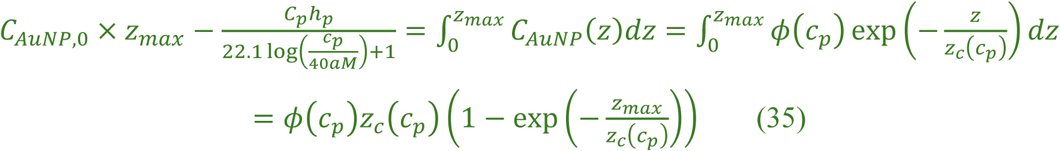

Where *C*_*AuNP*,0_ = 18/24 × 3.8 × 10^10^ #/mL=4.7 × 10^−11^M =47 pM is the initial AuNP concentration (diluted by a factor of 18/24 after mixing with the 6 uL testing sample) in the sensing tube prior to centrifugation or vortex agitation, and *z*_max_ = 6 mm is the solution height.

#### 4.3.6 Estimation of AuNP concentrations at different solution heights

To validate our hypothesis of the AuNP concentration profile, we customized a new tube holder with newly designed circuits to measure the PED signal at two different levels (*z*_1_ = 5.5 mm and *z*_2_ = 3.5 mm) in reference to the tube bottom (z=0) (Figure S13a). The PED signals were measured at each location for various concentrations of AS35 antibody (400 pM – 40 aM, and NC) using the WTRBD-functionalized AuNP sensors, consistently showing higher signals at the lower location, consistent with our model (Figure S13b-c). Therefore, the AuNP concentrations should follow:

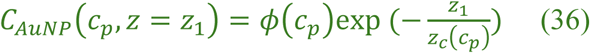

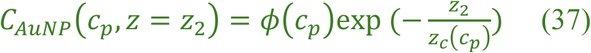

Following Beer–Lambert law, the estimated AuNPs concentrations and their signal variation (SD, or σ) can be calculated as follows:

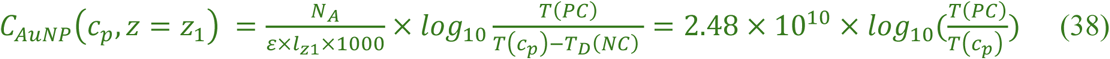

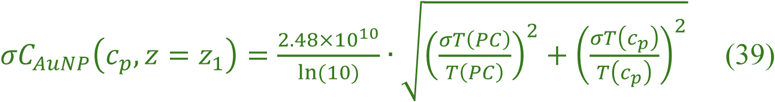

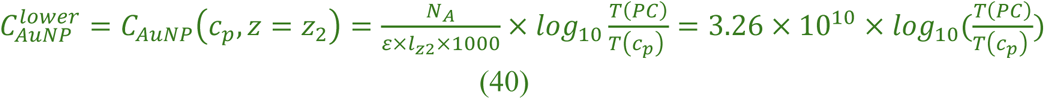

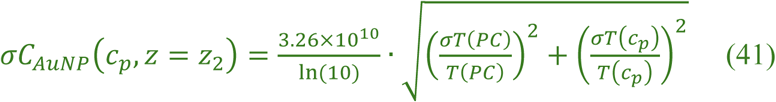

Where the optical paths *l*_*z*1_ = 0.315*cm* and *l*_*z*2_ = 0.24*cm* correspond to vertical positions *z*_1_ and *z*_2_, respectively.

With the above equations, the AuNP concentration profile could be calculated following an exponential function model and applying mass conservation (equation 35). Using the scipy library in Python^3^ (Appendix).

**Figure S7.**
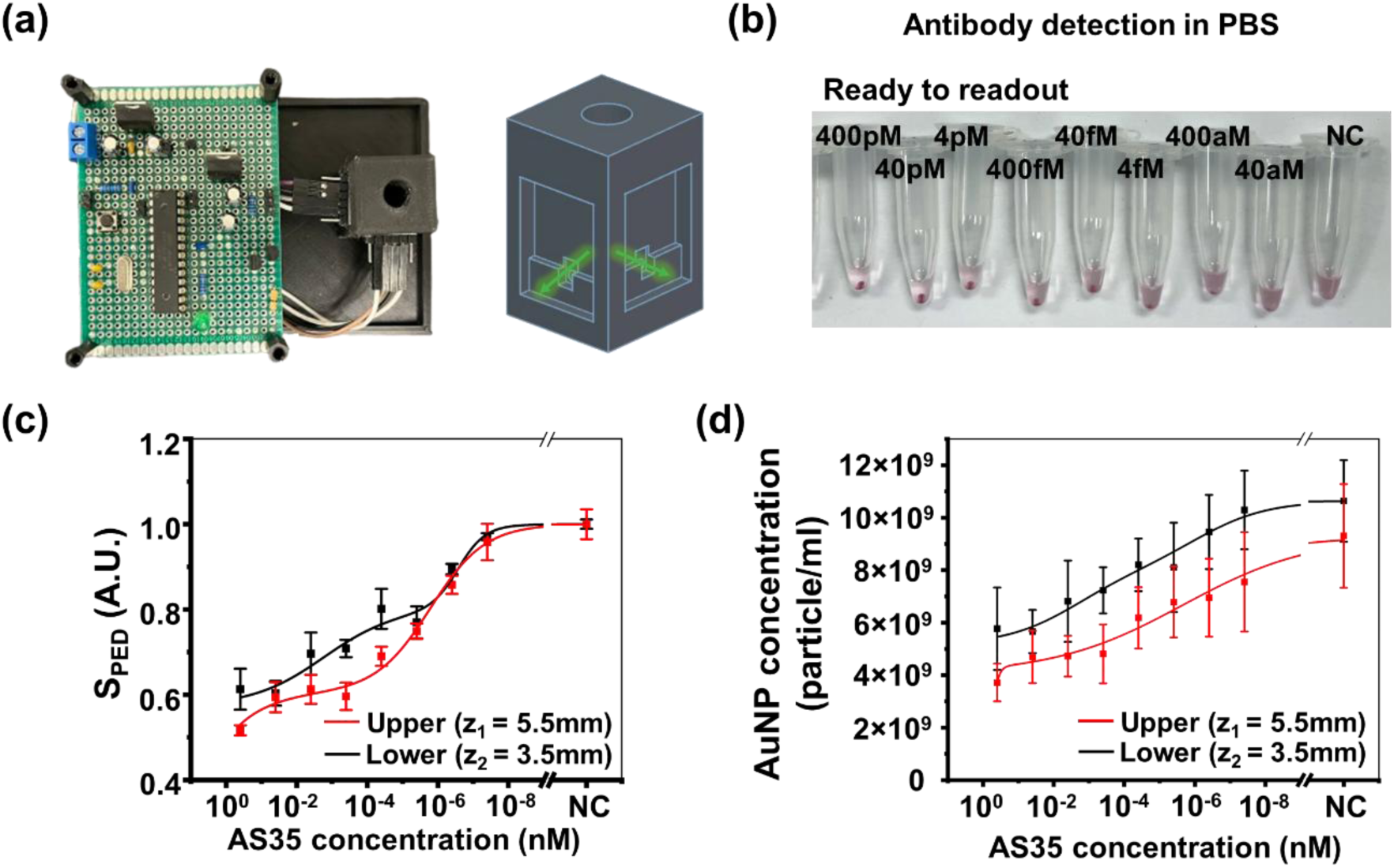
(a) Schematic configuration of the custom-built device and tube holder enabling dual-position PED signal readout at the upper (5.5 mm) and lower (3.5 mm) regions of the tube. (b) Optical image showing antibody samples ready for NasRED readout. (c) Extracted PED signals measured at the upper and lower positions across varying AS35 concentrations; the lower position consistently yielded stronger signals, indicating nanoparticle accumulation after vortex agitation. (d) AuNP concentrations at both positions were calculated using the Beer–Lambert law. Protocol parameter: centrifugation at 1,200 g for 5 minutes, incubation for 5 minutes, and vortex agitation at 34.5 rps for 5 seconds.

## 5. Sensing optimizations of SARS-CoV-2 antibody and antigen

### 5.1 ELISA test of SARS-CoV-2 antibody detection in PBS

**Figure S8.**
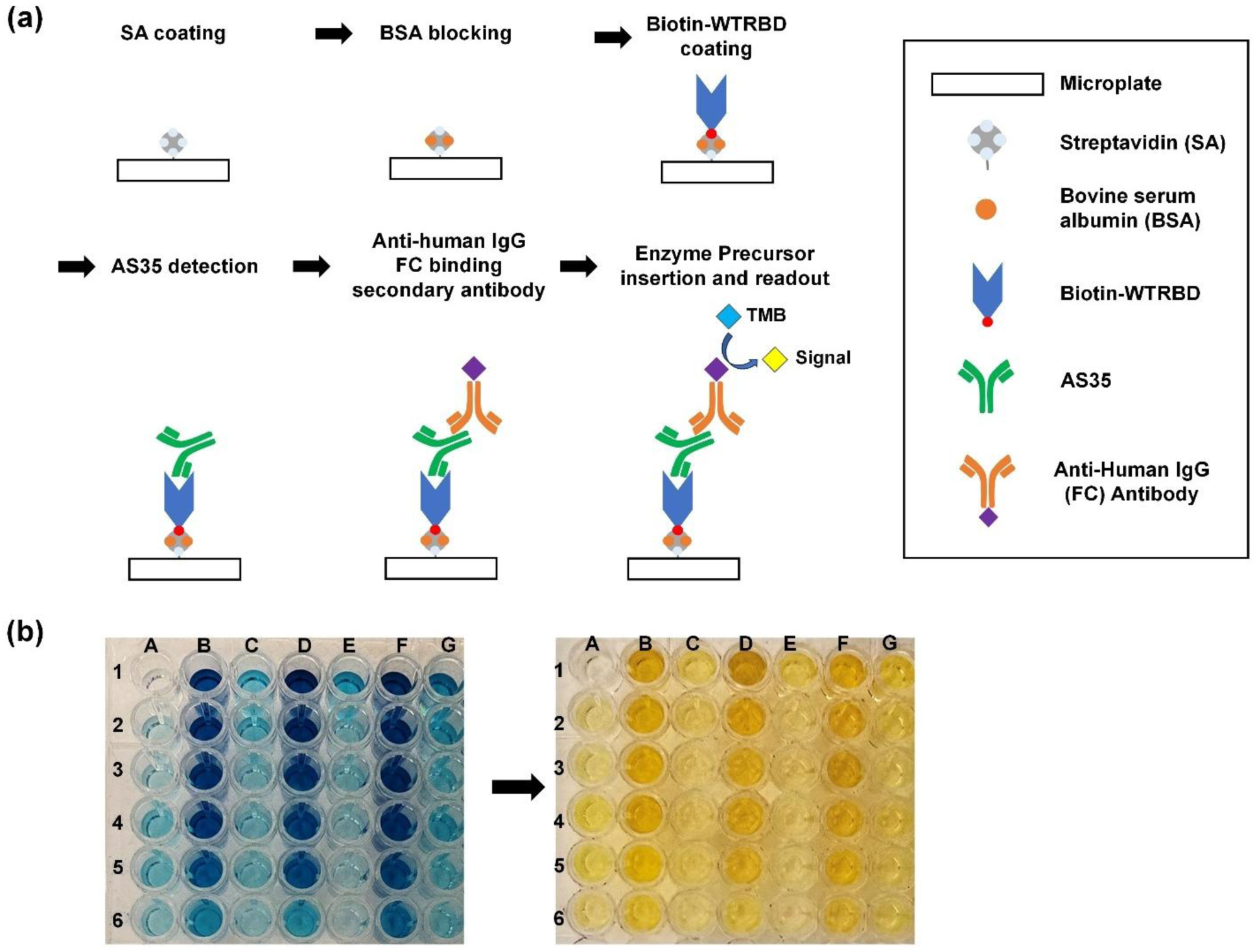
Schematic illustration of the ELISA procedure. (a) The process begins with streptavidin coating on a microplate, followed by BSA blocking to minimize non-specific binding. Biotin-WT-RBD is then coated onto the microplate, enabling AS35 detection—the addition of an anti-human IgG FC-binding secondary antibody to amplify the signal. (b) The addition of TMB substrate produces a detectable signal (i) after 15 minutes of adding TMB solution and (ii) after stopping the reaction with H_2_SO_4_

**Table S2.** ELISA plate sample information.

|  | A | B | C | D | E | F | G |
| --- | --- | --- | --- | --- | --- | --- | --- |
| 1 | Blank | 400 nM | 400 fM | 400 nM | 400 fM | 400 nM | 400 fM |
| 2 | SA | 40 nM | 40 fM | 40 nM | 40 fM | 40 nM | 40 fM |
| 3 | SA+BSA | 4 nM | 4 fM | 4 nM | 4 fM | 4 nM | 4 fM |
| 4 | SA+BSA+RBD | 400 pM | 400 aM | 400 pM | 400 aM | 400 pM | 400 aM |
| 5 | SA+BSA+RBD | 40 pM | 40 aM | 40 pM | 40 aM | 40 pM | 40 aM |
| 6 | SA+BSA+RBD | 4 pM | 4 aM | 4 pM | 4 aM | 4 pM | 4 aM |

### 5.2 Reproducibility test of SARS-CoV-2 antibody detection in PBS

**Figure S9.**
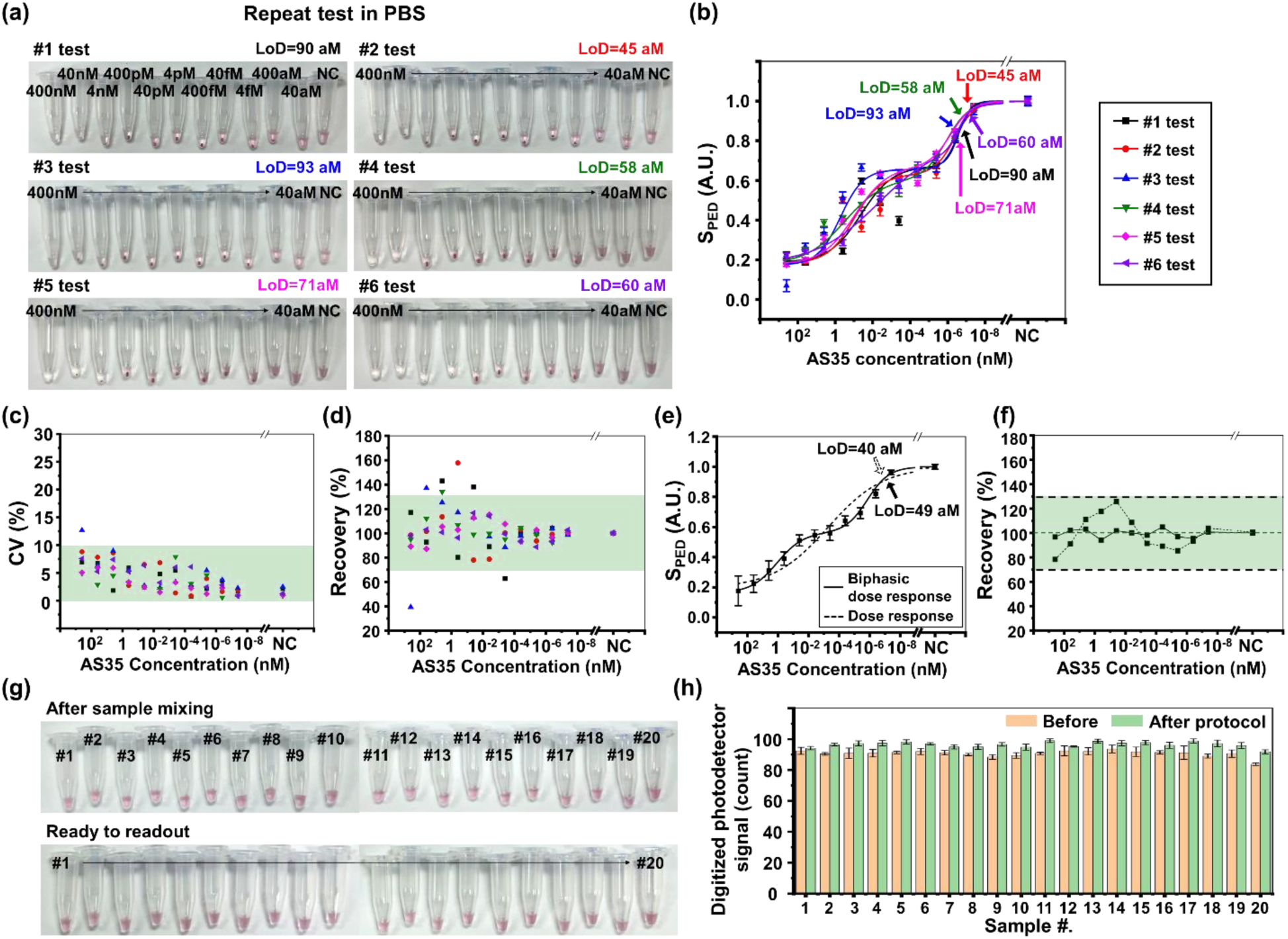
Reproducibility test of antibody in PBS. (a) Optical images of microcentrifuge tubes from 6 individual replicates ready for readout, (b) Extracted PED signals for the six replicates, producing LoDs of 90, 45, 93, 58, 71, and 60 aM, respectively. (c) Coefficient variation (CV) from 6 replicates in PBS, revealing a small coefficient variation (CV) (<10%, green shadow) of *S*_*PED*_. (d) Recovery (%) using Biphasic dose response fitting data from six replicate experiments, with the desired value range (100±30%) highlighted by green shadow, showing higher stability (100±5%) at low concentrations (40 fM - 40 aM). (e) Extracted PED signals fitted with sigmoidal (dash line, LoD ∼40 aM) and biphasic models (solid line, LoD ∼49 aM). (f) The recovery derived from sigmoidal (dash line) and biphasic (solid line), showing the biphasic fitting produced closer to 100% recovery at moderate and low concentrations and thus was more accurate. (g-h) Reproducibility test of 20 NC (blank) samples in PBS: (g) Optical images of microcentrifuge tubes in 20 NC samples after mixing the AuNP sensor solution with PBS buffer (top row) and the same samples after completion of sensing protocol (centrifugation at 1,200 g for 5 minutes, incubation for 5 minutes, and vortex agitation at 34.5 rps for 5 seconds). (h) Extracted photodetector signals plotted against each NC sample. NC samples were redispersed to their original state with less than a 6% difference before and after protocol. Protocol parameter: centrifugation at 1,200 g for 5 minutes, incubation for 5 minutes, and vortex agitation at 34.5 rps for 5 seconds.

### 5.3 Reproducibility test of SARS-CoV-2 antibody detection in human pooled serum

**Figure S10.**
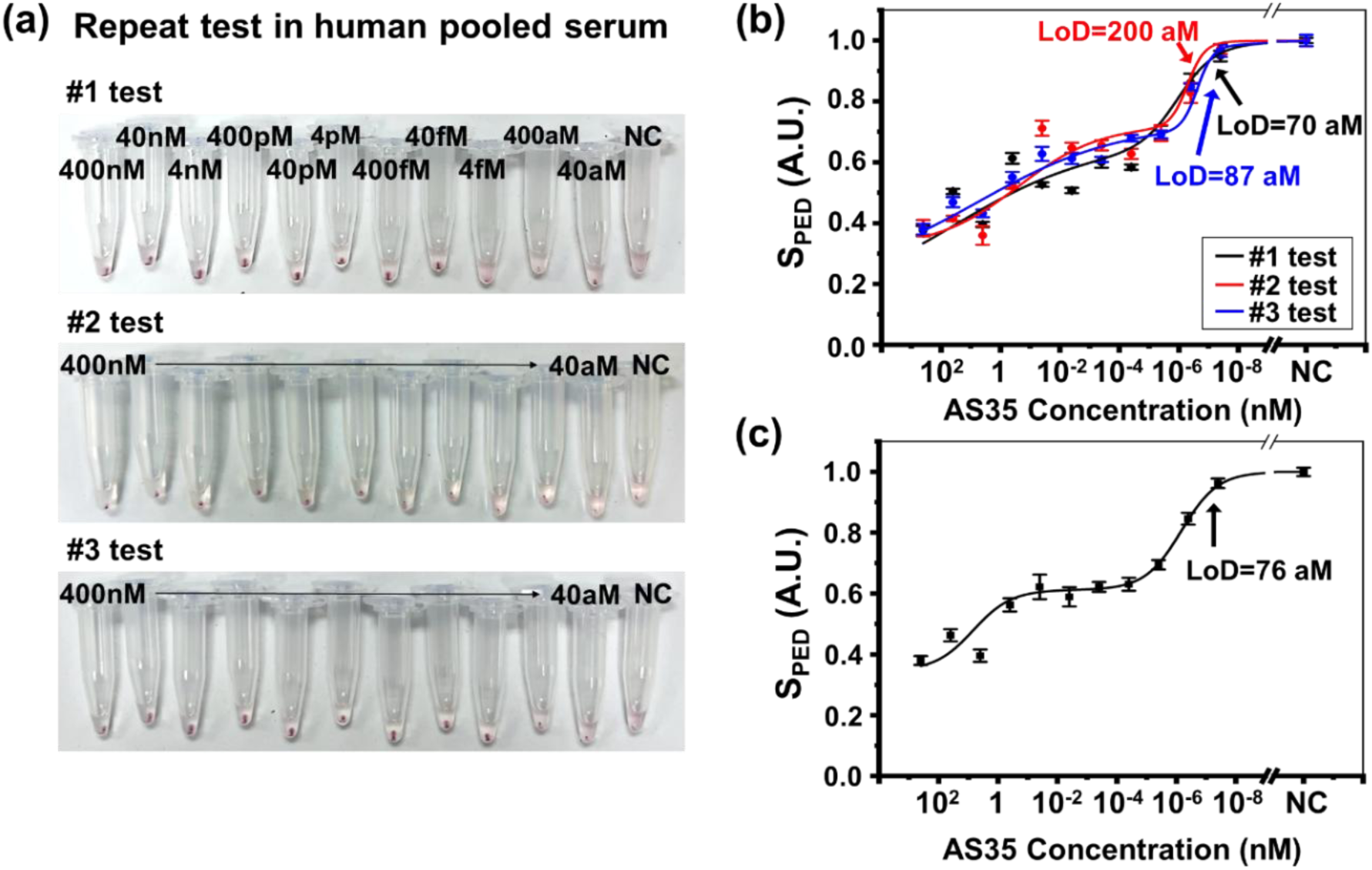
Reproducibility test of SARS-CoV-2 antibody AS35 in 100% human pooled serum. (a) Optical images of microcentrifuge tubes from 3 individual replicates ready for readout. (b) Extracted PED signals and Biphasic dose response fitted responses yielding a LoD of 70, 87, and 200 aM, respectively. (h) Averaged signals of the 3 replicate experiments in HPS with a LoD of 76 aM. Protocol parameter: centrifugation at 1,200 g for 5 minutes, incubation for 5 minutes, and vortex agitation at 34.5 rps for 5 seconds.

### 5.4 Antibody sensing in 1% diluted blood samples

**Figure S11.**
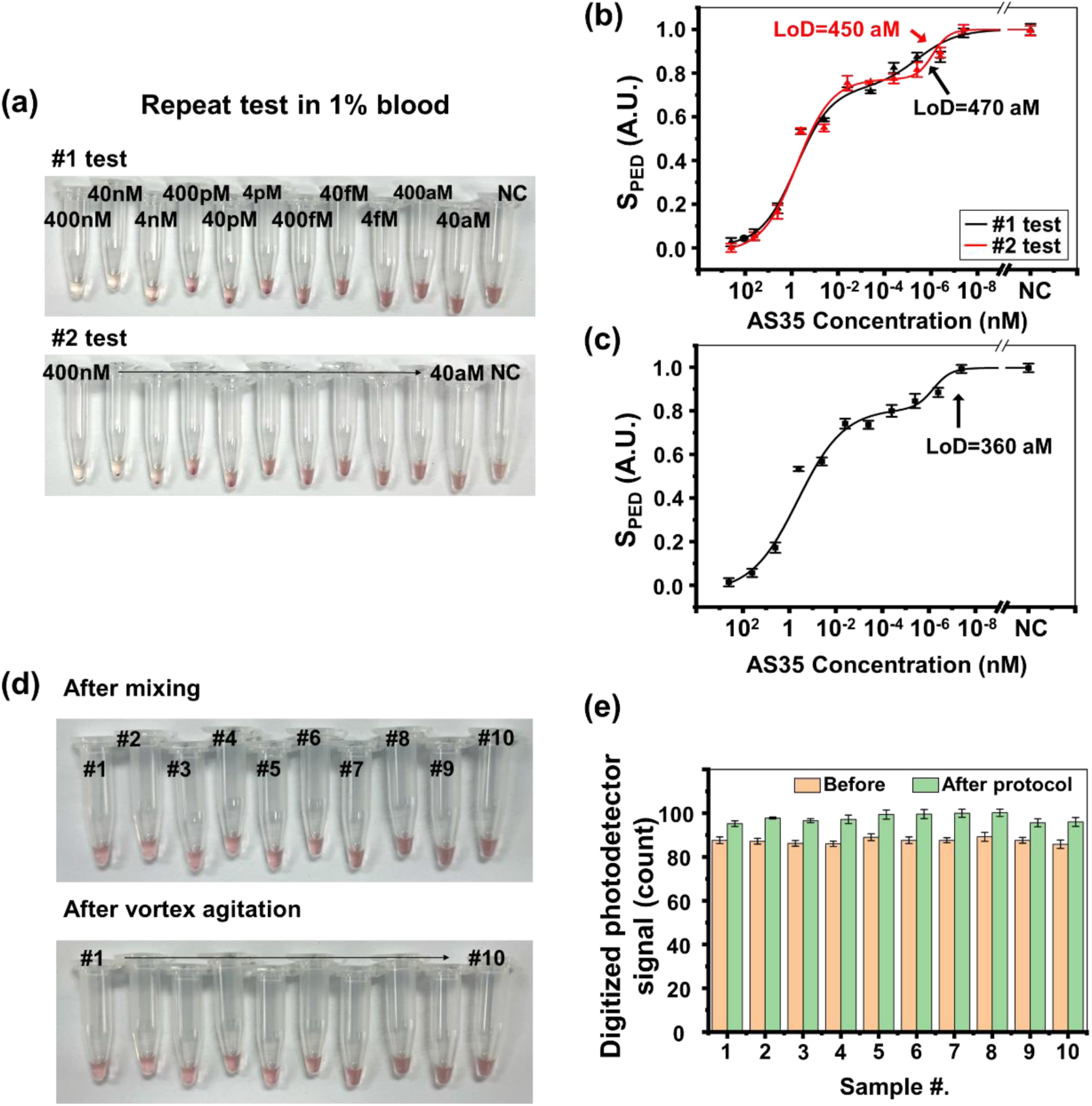
Antibody AS35 detection in 1% human whole blood (WB). (a) Optical images of microcentrifuge tubes from 2 individual replicates ready for readout. (b) Extracted PED signals and Biphasic dose response fitted responses yielding a LoD of 470 and 450 aM, respectively. (h) Averaged signals of the 2 replicate experiments in 1% WB, showing a LoD of 360 aM. (d) Reproducibility test of 10 NC (blank) samples in 1% WB. (a) Optical images of microcentrifuge tubes in 10 NC samples after mixing the AuNP sensor solution with 1% WB (top row) and the same samples after completion of our defined sensing protocol. (e) Extracted photodetector signals plotted against each NC sample in 1% WB. NC samples were redispersed to their original state with less than a 10% difference before and after protocol. Protocol parameter: centrifugation at 1,500 g for 5 minutes, incubation for 20 minutes, and vortex agitation at 37.5 rps for 5 seconds.

### 5.5 SARS-CoV-2 antibody detection in 20% whole blood samples

Hemoglobin in whole blood exhibits two characteristic absorbance peaks, one near 540 nm due to heme–heme interactions and the other near 578 nm related to oxygen binding affinity. These two peaks artificially produce an absorbance minimum in the intermediate region (550–575 nm) of the spectra. The 80 nm AuNPs exhibit a surface plasmon resonance (SPR) peak near ∼560 nm. The amount of free-floating AuNPs after sensing is dependent on the target protein concentrations, thus producing analyte-dependent extinction intensities as shown from our previous work (Chen et al., 2022). In the blood matrix, the AuNPs modulate the measured extinction spectra of the blood matrix, resulting in highest extinction for the NC sample but lowest for the high-concentration protein samples. The observations are consistent with our previous observations in Ebola antigen sensing (Chen et al., 2022).

**Figure S12.**
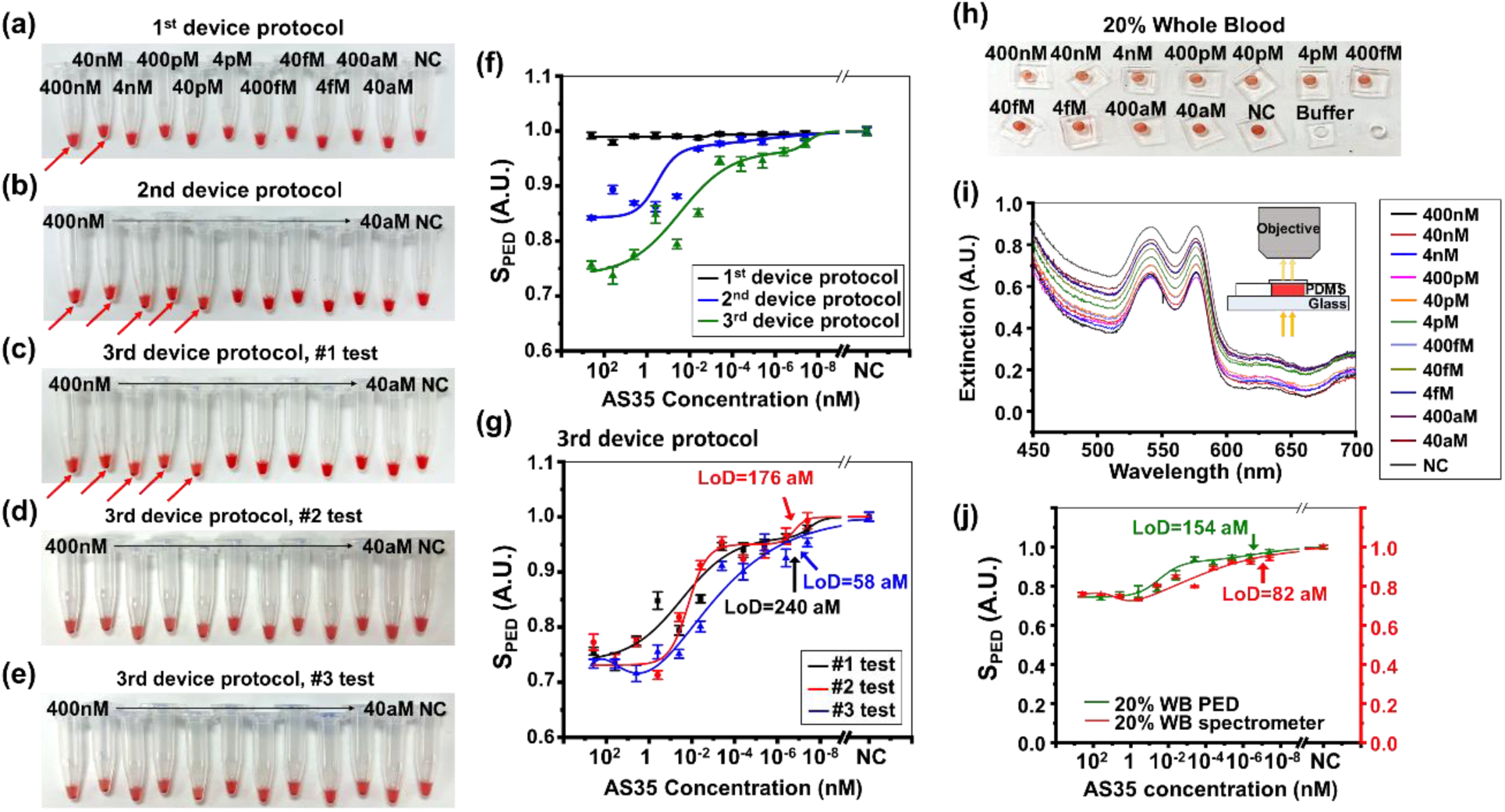
SARS-CoV-2 antibody detection in 20% diluted human whole blood (WB). (a-e) Optical images showing 20% WB samples ready for NasRED readout. (a-c): More visible AuNP clusters were observed at the tube bottoms under protocols #2 and #3. (c-e): Optical images of testing tubes in 3 replicates. (f) Extracted PED signals plotted against antibody concentration using the three sensing protocols. (g) PED signals and Biphasic dose fitting for replicate tests using protocol #3, yielding a LoD of 240, 176, and 58 aM, respectively, producing an averaged signals of a LoD in 20% WB of 160 aM (or 800 aM in 100% WB). (h-i) Optical images, spectra and schematic showing supernatant blood samples loaded into a PDMS plate for spectrometer measurement of the optical extinction. Spiked antibody concentrations: NC, 40 aM-400 nM. (j) PED signals (green line, LoD=154 aM, or 0.023 pg/mL) and spectrometer signals (red line, LoD=82 aM) were plotted against antibody concentration in 20% WB. Parameters of different sensing protocols are shown in Table S3.

**Table S3.**
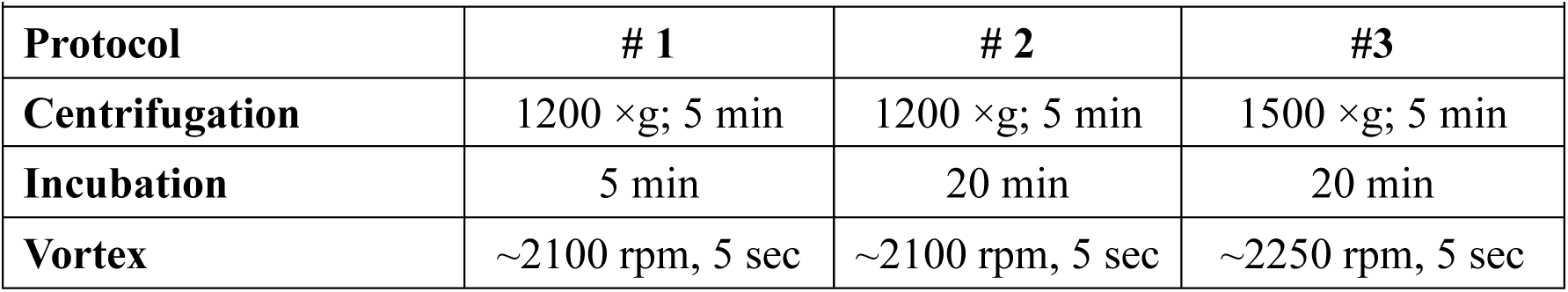
Parameters in WB sensing protocols.

### 5.6 SARS-CoV-2 antibody detection in sera and blood

**Figure S13.**
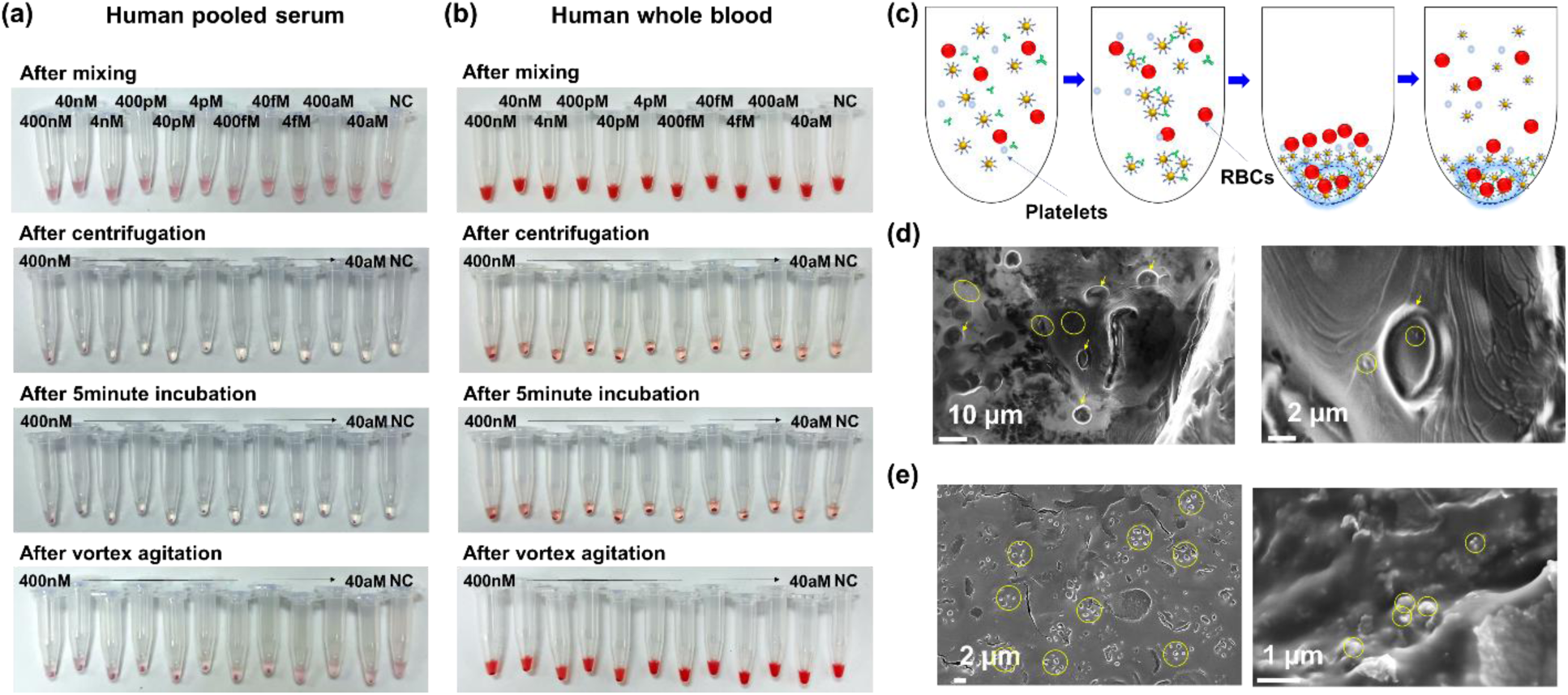
Extended data of microcentrifuge tubes used for SARS-CoV-2 antibody detection. (a-b) Images of centrifuge tubes for AS35 antibody detection in: (a) human pooled serum samples, (a) 20% human whole blood (WB). Immediately after mixing the sensor solutions in all tubes had identical and homogenous colors. Then, functionalized AuNPs precipitated after the centrifugation and incubation in the sera samples, while both AuNPs and blood cells precipitated in blood. After vortex agitation, the unbound free-floating AuNPs and blood cells were dispersed for both sera and WB samples, but the AuNP clusters stayed at the tube bottom. The clusters could also have included blood cells and platelets co-precipitated with AuNPs at the bottom of the tube. (c) Conceptual representation of the co-precipitation mechanism. (d-e) SEM images of the NC sample in blood matrix. (d) (left) NC samples showing blood cell (yellow arrows) and AuNPs (yellow circles) distributed on the imaging grid. (right) Zoomed-in image showing a red blood cell with AuNPs, indicating the possibility of AuNP uptake. (e) The sample with 40 fM antibody concentration, showing the AuNPs in the biological matrix, with a possibility that the AuNPs caused lysis of the blood cells. These SEM results indicated the complexity of AuNP interactions with blood cells during centrifugation, incubation, and vortex agitation, which could include cell uptake of individual AuNPs, AuNP cluster formation at the presence of complex biological matrix, cell lysis, etc. Future theoretical and empirical studies are needed to understand the mechanisms comprehensively.

### 5.7 SARS-CoV-2 antigen detection in PBS, saliva, and nasal fluid

**Figure S14.**
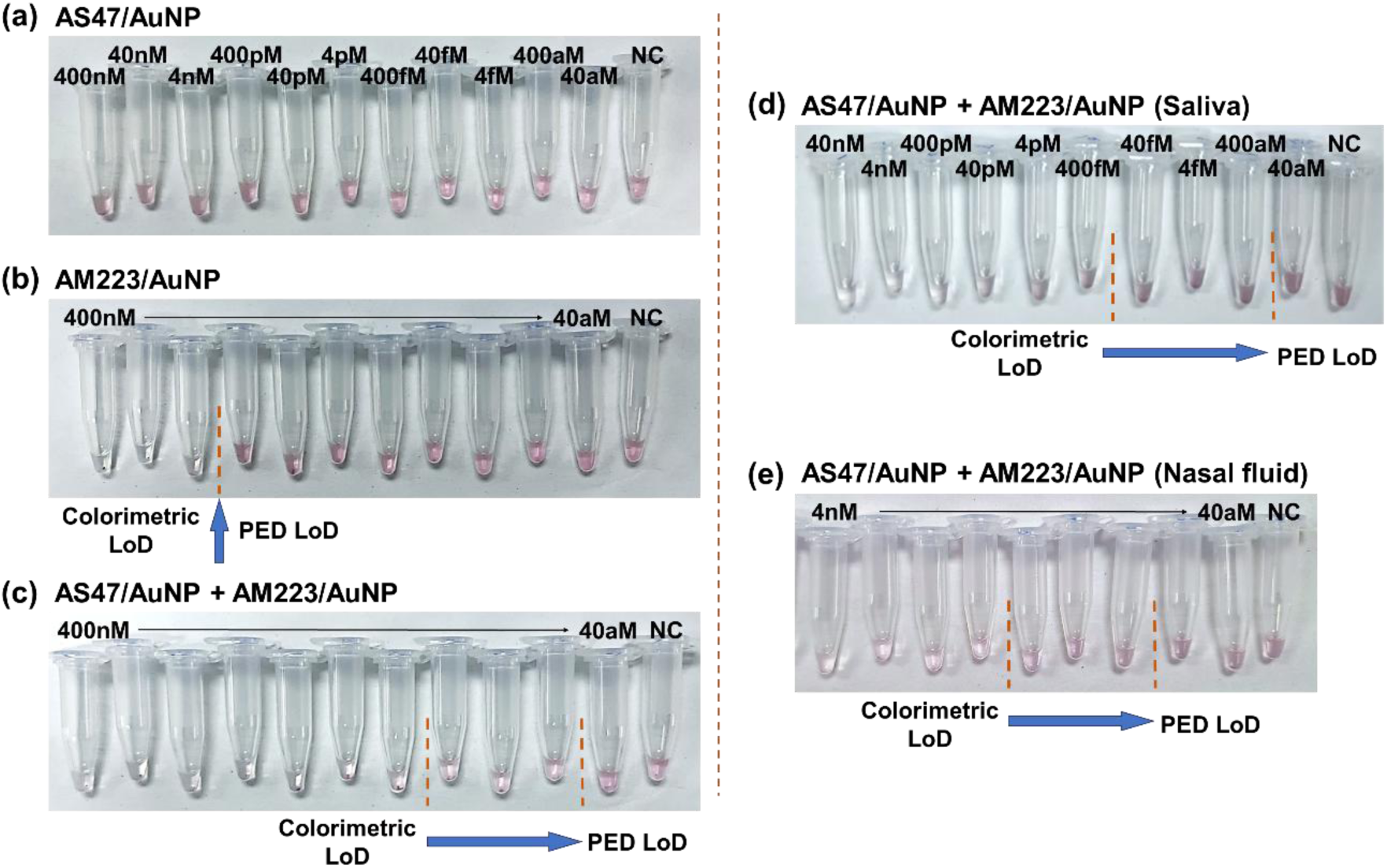
SARS-CoV-2 N-protein antigen detection in biological fluids. (a-c) Optical images showing antigen samples using AuNPs functionalized with different mAbs in PBS buffer: (a) only AS47, (b) only AM223, (c) co-binder (AS47 and AM223) on two sets of AuNPs. The final antigen concentrations ranged from 400 nM to 40 aM. NC: buffer only. (d-e) Optical images showing antigen samples ready for NasRED readout: (d) in human saliva and (e) in human nasal fluid. The samples were centrifuged at 1,200 g for 5 minutes, incubated for 20 minutes, and vortex-stirred at 34.2 rps for 5 seconds.

## 6. Quantitative Polymerase Chain Reaction (qPCR) response settings and conditions

In this work, qPCR was performed by ZeptoMetrix.

In the qPCR protocol for SARS-CoV-2 RNA, the temperature was set to 50°C for 2 minutes for the reverse transcription step to convert the RNA into complementary DNA (cDNA). This was followed by an initial denaturation step at 95°C for 20 seconds to separate the double-stranded cDNA into single strands. Then, during a thermocycling step that repeated for 40 times, the temperature was maintained at 95°C for 3 seconds and then at 60°C for 30 seconds for the annealing and extension step, where primers (Table S4) specific to N-protein coding gene bonded to the cDNA and new DNA strands were synthesized.

To quantify the concentrations of the PROtrol sample, the samples were diluted, tested in triplicate and each repeated in two separate runs. The undiluted sample (nt) had an average cycle threshold (C_T_) value of 12.57, corresponding to a concentration of 1.57 × 10^9^ copies/mL. However, given the high testing concentration, the nt values were only used as a reference but excluded from the calculation of virion particle concentrations to prevent skewing of the data. The 1:10 dilution had an average C_T_ value of 14.68, translating to a concentration of 4.92 × 10^9^ copies/mL for undiluted sample. The 1:100 dilution sample resulted in an average C_T_ value of 17.83, i.e. a concentration of 2.35 × 10^9^copies/mL for the undiluted sample. Further, the 1:1000 dilution yielded an average C_T_ value of 21.22, or a concentration of 5.23 × 10^9^copies/mL for the undiluted sample. The positive extraction control had a C_T_ value of 24.97, while the positive plate control had a C_T_ value of 26.42. The no-template control (water) showed an ‘Undetermined’ C_T_ value, indicating no contamination in the reagents used.

**Figure S15.**
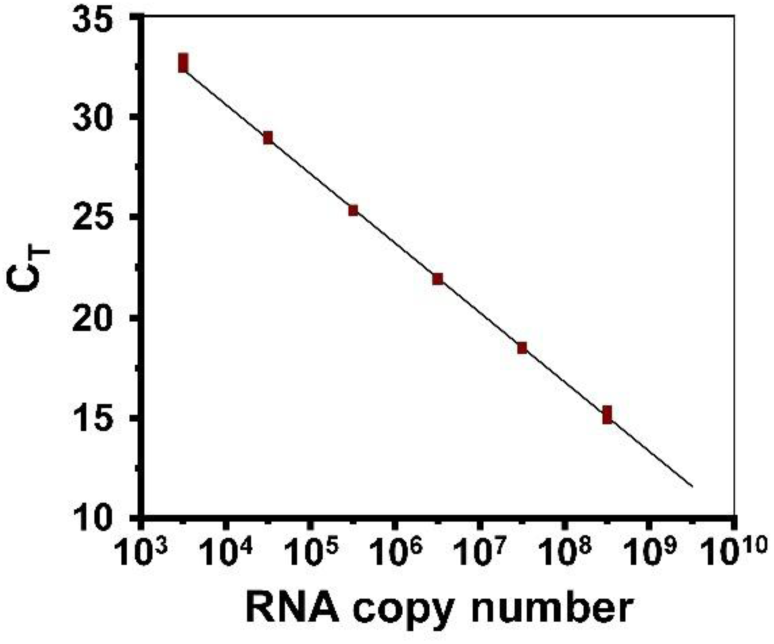
Standard curve of SARS-CoV-2 RNA detection using qPCR. Standard curve of the C_T_ versus the RNA copy number, showing a slope of −3.492, an efficiency of 93.376%, and an R-squared value of 1.00, indicating a highly efficient and reliable qPCR assay.

**Table S4.**
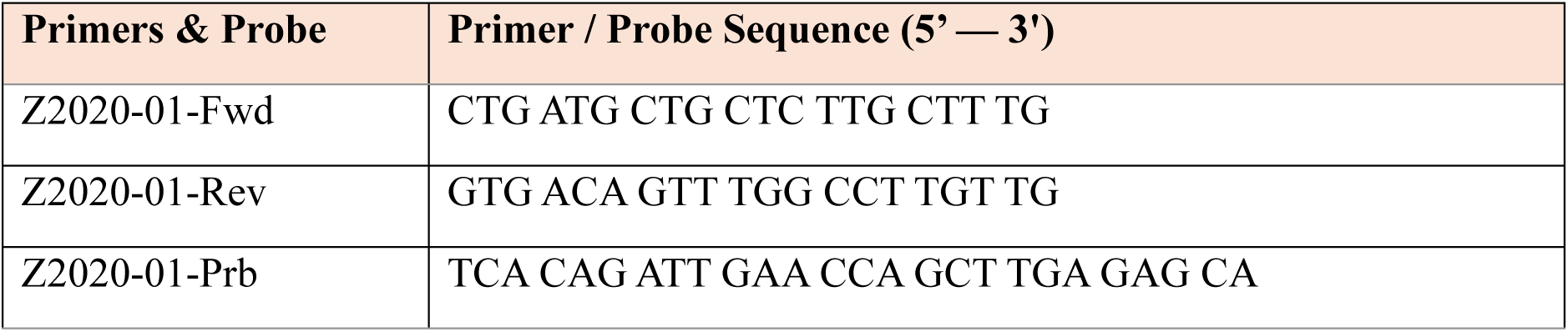
DNA sequence of primer and probe for qPCR reaction.

**Table S5.** Inactive virus sample summary and NaSRED test concentrations.

| Assay /Measured | Initial Concentration | Initial Volume (μl) | Triton X-100 Volume (μl) | Final Volume (μl) | NasRED Test Concentration Before Dilution |
| --- | --- | --- | --- | --- | --- |
| ELISA/N-protein | 60.5 nM | 10 | 10 | 20 | 30.03 nM |
|  | 2995 ng/mL | 10 | 10 | 20 | 1500 ng/mL |
| qPCR | $4.69 \times 10^9$ copies/mL | 10 | 10 | 20 | $2.35 \times 10^9$ copies/mL |
| Culture | $2.73 \times 10^7$ TCID50/mL | 10 | 10 | 20 | $1.365 \times 10^7$ TCID50/mL |

## 7. Comparison of NasRED with representative diagnostic technologies

**Table S6.** Competing diagnostic approaches for SARS-CoV-2 detection: commercial LFAs.

| Test name | LoD (ng/mL) | LoD (TCID<sub>50</sub>/mL) | Detection time (min) | Testing cost | Ref. |
| --- | --- | --- | --- | --- | --- |
| Flowflex COVID-19 Antigen Home Test | 0.43 | $3.57 \times 10^3$ | 15-30 | ~\$8 | 11 |
| Genabio COVID-19 Rapid Self-Test kit | 3 | $2.52 \times 10^4$ | 15 | ~\$12 | 12 |
| iHealth COVID-19 Antigen Rapid Test | 3.374 | $2.83 \times 10^4$ | 15 | ~\$10 | 13 |
| Panbio COVID19 Ag Rapid Test Device | 0.027 | $2.26 \times 10^2$ | 15 | ~\$12 | 14 |
| Fastep COVID-19 Antigen Home Test | 0.047 | $3.93 \times 10^2$ | 15 | ~\$3 | 15 |
| NasRED | 0.200 | $1.7 \times 10^3$ | 15-30 | ~\$2 | This Work |

**Figure S15.**
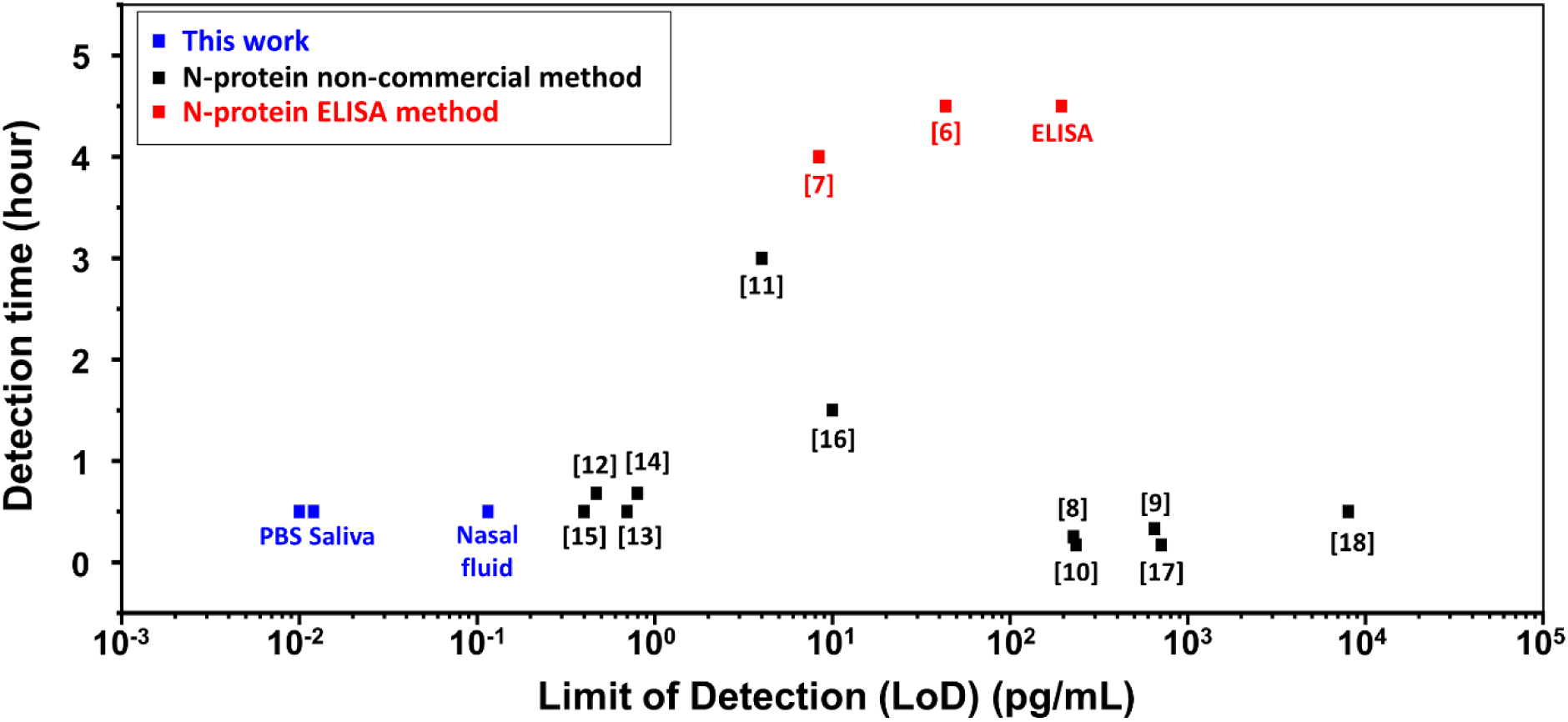
Assay LoD and time comparison. NasRED presented a low LoD and short detection time, both the best among the various methods examined in this study.

**Table S7.**
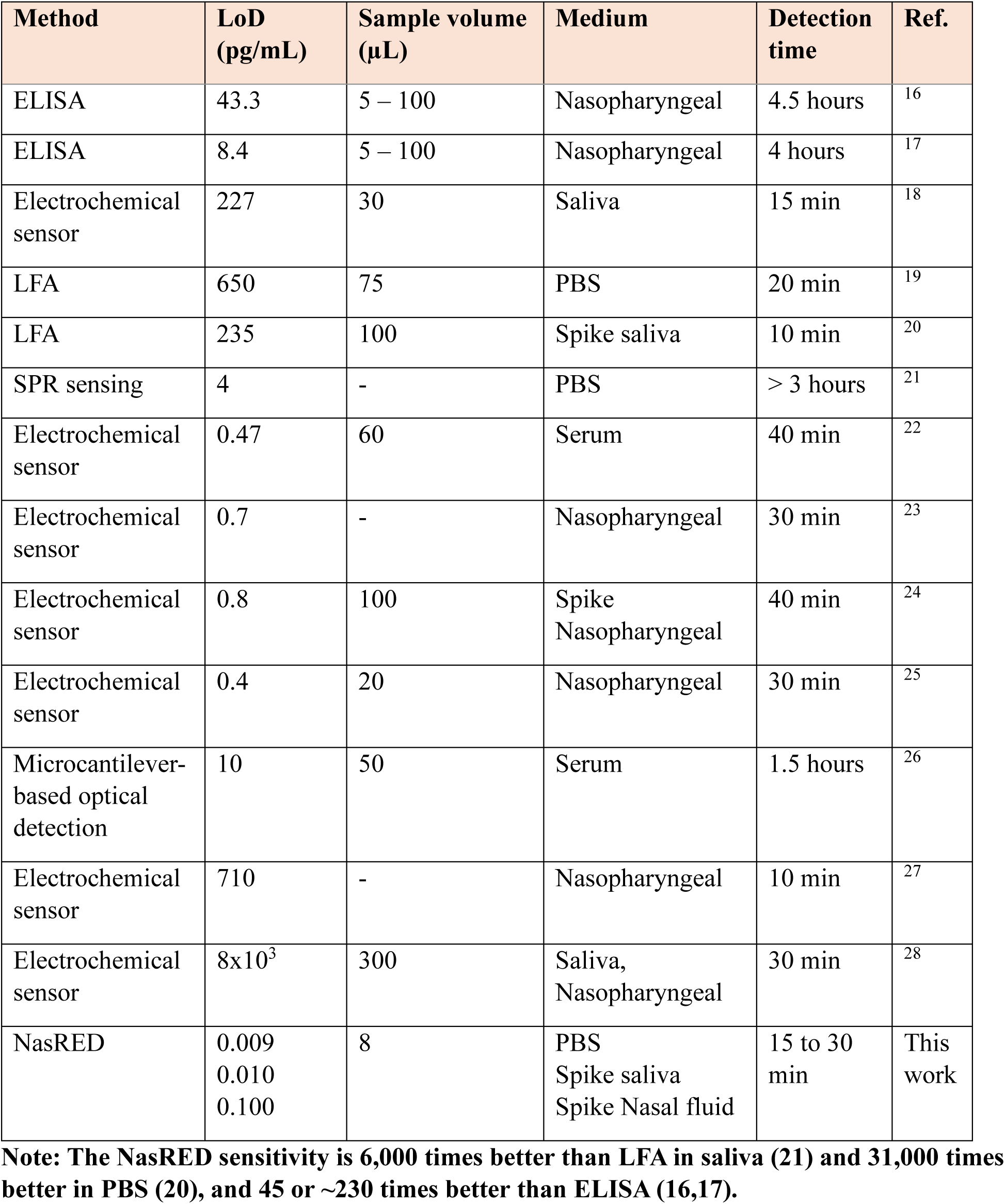
Competing diagnostic approaches for SARS-CoV-2 N-Protein detection: A literature review.

| Method | LoD (pg/mL) | Sample volume (μL) | Medium | Detection time | Ref. |
| --- | --- | --- | --- | --- | --- |
| ELISA | 43.3 | 5 – 100 | Nasopharyngeal | 4.5 hours | <sup>16</sup> |
| ELISA | 8.4 | 5 – 100 | Nasopharyngeal | 4 hours | <sup>17</sup> |
| Electrochemical sensor | 227 | 30 | Saliva | 15 min | <sup>18</sup> |
| LFA | 650 | 75 | PBS | 20 min | <sup>19</sup> |
| LFA | 235 | 100 | Spike saliva | 10 min | <sup>20</sup> |
| SPR sensing | 4 | - | PBS | > 3 hours | <sup>21</sup> |
| Electrochemical sensor | 0.47 | 60 | Serum | 40 min | <sup>22</sup> |
| Electrochemical sensor | 0.7 | - | Nasopharyngeal | 30 min | <sup>23</sup> |
| Electrochemical sensor | 0.8 | 100 | Spike Nasopharyngeal | 40 min | <sup>24</sup> |
| Electrochemical sensor | 0.4 | 20 | Nasopharyngeal | 30 min | <sup>25</sup> |
| Microcantilever-based optical detection | 10 | 50 | Serum | 1.5 hours | <sup>26</sup> |
| Electrochemical sensor | 710 | - | Nasopharyngeal | 10 min | <sup>27</sup> |
| Electrochemical sensor | 8x10<sup>3</sup> | 300 | Saliva, Nasopharyngeal | 30 min | <sup>28</sup> |
| NasRED | 0.009<br>0.010<br>0.100 | 8 | PBS<br>Spike saliva<br>Spike Nasal fluid | 15 to 30 min | This work |

**Note:** The NasRED sensitivity is 6,000 times better than LFA in saliva (21) and 31,000 times better in PBS (20), and 45 or ~230 times better than ELISA (16,17).

**Table S8.** Competing diagnostic approaches for SARS-CoV-2 detection: PCR.

| Manufacturer and instrument | LoD (copies/mL) | Time-to-result | Sample volume (μL) | Size (width x height x depth, cm<sup>3</sup>) and weight (kg) | Instrument Cost | Ref. |
| --- | --- | --- | --- | --- | --- | --- |
| Roche, Cobas 6800 | 250 | 3.5 hours | 400 | 292x222x129, 1624kg | ~\$20,000 | 29 |
| Abbott, M2000 | 100 | 5 hours | 250 | 145x217x79, 314kg | ~\$100,000 | 30 |
| Hologic, Panther | 600 | 2.4 hours | 25 | 193x175x81, 363kg | ~\$80,000 | 31 |
| DiaSorin, Simplexa | 500 | 1.5 hours | 50 | 30x30x20, 30kg | ~\$40,000 | 32 |
| Roche, GenMark ePlex | 10<sup>6</sup> | 1.5 hours | 200 | 54x59x48, 49kg | ~\$20,000 | 33 |
| Roche, LightCycler 480 | 785 | 40 min | 10-50 | 57x50x59, 56kg | ~\$15,000 | 34 |
| Thermo Fisher, QuanStudio™ 5 | 350 | 30 min | 20 | 27x40x50, 26kg | ~\$55,000 | 35 |
| BIO RAD, CFX96 Dx | 250 | 30 min | 10-25 | 33x36x46, 21kg | ~\$12,000 | 36 |
| Thermo fisher, Applied Biosystems 7500 Fast Dx | 250 | 40 min | 10-30 | 34x49x45, 34kg | ~\$12,000 | 36 |
| NasRED | 2.9×10<sup>5</sup>* | 15 to 30 min | 8 | 5.8x5.4x8.5, <1kg ** | ~\$20 | This work |
| | | | | 35x36x25, <15kg *** | ~\$3,000 | |

\*The LoD was found to be 2.9×10<sup>5</sup> copies/mL without amplification.
\*\* The PED reader itself weighs only 125 grams.
\*\*\* The weight included the centrifuge, vortexer, and the PED reader. The centrifuge weighs 10 kg, and the vortex weighs 4.5kg.

## Appendix: Python code for AuNP concentration calculation

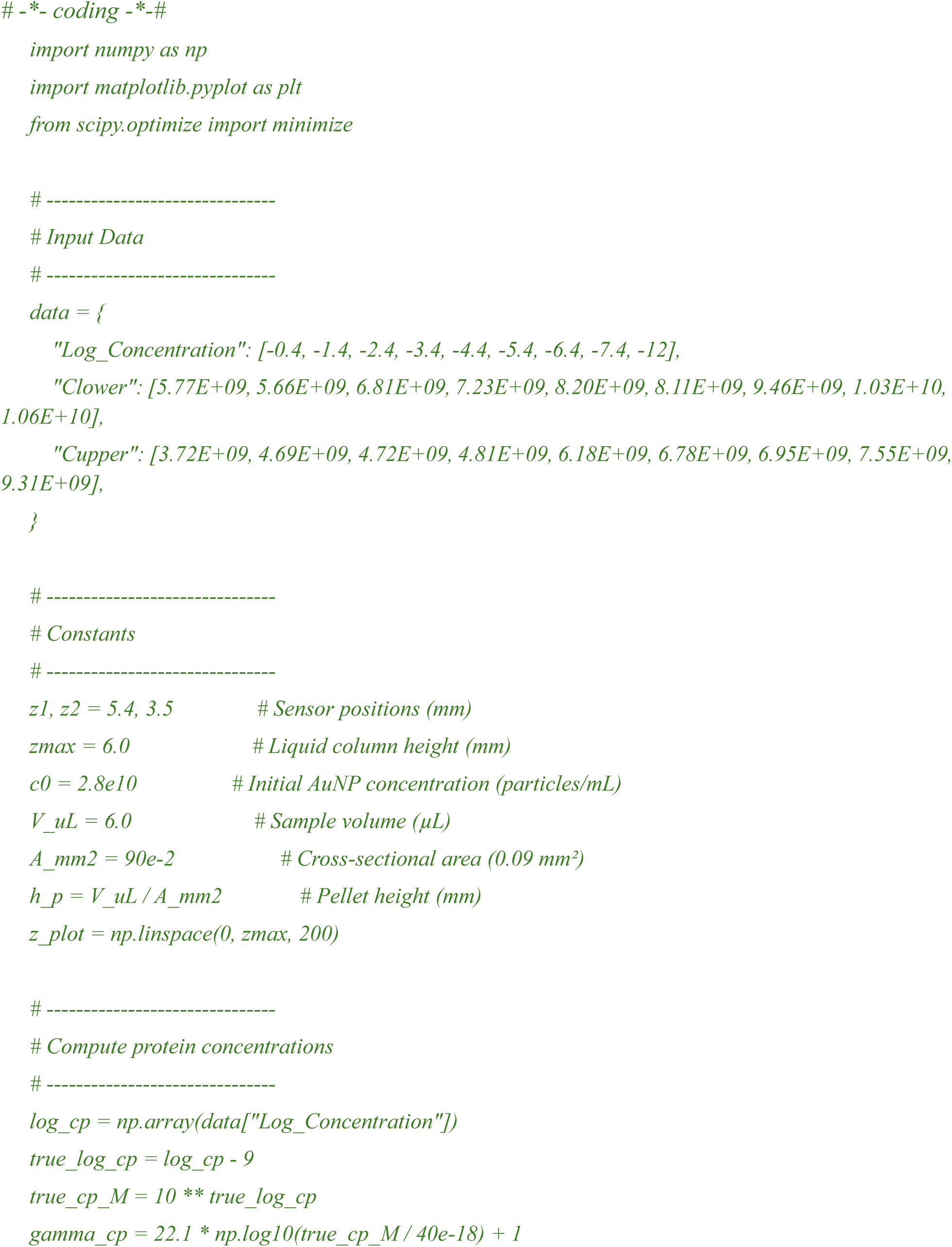
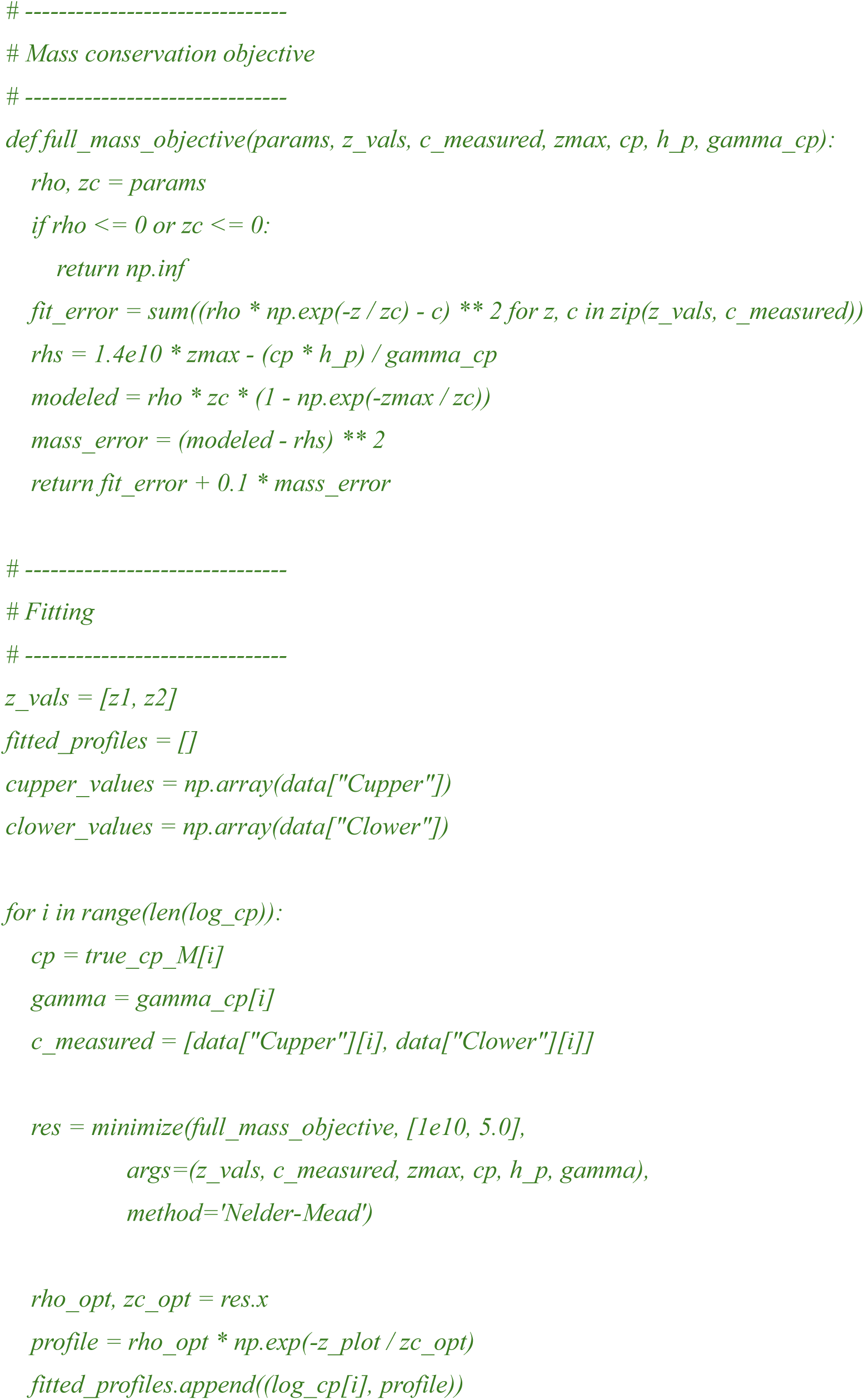
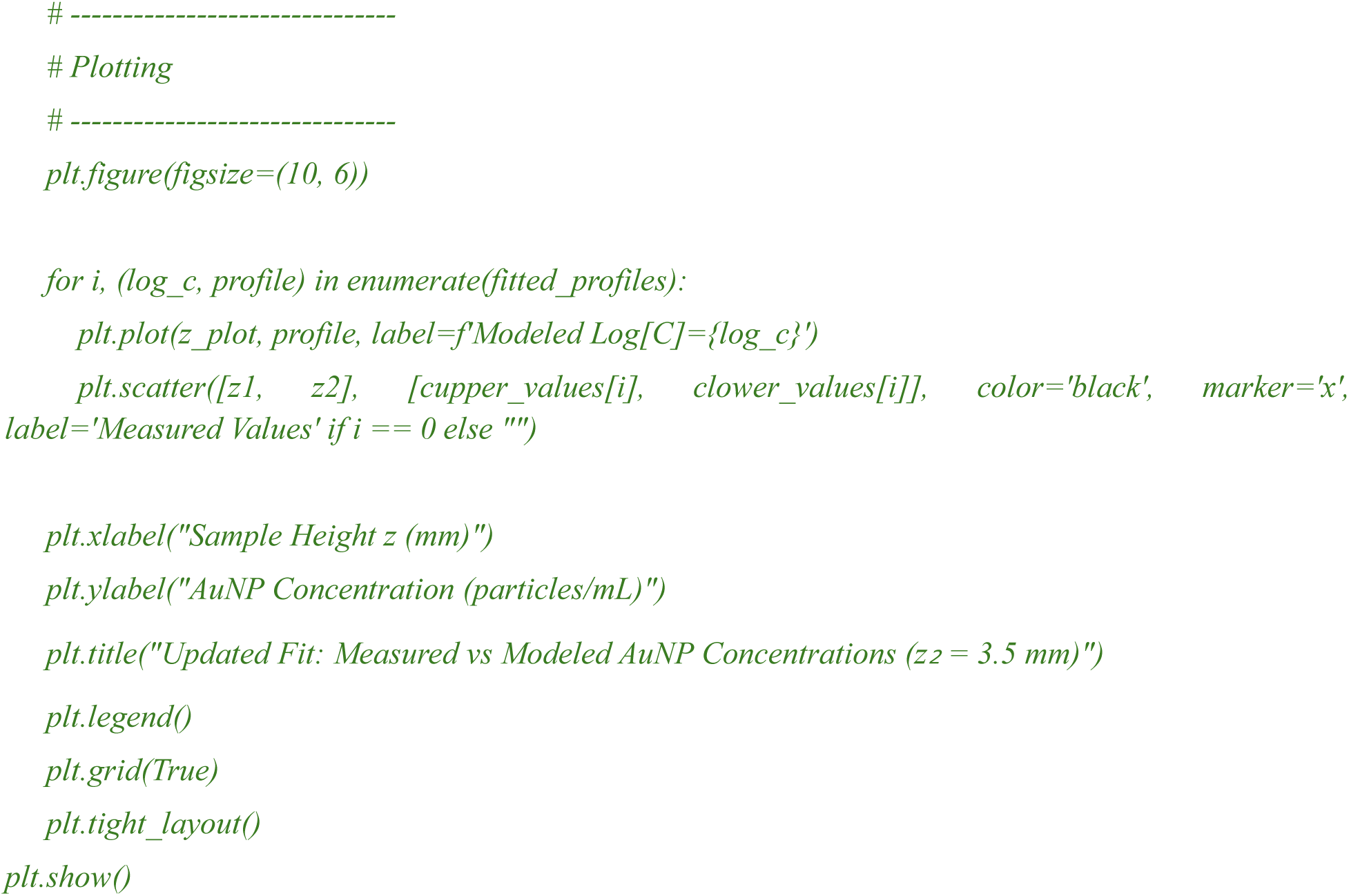

